# The Arabidopsis transcription factor NLP2 regulates early nitrate responses and integrates nitrate assimilation with energy and carbon skeleton supply

**DOI:** 10.1101/2022.03.21.485211

**Authors:** Mickaël Durand, Virginie Brehaut, Gilles Clément, Zsolt Kelemen, Julien Macé, Regina Feil, Garry Duville, Alexandra Launay-Avon, Christine Paysant-Le Roux, John E. Lunn, François Roudier, Anne Krapp

## Abstract

Nitrate signaling improves plant growth under limited nitrate availability and, hence, optimal resource use for crop production. Ongoing work has identified several transcriptional regulators of nitrate signaling, including the *Arabidopsis thaliana* transcription factor NIN-LIKE PROTEIN 7 (NLP7), but additional regulators likely remain to be identified. Here, we characterized Arabidopsis NLP2 as a master upstream transcriptional regulator of early nitrate responses that interacts with NLP7 *in vivo* and shares key molecular features such as nitrate-dependent nuclear localization, a DNA binding motif, and some target genes with NLP7. Additional genetic, genomic and metabolic approaches revealed a specific role for NLP2 in the nitrate-dependent regulation of carbon and energy-related processes that likely influence plant growth under distinct nitrogen environments. Our findings highlight the complementarity and specificity of NLP2 and NLP7 in orchestrating a multi-tiered nitrate regulatory network that links nitrate assimilation with carbon and energy metabolism for efficient nitrogen use and biomass production.

**One-sentence summary:** NLP2 and NLP7 orchestrate plant responses to nitrate supply and control nitrate- dependent regulation of carbon and energy metabolism.

## Introduction

Plants have the crucial ability to sense and promptly acclimate to fluctuating environmental conditions. Plants exhibit incredible phenotypic plasticity, with changes ranging from the transcript level to the whole-organism level, that helps them cope with stresses such as mineral nutrient deficiency (Sultan, 2000). Among these nutrients, availability of the macronutrient nitrogen (N) frequently limits plant growth and development (Marschner, 1995). N availability in the soil fluctuates, reaching sub- or supra-optimal levels and this variation occurs over both temporal and local scales, particularly in agricultural systems where growers periodically apply N fertilizers (Xu et al., 2012).

Roots take up N mainly in the form of nitrate (NO_3_^−^) and ammonium (NH_4_^+^). Nitrate is successively reduced by nitrate reductase (NR) and nitrite reductase (NiR) into ammonium to be finally incorporated into amino acids by the glutamine synthetase/GOGAT cycle (Krapp et al., 2014). Nitrate reduction from nitrate to ammonium is a highly reductant-consuming process usually considered the second largest sink for reducing equivalents after carbon fixation (Noctor and Foyer, 1998). The oxidative pentose phosphate pathway (OPP) is a critical pathway that provides the reductants required for nitrate assimilation (Kruger and von Schaewen, 2003). In addition to energy, incorporation of ammonium into N-containing organic compounds requires carbon skeletons provided by intermediates of the tricarboxylic acid cycle (TCA cycle) and glycolysis, thus highlighting the close link between N and carbon metabolism. N acquisition and interconnected pathways therefore need to be precisely fine-tuned to efficiently cope with transient soil N availability to sustain plant growth while maintaining nutrient homeostasis.

In addition to its role as a major plant nutrient, nitrate itself acts as a signaling molecule influencing the expression of a plethora of genes related to N metabolism as well as carbon metabolism, sulfur and phosphorus assimilation, phytohormone metabolism and responses, and myriad genes encoding transcription factors (TFs) (Wang et al., 2000, 2003; Scheible et al., 2004; Varala et al., 2018). Nitrate triggers the rapid expression alteration of over 1,000 genes in a time- and cell type–dependent manner, referred to as the primary nitrate response (PNR, Medici and Krouk, 2014). The PNR is specific and highly sensitive for nitrate (Wang et al., 2000, 2003) and independent of protein synthesis (Gowri et al., 1992).

Recent advances based on genomics and systems biology approaches (Gaudinier et al., 2018; Medici et al., 2015; Brooks et al., 2019; O’Malley et al., 2016) contributed to establishing a blueprint of a multi-tiered N-related gene regulatory network (GRN) (Vidal et al., 2020). Among the key TFs involved in the nitrate-related GRN in Arabidopsis (*Arabidopsis thaliana*), NIN-LIKE PROTEIN 7 (NLP7), a member of the NIN-Like protein family, stands out as an orchestrator of the nitrate-regulated transcriptional response (Marchive et al., 2013). The Arabidopsis genome encodes nine NLPs that are all thought to contribute to the nitrate response, as they can all bind the nitrate responsive *cis*-element (NRE) *in vitro* (Konishi and Yanagisawa, 2013). Nitrate itself regulates the nuclear accumulation of Arabidopsis NLP7 and NLP6 via a nuclear retention mechanism that leads to the transcriptional activation of early nitrate-responsive genes and involves phosphorylation of a conserved S205 residue by calcium-dependent protein kinases (CDPKs) (Marchive et al., 2013; Guan et al., 2017; Liu et al., 2017). Although essential for nitrate-promoted seed germination, NLP8 function is not regulated by nitrate-dependent nuclear retention, as the protein remains in the nucleus regardless of nitrate application (Yan et al., 2016). More recent work showed that Arabidopsis NLP2, like NLP7, plays a major role in vegetative growth, as both *nlp2* and *nlp7* single mutants grown in the presence of nitrate exhibited reduced rosette biomass (Konishi et al., 2021). Therefore, although they belong to distinct phylogenetic clades (Schauser et al., 2005; Chardin et al., 2014; Mu and Luo, 2019; Liu and Bisseling, 2020), NLP2 and NLP7 share similar properties such as nitrate-dependent activation (Konishi et al., 2021). However, how NLP2 affects nitrate-dependent growth remains elusive, as does its relative position and influence within the nitrate-related GRN.

To understand the role of NLP2, we identified the loci bound by NLP2 and the perturbations of the transcriptome associated with loss of NLP2 function in response to nitrate. These analyses revealed the role of NLP2 in orchestrating the early response to nitrate and part of its downstream GRN, which includes feed-forward loops that enable rapid and strong responses to nitrate availability. In parallel, we assessed the changes in biomass and metabolism in *nlp2* mutants to decipher how NLP2 governs nitrate-dependent growth, which unraveled specific and overlapping functions with NLP7. Thus, we discovered a major role for NLP2 in regulating carbon and energy metabolism in response to nitrate. Our findings uncover the influential role of NLP2 in nitrate-dependent orchestration of N metabolism and associated processes that rely in part on the molecular interplay between NLP2 and NLP7.

## Results

### NLP2 is retained in the nucleus in a nitrate-dependent manner

Since several, but not all, NLP TFs are regulated via nitrate-triggered nuclear accumulation, we tested if the intracellular localization of NLP2 was regulated via a rapid nuclear retention mechanism in response to nitrate in roots. Accordingly, we introduced a construct encoding a translational fusion between NLP2 and the green fluorescent protein (GFP) and driven by the *NLP2* promoter (*NLP2pro:NLP2-GFP*) in the *nlp2-2* mutant background. This construct fully complemented the *nlp2-2* phenotype and exhibited relative *NLP2-GFP* transcript levels similar to those of endogenous *NLP2* in the wild-type (WT) Col-4 (Supplemental Figure S1). We inspected GFP fluorescence in the root tips of N-depleted seedlings before and after nitrate resupply. In seedlings grown under N starvation conditions, we detected NLP2- GFP exclusively in the cytoplasm, while a 10-min nitrate addition led to the

relocalization of NLP2-GFP to the nucleus (Figure 1, Supplemental Movie 1). In the elongation zone, we observed NLP2-GFP in the nucleus of stele cells and to a lesser extent in the nucleus of cortex cells (Supplemental Figure S2, A, B and C). This nuclear accumulation was specifically triggered by nitrate and did not occur after supplying other N sources such as ammonium (Supplemental Figure S2, D and E). NLP2-GFP also was relocalized in the nucleus in response to nitrate in seedlings overexpressing *NLP2-GFP* using the constitutive *UBIQUITIN10* (*UBQ10*) promoter (Supplemental Figure S2, G and H) suggesting that transcriptional regulation is not part of the NLP2 nuclear accumulation mechanism.

**Figure 1.**
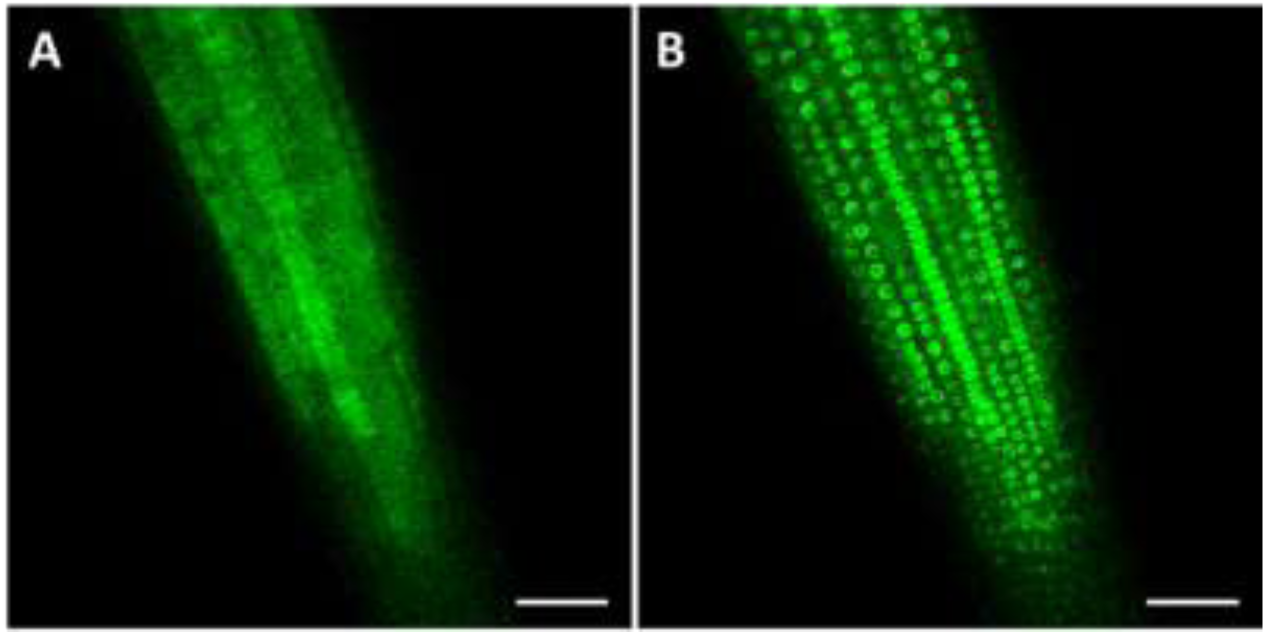
NLP2 rapidly accumulates in the nucleus after nitrate resupply to N-starved seedlings in the root tip. Confocal imaging was performed on *nlp2-2* seedlings harboring the *NLP2pro:NLP2-GFP* transgene. **(A)** Root tip surface of N-starved seedlings. **(B)** Root tip surface of N-starved seedlings after 10 min of nitrate resupply. Scale bar = 50 µm.

In addition, this nitrate-dependent control of NLP2 subcellular localization was effectively mimicked by treatment with the nuclear export inhibitor leptomycin B (Supplemental Figure S2F). Thus, NLP2 accumulates rapidly in the nucleus in response to nitrate, likely governed by a post-translational mechanism based on nuclear retention, as previously shown for NLP7 and NLP6 (Marchive et al., 2013; Liu et al., 2017; Guan et al., 2017).

### NLP2 binds hundreds of genes related to transcriptional response to nitrate

The nitrate-specific regulation of NLP2 nuclear accumulation suggested a prominent upstream regulatory role for this TF in the nitrate transcriptional response, which prompted us to identify the genomic regions bound by NLP2 *in vivo*. To this end, N- starved seedlings harboring the *NLP2pro:NLP2-GFP* transgene in the *nlp2-2* mutant background were resupplied with nitrate for 30 min before crosslinking and chromatin immunoprecipitation (ChIP) using an anti-GFP antibody. ChIP-seq analysis identified a robust set of 462 regions bound by NLP2 in response to nitrate addition that correspond to 550 annotated genes (Supplemental Table S1). Representative examples of NLP2 target loci are presented in Supplemental Figure S3.

Peak-summit mapping over bound loci showed prominent (≈60%) binding for NLP2 toward the transcription start site of target genes, with a smaller proportion (≈30%) of binding sites near the transcription termination site (TTS) (Figure 2A). We applied the MEME suite (Bailey et al., 2015) to search for overrepresented DNA motifs in 200-bp fragments centered on NLP2 peak summits and identified a consensus DNA sequence (TGNCYYTT) in all 462 regions bound by NLP2 (Figure 2B). This consensus sequence displayed a striking similarity to the NLP7 binding motif identified *in vitro* by DNA affinity purification sequencing (DAP-seq) (O’Malley et al., 2016). Furthermore, this motif was also highly similar to the 5′ half of the palindromic nitrate-responsive element (NRE, Konishi and Yanagisawa, 2010) that is bound by all NLP proteins *in vitro* (Konishi and Yanagisawa, 2013). Looking for complete palindromic NREs we identified 11 and 89 NLP2-bound regions with either full (AARRGNCA) or partial (AARRG) 3′ fragments of the NRE, respectively located at 8 to 12 bp 3’ of the consensus DNA sequence found in all NLP2-bound regions (Supplemental Table S1). Thus, a substantial fraction of NLP2 binding sites correspond to an NRE-like sequence, suggesting a tight relationship between NLP2 and the transcriptional response to nitrate.

**Figure 2.**
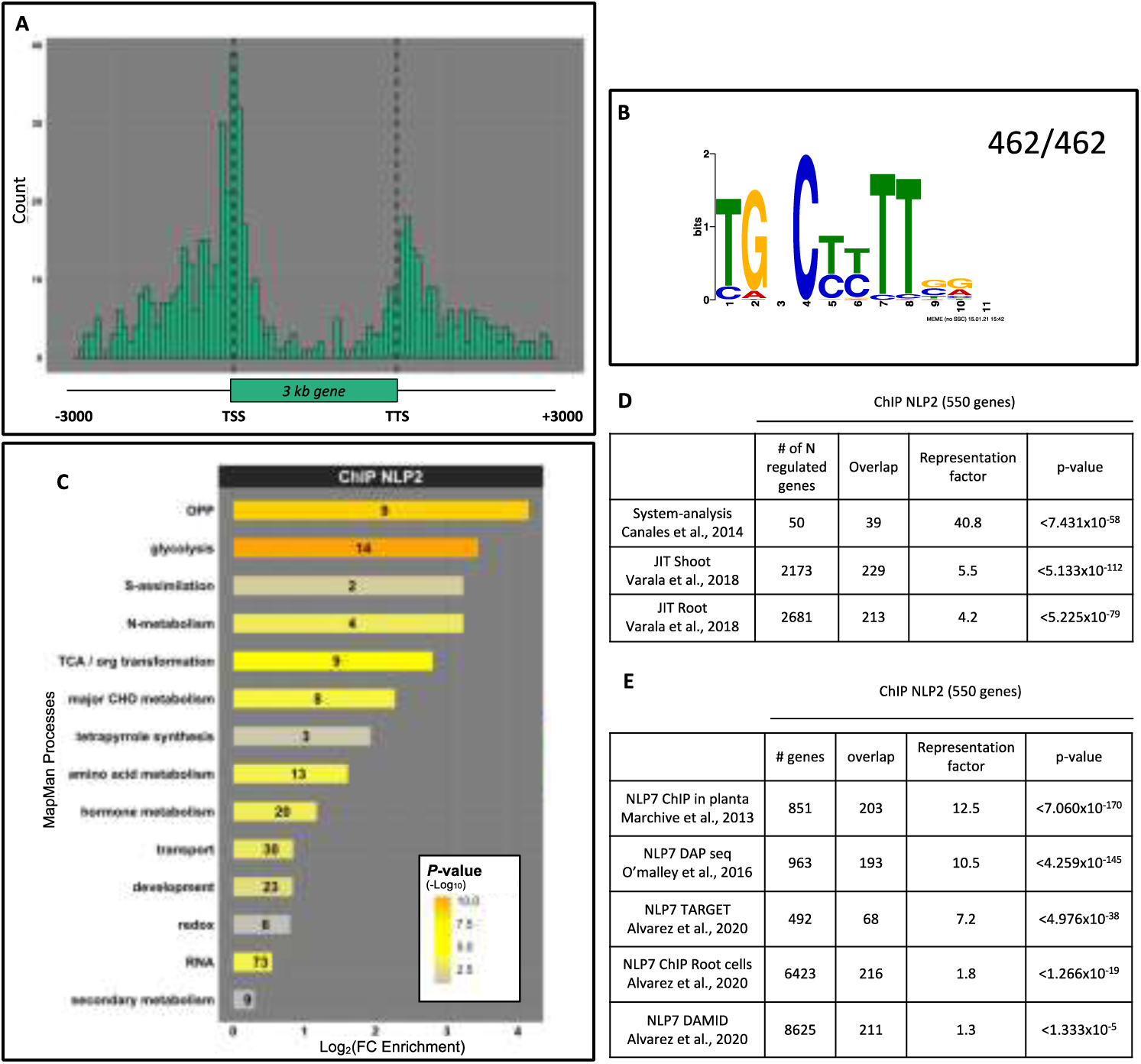
NLP2 binds an NRE-like *cis*-element highly associated with genes related to the transcriptional response to nitrate. Seedlings carrying the *NLP2pro:NLP2-GFP* transgene were grown in N- replete (3mM nitrate) liquid culture for 11 days, followed by N depletion for 3 days; 3 mM nitrate was then added for 30 min. ChIP was conducted using an antibody against GFP. **(A)** Distribution of NLP2 binding sites along the gene body of targets. TSS, transcription start site; TTS, transcription termination site. To account for different gene sizes, the positions of binding sites within transcribed regions were normalized to gene length. **(B)** Consensus DNA motif enriched in NLP2-bound DNA sequences. **(C)** Major Gene Ontology (GO) term enrichment analysis for NLP2-bound genes. GO terms were defined using MapMan bins and significant enriched GO terms. The number of genes is indicated in each bar; the supporting *P*-value is shown as a color scale. Significance is represented as a gradual color scale from grey to orange. **(D)** Overrepresentation of NLP2-bound genes among nitrate-regulated genes. **(E)** Overlap of NLP2- and NLP7-bound genes with published data for NLP7.

Gene Ontology (GO) analysis of the NLP2-bound genes highlighted a significant enrichment (p < 0.05) in N-metabolism and connected metabolic pathways such as the OPP, amino acid metabolism, sulfur assimilation, and carbon metabolism pathways (Figure 2C). Indeed, a substantial proportion of NLP2 target genes were among the top 50 nitrate-responsive genes identified in a metastudy (Canales et al., 2014) and among the genes regulated by N in a temporal manner (Varala et al., 2018) (Figure 2D). In addition, NLP2-bound genes showed a significant enrichment in GO terms related to phytohormone metabolism, transport, and transcriptional regulation (Figure 2C). The NLP2-bound genes contributing to this last GO term included several genes encoding TFs that are involved in the transcriptional response to nitrate availability such as HRS1 HOMOLOG1 (HHO1), HHO2, HHO3 (Medici et al., 2015; Maeda et al., 2018; Kiba et al., 2018; Safi et al., 2021), LOB DOMAIN-CONTAINING PROTEIN37, 38 and 39 (Rubin et al., 2009), TGACG SEQUENCE-SPECIFIC BINDING PROTEIN 1 and 4 (Alvarez et al., 2014), and ABSCISIC ACID RESPONSIVE ELEMENTS-BINDING FACTOR 2 (ABF2) (Contreras-López et al., 2022).

In agreement with the similarity between the NLP2 and NLP7 binding motifs, we detected a significant overlap between NLP2-bound genes and previously identified repertoires of NLP7 targets. We obtained the most significant overlaps with datasets for NLP7 regions bound *in vivo* (Marchive et al., 2013) and *in vitro* (O’Malley et al., 2016) as well as in root protoplasts (Alvarez et al., 2020) (Figure 2E, Supplemental Table S1). Taken together, these observations confirm that NLP2 likely has a prominent role in the nitrate response GRN.

### NLP2 binding is mainly associated with nitrate-induced gene expression

To unravel the relationship between NLP2 binding to its target loci and the transcriptional response to nitrate, we isolated two T-DNA insertional mutants in *NLP2*, the knockout mutant lines *nlp2-2* and *nlp2-3* (Supplemental Figure S4), and performed a transcriptome deep sequencing (RNA-seq) analysis of the *nlp2-2* mutant compared to WT seedlings after resupplying nitrate for 30 min (labeled as WT30 or *nlp2-2* 30) after 3 days of N depletion (labeled as WT0 or *nlp2-2* 0, corresponding to the no added N control). We identified 2,948 differentially expressed genes (DEG) in response to nitrate in the WT (corrected *p*-value ≤ 0.05; Log_2_ [WT30/WT0] fold-change ≥ 0.6 or ≤ –0.6).

We grouped these DEGs into four classes depending on the variation of steady state mRNA levels after nitrate addition. Groups 1 and 2 consisted of genes with strongly increased and decreased transcript levels, respectively (N++, n = 953; N--, n = 356; Log_2_ [WT30/WT0] fold-change ≥ or ≤ –1), while groups 3 and 4 comprised genes with weakly increased or decreased transcript levels, respectively (N+, n = 762; N−, n = 877, 0.6 ≤ Log_2_ [WT30/WT0] fold-change ≤ 1, or –0.6 ≥Log_2_ [WT30/WT0] fold-change ≥ –1).

We determined that 10% of the DEGs detected in the WT in response to N resupply are bound by NLP2 in response to nitrate, with 187 (20%), 76 (10%), 3 (0.3%), and 3 (1%) genes belonging to the N++, N+, N−, and N-- DEG classes, respectively. Importantly, nearly 50% of the 550 NLP2-bound genes identified above by ChIP-seq displayed differential expression after 30 min of nitrate resupply, with over 95% showing increased steady state mRNA levels after 30 min of nitrate resupply. These data indicate that NLP2 binding is preferentially associated with the transcriptional activation of genes in response to nitrate.

### The transcriptome-level changes in the early response to nitrate are impaired in *nlp2* mutants

We also compared transcript levels between *nlp2-2* and the WT after nitrate treatment. Among the 2,948 nitrate-responsive genes detected in the WT, 474 genes were differentially expressed in *nlp2-2* (corrected *p*-value ≤ 0.05, Log_2_ [*nlp2-2* 30 / WT30] fold-change ≤ –0.3 or ≥ 0.3, Supplemental Tables S2 and S3), with 70% (335) showing lower steady state mRNA levels in *nlp2-2* compared to the WT. Most of these genes (289/335) were upregulated in response to nitrate in WT (207 N++ and 82 N+), with only 46 showing downregulation in WT (16 N-- and 30 N−) (Figure 3A).

**Figure 3.**
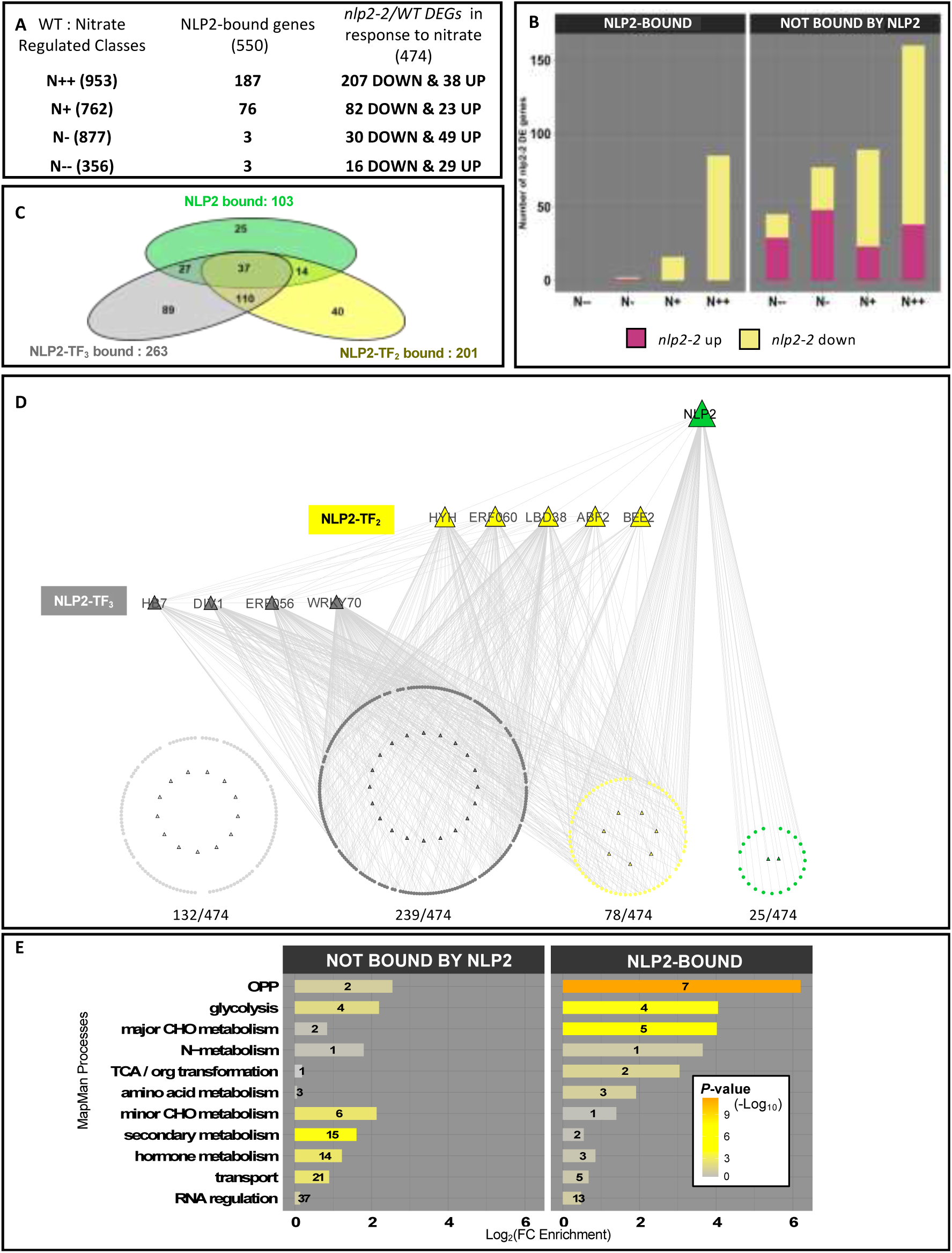
NLP2 orchestrates early nitrate response. Seedlings were cultivated and treated as in Figure 2. **(A)** Overlap between NLP2-bound genes in WT and nitrate-regulated genes, identified as differentially expressed genes (DEGs) between *nlp2-2* and the WT. Nitrate-regulated genes in the WT were selected according to Log2 fold-changes between nitrate treatments at 0 and 30 min with false discovery rate adjustment (*P*-value ≤ 0.05). N++, N-- are DEGs with Log2 (WT30/WT0) fold-change ≥ 1 or ≤ –1, respectively. N+, N− are DEGs with Log2 (WT30/WT0) fold-change ≤ 1 and ≥ 0.6, Log2 (WT30/WT0) fold-change ≤ –0.6 and ≥ –1, respectively. *nlp2-2*/WT DEGs were selected based on Log2 fold-change (Log2FC ≥ 0.3 or Log2FC ≤ -0.3) and false discovery rate adjustment (*P*-value ≤ 0.05) after 30 min of nitrate treatment. **(B)** Number of *nlp2-2*/WT DEGs as a function of nitrate-regulated classes and NLP2-binding status in the WT. **(C)** Overlap of genes bound by NLP2 and genes predicted to be bound by NLP2-TF2s and/or NLP2-TF3s among the *nlp2-2*/WT DEGs in response to nitrate resupply (30 min). NLP2-bound genes are from this study; NLP2-TF2- and -TF3-bound genes were retrieved from ConnecTF. **(D)** NLP2-dependent nitrate gene regulatory network. NLP2 and NLP2-TF2/TF3 binding explains 72% of the transcriptional alterations to nitrate in *nlp2-2*. The network was constructed using Cytoscape (Shannon et al., 2003). Triangles, TFs; circles, other genes. Green, *nlp2-2*/WT DEGs bound by NLP2 only; yellow, *nlp2-2*/WT DEGs bound by NLP2 and other NLP2-TF2/TF3; gray, *nlp2-2*/WT DEGs bound by NLP2-TF2/TF3; light gray, *nlp2-2*/WT DEGs not bound by NLP2 or NLP2-TF2/TF3. **(E)** Enrichment in major MapMan GO terms related to *nlp2-2*/WT nitrate-responsive DEGs, bound or not by NLP2. Significance is represented as a gradual color scale from grey to orange.

To confirm that the observed changes in gene expression were due to the loss of NLP2 function, we measured the relative transcript levels of several DEGs in *nlp2- 2* and the other *nlp2* mutant *nlp2-3* by reverse-transcription quantitative PCR (RT-qPCR) and obtained similar results (Supplemental Figure S5A). In addition, complementation of *nlp2-2* with the introduction of the *NLP2pro:NLP2-GFP* transgene also restored the nitrate regulation of all studied genes, as seen by RT-qPCR (Supplemental Figure S5B). Taken together, these results show that the primary nitrate response is impaired in *nlp2* mutants and that the NLP2 loss of function is mainly associated with a weaker induction of nitrate-regulated genes, suggesting that NLP2 is directly or indirectly related to the transcriptional activation by nitrate.

### NLP2 directly regulates gene expression in response to nitrate

To distinguish between direct transcriptional regulation by NLP2 from indirect effects on the primary nitrate response, we focused on the 474 nitrate-regulated genes showing altered transcript levels after 30 min of nitrate resupply in the *nlp2-2* mutant compared to the WT. Of those DEGs, about 20% (103/474) were bound by NLP2 and almost 100% corresponded to genes strongly induced by nitrate in the WT (101/103; 85 N++ and 16 N+, Figure 3B). All genes showed decreased transcript levels in the *nlp2-2* mutant after nitrate addition in comparison to WT, further substantiating the conclusion that NLP2 is a positive and direct regulator of the PNR.

### NLP2 also contributes to orchestrating the primary nitrate response by directly regulating the expression of downstream TF genes

A sizable fraction of the nitrate-responsive genes differentially expressed in *nlp2-2* relative to the WT but not bound by NLP2 exhibited either increased (249) or decreased (122) transcript levels and were distributed between the different classes of nitrate-regulated genes in the WT (N--, N−, N+, and N++), with an overrepresentation of strong nitrate-regulated genes (160 N++) (Figure 3B). While the absence of NLP2 binding might be due to experimental limitations or transient DNA binding that could not be captured by ChIP assays, we hypothesized that these DEGs are under the control of TFs, and the expression of the genes encoding these TFs in response to nitrate directly depends on NLP2. Indeed, genes encoding several highly influential TFs in the N regulatory network (HHO1, HRS1, LBD38, ETHYLENE RESPONSE FACTOR 056 [ERF056], ERF060, HY5 HOMOLOG [HYH], DIVARICATA1 [DIV1], BR-ENHANCED EXPRESSION2 [BEE2], and ABF2) (Canales et al., 2014; Vidal et al., 2020; Alvarez et al., 2020) were among the *nlp2-2*/WT DEGs and were bound by NLP2.

Using the ConnecTF database (Brooks et al., 2019), we retrieved the putative target genes for five of these ‘secondary’ TFs (hereafter referred to as NLP2-TF_2_: HYH, ERF060, BEE2, LBD38, and ABF2) and determined that 201 *nlp2-2*/WT DEGs are indeed NLP2-TF_2_ target genes (Figure 3C and D; Supplemental Table S3). Thus, over 40% of the transcriptional alterations resulting from NLP2 loss of function might be explained by the action of one or more NLP2-TF_2_ members. We extended this analysis to identify a potential third tier of the NLP2-dependent nitrate regulatory network by looking for TF genes that are bound by NLP2-TF_2_s (hereafter called NLP2-TF_3_) and belonging to the *nlp2-2*/WT DEGs in response to nitrate. We identified 14 TF genes fitting these criteria and obtained putative target genes for four of them (DIV1, ERF056, HOMEOBOX7 [HB7], and WRKY70). These four NLP2-TF_3_s appeared to bind to 263 out of the 474 *nlp2-2*/WT DEGs, of which 199 were not directly bound by NLP2. Thus, by summing up the genes bound by NLP2, NLP2-TF_2_s, and NLP2-TF_3_s, we retrieved almost 75% (342/474) of the *nlp2-2/*WT DEGs and linked their dysregulated transcript levels directly or indirectly to NLP2 (Figure 3C and D; Supplemental Table S3). In addition, 78 of the 103 *nlp2-2/*WT DEGs in response to nitrate that are bound by NLP2 were predicted to be bound also by one or more NLP2-TF_2_ and/or NLP2-TF_3_, suggesting that NLP2-regulated TFs may cooperate with NLP2 to regulate the initial wave of the primary nitrate response. Thus, NLP2 contributes largely to the orchestration of the primary nitrate response by targeting and regulating not only many N metabolism–related genes but also highly influential TF genes in the N regulatory networks.

### NLP2 regulates genes encoding catalytic proteins belonging to N metabolism and associated processes

We observed that the most enriched GO terms among NLP2-bound genes that are also differentially expressed in *nlp2-2* in response to nitrate are directly related to N metabolism or closely associated processes (Figure 3E). This observation was particularly remarkable for the OPP, which provides reductants for nitrate assimilation, as seven of the nine genes showing downregulation in response to nitrate in *nlp2-2* were direct NLP2 targets. We also detected an enrichment for NLP2 target genes among genes related to nitrate transport, reduction, and assimilation, as well as amino acid metabolism. In addition, we also identified genes directly regulated by NLP2 with roles in processes associated with central carbon metabolism such as major CHO metabolism, the TCA cycle, and glycolysis (Figure 3E). Moreover, genes belonging to these GO terms were also the targets of NLP2-TF_2_ and/or NLP2-TF_3_ proteins mentioned above. In addition, targets of NLP2-TF_2/3_ were enriched in processes such as secondary metabolism, hormone metabolism, minor carbon metabolism and transport. Thus, NLP2 also contributes to orchestrating the early nitrate response by directly regulating the expression of downstream TFs to fine-tune genes involved in N metabolism and associated processes.

### NLP2- and NLP7-dependent biomass production under non-limiting N supply shows distinct association with nitrate homeostasis

To assess the physiological role of the NLP2-dependent nitrate regulatory network, we analyzed the growth and metabolic phenotypes of *NLP2* loss-of-function mutants (*nlp2-2* and *nlp2-3*) under contrasting nitrate supply. As we identified a partially overlapping direct target repertoire between NLP2 and NLP7 and a similar DNA binding motif, we included the *nlp7-1* mutant in the analysis. Indeed, *nlp7* mutants display a nitrate-dependent growth delay and accumulate nitrate (Castaings et al., 2009).

To directly compare the consequences of the loss of function of either gene, we grew *nlp2* mutants and the *nlp7-1* mutant in sand culture under short-day conditions (8 h light/16 h dark) with either 0.5 mM nitrate or 5 mM nitrate supply for 42 days and determined rosette and root biomass, as well as nitrate contents (Figure 4; Supplemental Figure S6, A and B). Under growth-limiting N supply (0.5 mM nitrate), the rosette biomass of *nlp2* and *nlp7* mutants was only slightly lower compared to that of the WT (by 25% and 15%, respectively), whereas root biomass decreased by 35% and 50%, respectively. Under non-limiting N supply, WT plants displayed the expected strong (2.1-fold) increase in rosette fresh weight compared to WT plants grown under 0.5 mM N supply. Under the same non-limiting N supply conditions (5 mM nitrate), *nlp2* mutants exhibited a 50% lower rosette biomass and 30% lower root biomass compared to the WT (Figure 4, A, B and C; Supplemental Figure S6, A and B). Rosette biomass of *nlp2* mutants was increased by 34% for *nlp2-3* and 6% for *nlp2-2* compared to the rosette biomass under limiting N supply (0.5mM nitrate). This indicates that *nlp2* mutants could not produce additional biomass when nitrate supply was increased in contrary to WT. This growth phenotype was similar to that of the *nlp7-1* mutant grown under the same conditions, as reported previously (Castaings et al., 2009).

**Figure 4.**
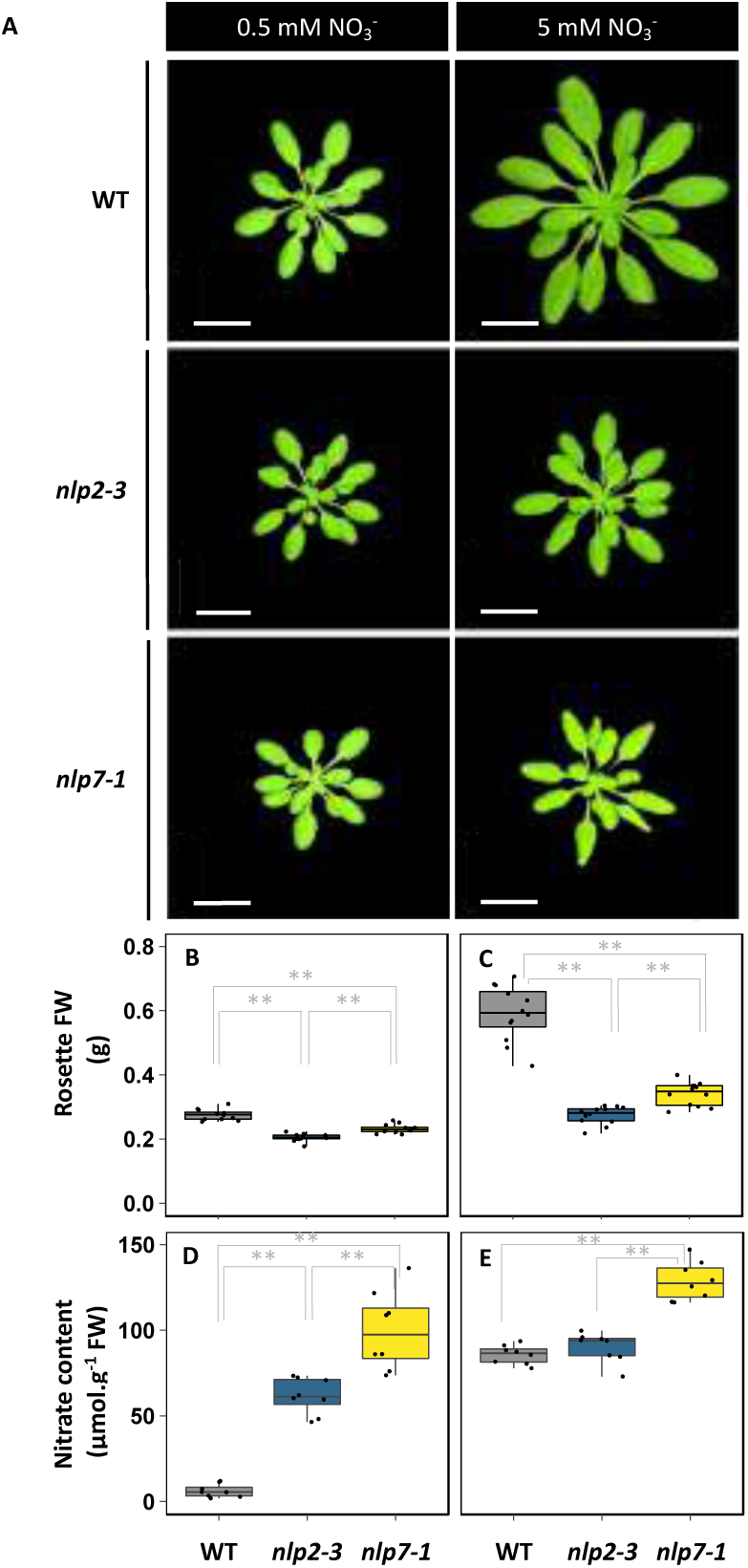
Biomass and nitrate contents of *nlp2-3* and *nlp7-1* mutants grown under limiting and non-limiting nitrate supply. Plants were grown on sand under short days with a light intensity of 150 µE. **(A)** Top view of 42-day-old rosettes of WT, *nlp2-3*, and *nlp7-1* plants grown under limiting (0.5 mM) and non-limiting (5 mM) nitrate supply. Scale bar = 2 cm. (**B, C**) Rosette fresh weight of plants grown under limiting (0.5 mM, **B**) and non-limiting (5 mM, **C**) nitrate supply. (**D, E**) Rosette nitrate contents of WT, *nlp2-3*, and *nlp7-1* plants grown under limiting (0.5 mM, **D**) and non-limiting (5 mM, **E**) nitrate supply. For B-E, data are presented as boxplots with each value shown as a black circle. For B and C, data were obtained from six plants per experiment from two independent experiments. For D and E, data were obtained from four plants per experiment and two independent experiments. Significant differences between genotypes for each nitrate supply condition were determined by a pairwise permutation test followed by false discovery rate adjustment (* *P*-value ≤ 0.05, ** *P*-value ≤ 0.001).

We then measured nitrate contents of rosettes and roots to assess the link between N metabolism and impaired biomass production in *nlp2* mutants (Figure 4, D and F, Supplemental Figure S6, C and D). Under N-limiting conditions, both *nlp2* and *nlp7* mutants accumulated 10-fold and 15-fold more nitrate, respectively, in their rosettes despite the minor difference observed for rosette biomass relative to the WT. Root nitrate contents also showed a similar increase in the *nlp2* and *nlp7* mutants compared to the WT. However, under non-limiting nitrate supply, we observed no differences in nitrate contents between *nlp2* mutants and the WT, while the *nlp7* mutant accumulated 1.5-fold more nitrate in its rosettes compared to the WT (Figure 4E).

We also measured nitrate reductase (NR) activity in the rosettes of plants grown under non-limiting nitrate supply (Supplemental Figure S7). NR activity reached WT levels in *nlp2* mutants and decreased by about 25% in the rosettes of the *nlp7-1* mutant, but this difference appeared too small to explain the difference in nitrate accumulation between the *nlp2* and *nlp7* mutants when taking into account the phenotypes of *nr* mutants (Wilkinson and Crawford, 1993). To confirm that the T-DNA insertion at *NLP2* caused the observed phenotypes, we determined biomass production and nitrate contents in *nlp2-2* mutant lines harboring the *NLP2pro:NLP2- GFP* construct, as shown in Supplemental Figure S1. The complemented line *nlp2-2 NLP2pro:NLP2-GFP* #2.1 restored biomass and nitrate contents to WT levels in its root and rosettes when grown under both nitrate supply conditions, confirming the specificity of the above phenotypes to the loss of NLP2 function (Supplemental Figure S8).

Taken together, these observations suggest that, despite the similar biomass and nitrate accumulation under limiting N supply seen in *nlp2* and *nlp7* mutants, loss-of-function mutations in NLP2 or NLP7 result in a very different pattern of nitrate accumulation under non-limiting N supply, suggesting contrasting metabolic regulation between these two NLPs.

### The metabolic profiles of *nlp2* and *nlp7* mutants reveal contrasting acclimation under non-limiting nitrate supply

The contrasting nitrate accumulation pattern of *nlp2* and *nlp7* mutants under non-limiting nitrate supply described above prompted us to perform an untargeted gas chromatography–mass spectrometry (GC-MS) metabolite profiling of the rosettes from 42-day-old plants grown under limiting and non-limiting nitrate supply. We thus determined the relative contents of 105 detected and identified metabolites in *nlp2-2*, *nlp2-3* and *nlp7-1* mutants compared to their respective wild types (Supplemental Table S4). As all metabolites showed a similar trend for *nlp2-2* and *nlp2-3* compared to their WT, we displayed only the data for *nlp2-3* in Figure 5.

**Figure 5.**
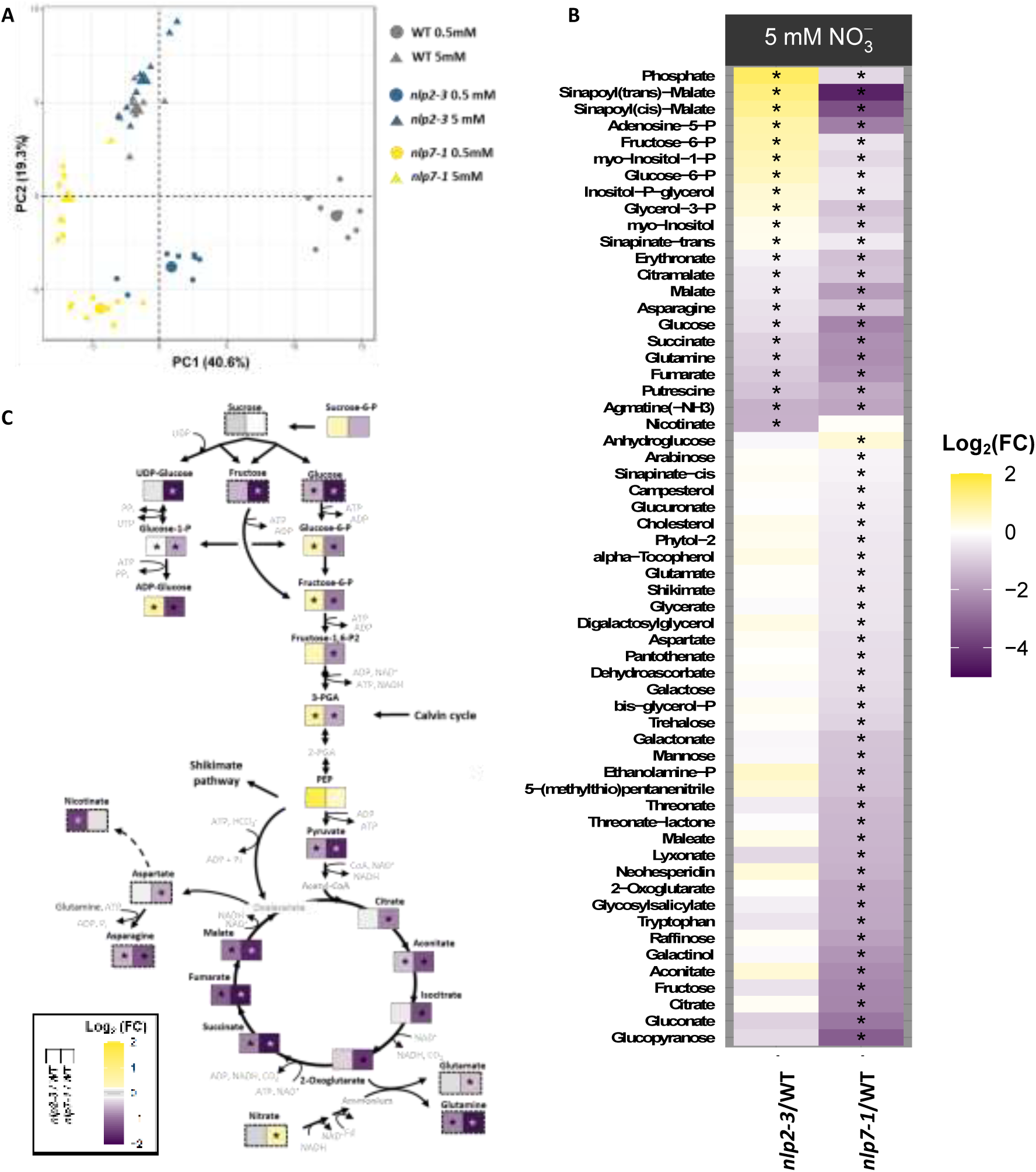
Rosette metabolite contents of *nlp2-3* and *nlp7-1* mutants grown under limiting and non- limiting nitrate supply. Plants were grown as in Figure 4. **(A)** Principal component analysis using genotype and nitrate supply as individuals and the measured contents of 105 metabolites as variables. Mean values are indicated as larger symbols. **(B)** Heatmap representation of the Log2(fold-change) in metabolite contents in *nlp2- 3* and *nlp7-1* compared to the WT under non-limiting nitrate supply. Purple, lower abundance; yellow, higher abundance. For A and B, data were from four plants per experiment, with two independent experiments and analyzed by GC-MS. **(C)** Major changes in central metabolism metabolites in the rosettes of *nlp2* and *nlp7-1* mutants compared to the WT. Data were obtained by LC-MS analysis from 3 independent experiments. Several metabolite data obtained by colorimetric assays or by GC-MS are included and indicated with a dashed line. The metabolic pathways are given without considering compartmentation in cell organelles and are not exhaustive for all metabolic pathways. Compounds indicated in light gray were not detected. *, *P*-value ≤ 0.05 as determined by a pairwise permutation test between the *nlp* mutants and the WT.

To compare the metabolite contents in *nlp2-3* and *nlp7-1* mutants, we performed a principal component analysis using genotype and nitrate supply conditions as individuals and the measured contents as variables. Principal component 1 (PC1) and PC2 explained 40.6% and 19.3% of the total variance, respectively (Figure 5A). Under limiting nitrate supply, both *NLP2* and *NLP7* loss-of-function mutants were accompanied by an almost equal position shift compared to the WT mainly along PC1, suggesting similar alterations for their shoot metabolite contents in comparison to the WT. Under non-limiting nitrate supply, WT and *nlp2-3* samples clustered together, with only the *nlp7-1* mutant occupying a different position in the PCA plot, mostly along PC2, suggesting that the *nlp2* metabolome only differs by a few compounds relative to that of the WT, despite the clear lower biomass in the mutant (Figure 5A).

To further decipher the contrasting metabolic phenotypes of *nlp2* and *nlp7* mutants under non-limiting N supply, we performed a non-parametric statistical analysis of the 105 measured metabolites in mutant and WT plants grown under non-limiting nitrate conditions and identified 59 metabolites displaying a significant (*P* ≤ 0.05) difference in either or both mutants in comparison to the WT (Figure 5B). We determined that the contents of 37 compounds are specifically altered in the *nlp7-1* mutant (36 down and 1 up), with 10 compounds showing lower contents in both *nlp2- 3* and *nlp7-1* mutants, 11 compounds showing higher contents in *nlp2-3* and lower contents in *nlp7-1*, and one compound showing specifically lower contents in *nlp2-3*.

As many metabolites of central metabolism exhibited differential accumulation, we further quantified the low-level phosphorylated metabolites by LC-MS in *nlp2-2*, *nlp2-3* and *nlp7-1* mutants compared to their respective WT. As the changes in metabolite levels were very similar between *nlp2-2* and *nlp2-3*, we show the data for one *nlp2* mutant in Figure 5 and data for both mutants are given in Supplemental Table S5. In the context of central metabolism (Figure 5C; Supplemental Table S5), both *nlp2* mutants showed lower contents for compounds belonging to the final part of the TCA cycle (succinate, fumarate, and malate), whereas the levels of all measured TCA cycle intermediates strongly decreased in *nlp7-1* rosettes (citrate, aconitate, 2- oxoglutarate, succinate, fumarate, and malate). Similarly, asparagine and glutamine accumulated to lower levels in both *nlp2* and *nlp7-1* mutants, while aspartate and glutamate contents only decreased in the *nlp7-1* mutant. NLP2 loss of function was specifically characterized by increased contents for phosphorylated compounds and inorganic phosphate, as well as a strong decrease in nicotinate contents.

These changes in metabolite levels in *nlp2* mutants were either opposite to those seen (phosphorylated compounds) or absent (nicotinate) in the *nlp7-1* mutant. In particular, the accumulation of 3-phosphoglycerate accompanied by a decrease in TCA cycle intermediates and an increase in sugar phosphates despite decreased or unmodified levels of free sugars, illustrate strong metabolic deregulations in the *nlp2* mutants. Thus, *nlp2* and *nlp7-1* mutants showed almost similar metabolic patterns when grown under nitrate-limiting conditions, while their metabolic profiles were clearly different under non-limiting nitrate conditions.

### Global gene expression patterns of *nlp2* and *nlp7* mutants grown under non- limiting nitrate supply supports distinct acclimation

As *nlp2* and *nlp7* displayed distinct metabolic patterns when grown under non-limiting nitrate supply, we performed a transcriptome analysis on roots and rosettes of 42-day- old plants grown under 5 mM nitrate supply. For each organ, we compared transcript levels in *nlp2-2*, *nlp2-3* and *nlp7-1* mutants relative to the WT (Figure 6; Supplemental Figure S9, Supplemental Tables S6, S7). We first compared *nlp2-2*/WT with *nlp2- 3*/WT DEGs and retained for further analyses genes that were differentially expressed in both *nlp2* mutants and in the same manner (up or down). Figure 6 and Supplemental Figure S9 display alterations in gene expression for *nlp2-3*. Data for *nlp2-2* are given in Supplemental Tables S6 and S7. A core set of genes showed similar differential expression compared to the WT in *nlp2-3* and *nlp7-1* mutants in roots (494 genes, 247 up, 247 down) and rosettes (307 genes, 291 down, 16 up) (corrected *P*-value ≤ 0.05; Log_2_ [mutant/WT] fold-change ≥ |0.4|; Figure 6A and C, Supplemental Figure S9). An additional set of 355 and 863 genes were specifically misexpressed in *nlp2-3* and *nlp7- 1* roots, respectively, compared to the WT. In rosettes, we observed mutant-specific differential expression patterns, with mainly lower steady state mRNA levels for *nlp2*/WT DEGs (318 out of 326), while *nlp7*/WT DEGs changed in both directions (427 up, 229 down).

**Figure 6.**
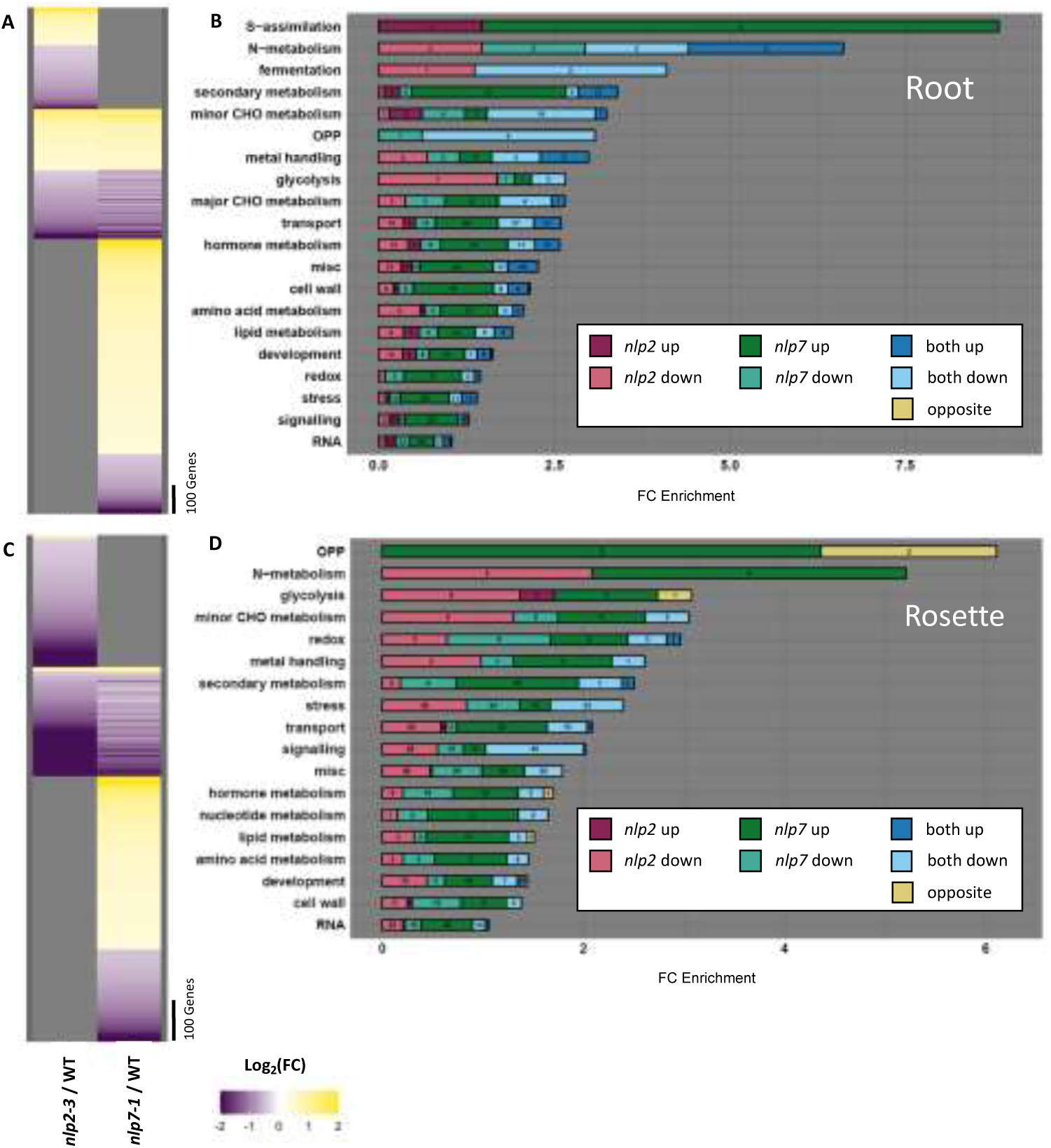
Transcriptome analysis of roots and rosettes of *nlp2-3* and *nlp7-1* mutants grown under non- limiting nitrate supply. Plants were grown as in Figure 4 and transcriptome analysis was performed by RNA-seq. **(A, C)** Heatmap representation of the Log2(fold-change) for transcript levels in the roots (**A**) and rosettes (**C**) of the *nlp2-3* and *nlp7-1* mutants compared to the WT under non-limiting nitrate supply. Purple, lower abundance; yellow, higher abundance. Gray, non-significant changes in gene expression in *nlp2-3* or *nlp7-1* compared to the WT. Significant differences in gene expression were determined according to Log2(fold-change) (Log2-FC ≥ 0.4 or Log2-FC ≤ –0.4) with a false discovery rate adjustment (*p*-value ≤ 0.05). **(B, D)** Enrichment in major GO terms related to DEGs in roots (**B**) and rosettes (**D**) in *nlp2-3* and/or *nlp7-1* compared to the WT. Blue, DEGs in both *nlp2-3* and *nlp7-*1; green, DEGs in *nlp7-1*; violet, DEGs in *nlp2-3*. Dark colors, upregulated genes; light colors, downregulated genes, compared to the WT. Yellow, opposite gene deregulation between *nlp2-3* and *nlp7* compared to the WT. Black numbers indicate the number of genes associated within each class.

To further decipher the consequences of the loss of NLP2 or NLP7 function, we focused on significantly enriched processes (enrichment ≥ 1;*P*-value ≤ 0.05) by looking at the number of DEGs either specific or in common to both mutants compared to the WT (*nlp2* up, *nlp2* down, *nlp7* up, *nlp7* down, both up, both down, and opposite, Figure 6, B and D). At first approximation, DEGs in *nlp2* and *nlp7* mutants were enriched in MapMan GO terms closely linked to metabolism, but also development, signaling, and RNA. Remarkably, despite several *nlp2*/WT and *nlp7*/WT DEGs in common, in most GO classes, the loss of function of either NLP2 or NLP7 modified the expression of different genes belonging to the same GO category. For example, in roots (Figure 6B), the transcript levels of several N metabolism genes specifically decreased in one of both mutants, whereas other genes were differentially expressed in both mutants, with some genes being less strongly upregulated and downregulated in *nlp2* mutants than in *nlp7-1* (Supplemental Table S8). The GO term ‘glycolysis’ was highly enriched among DEGs specific for the *nlp2* mutant. The loss of NLP7 function also led to specific alterations in gene expression that did not occur in *nlp2* mutants, e.g., for genes related to S-assimilation and secondary metabolism.

In rosettes, we observed similar enriched GO terms as in roots, but NLP2 and NLP7 loss of function altered transcript levels in opposite ways, as for example with N metabolism–associated genes. As in roots, several genes related to glycolysis were specifically downregulated in the rosettes of the *nlp2* mutant (Supplemental Table S8).

We next looked for the extent of overlap between genes bound by NLP2 after 30 min of nitrate resupply at the seedling stage and the nlp2/WT DEGs in 42-day-old plants (Supplemental Figure S10). Despite the different age and growth conditions of the material used for ChIP-seq and RNA-seq, the overlap of bound targets and DEGs supported our finding that NLP2 is predominantly involved in the integration of nitrate assimilation and carbon metabolism in response to nitrate. In addition, *ASPARTATE OXIDASE* (*AO*) is among the 15 genes that are direct NLP2 targets and whose expression depended on NLP2 in adult plants grown under non-limiting nitrate supply. The direct regulation of *AO* pointed to a link with NAD metabolism, as AO catalyzes the first step of NAD biosynthesis. Taken together, the analysis of DEGs in individual GO classes clearly revealed the complementary pattern of DEGs related to NLP2 and NLP7 loss of function for most enriched processes and the regulatory role of NLP2 in the expression of genes involved in glycolysis, in addition to those involved in N metabolism.

### Double mutant phenotypes support specific roles of NLP2 and NLP7

To dissect the shared and specific roles played by NLP2 and NLP7 in regulating nitrate response and plant homeostasis, we generated the *nlp2-3 nlp7-1* double mutant. We first measured by RT-qPCR the relative transcript levels of dozens of genes during a time course of N-starved seedlings following nitrate resupply in the WT, single mutants, and the *nlp2-3 nlp7-1* double mutant.

We organized these genes into four classes according to their nitrate response pattern in single and double mutants (Figure 7; Supplemental Figure S11). Class I consisted of genes that do not exhibit additional differential expression in the double mutant relative to the *nlp2-3* or *nlp7-1* single mutants (e.g., *HYH* and *PHOSPHOGLYCERATE MUTASE* [*PGM*]). Class II comprised genes with accentuated gene expression changes in the double mutant compared to either single mutant (e.g., *HHO1* and *BETA-CARBOANHYDRASE4* [*BCA4*]), suggesting additive effects of NLP2 and NLP7. Class III and IV included genes specifically or more prominently regulated by NLP2 (*GLYCERALDEHYDE-3-PHOSPHATE DEHYDROGENASE OF PLASTID 2* [*GAPCP2*] and *FUMARASE 2* [*FUM2*]) or by NLP7 (*NIR1* and *NRT2.1*), respectively. Thus, in addition to some degree of functional redundancy and additive effects between NLP2 and NLP7, the nitrate-responsive expression patterns observed for class III and IV genes argue in favor of functions specific for NLP2 or NLP7.

**Figure 7.**
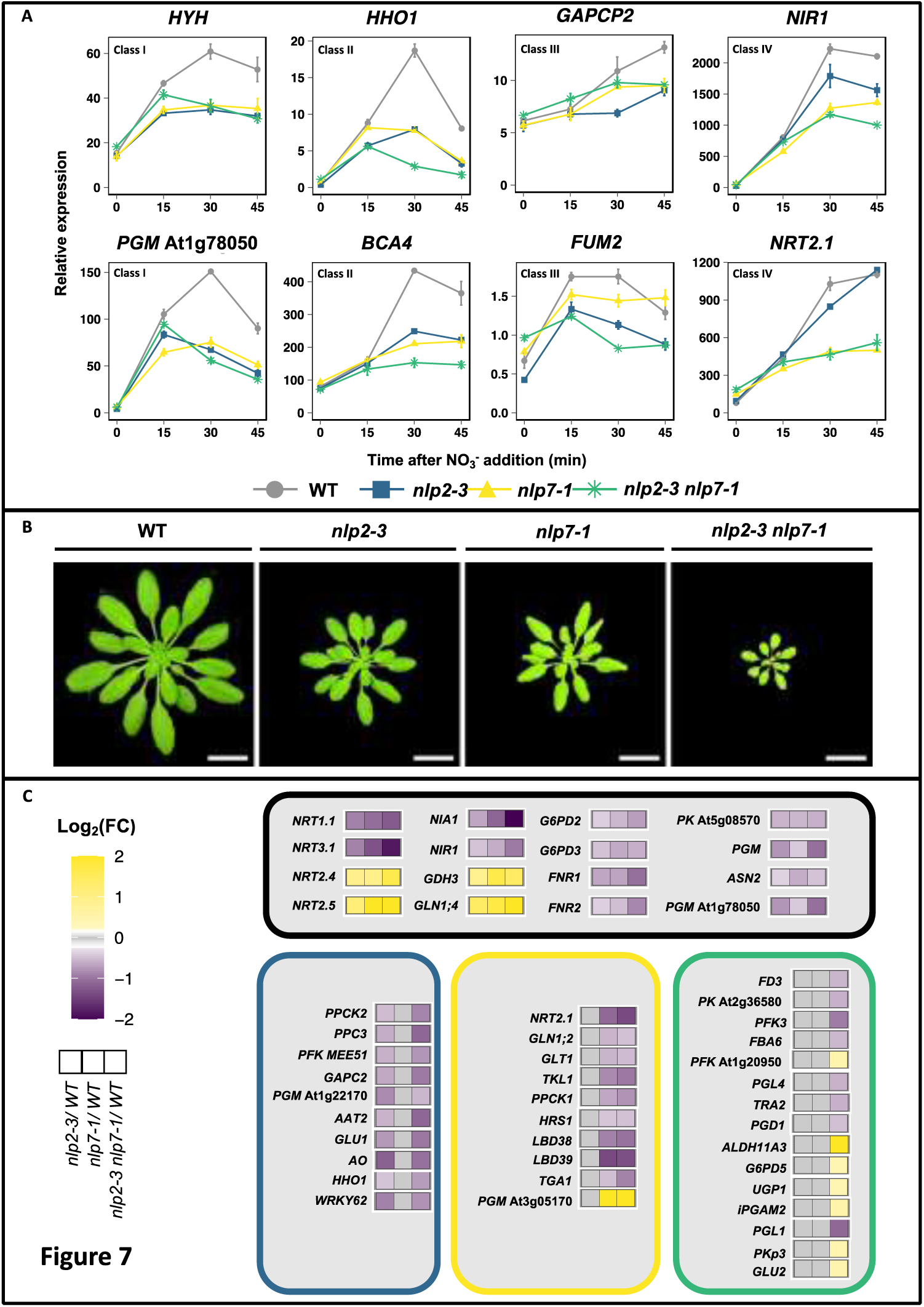
Molecular and physiological alterations in the double *nlp2-3 nlp7-1* mutant suggest distinct roles for NLP2 and NLP7. **(A)** Relative transcript levels of selected genes after nitrate addition to N-starved seedlings for WT (gray), *nlp2-3* (blue), *nlp7-1* (yellow), and *nlp2-3 nlp7-1* (green). Seedlings were cultivated and treated as in Figure 2. Expression was measured by RT-qPCR and normalized as a percentage of the expression of a constitutive synthetic gene composed of *TIP41* (At4g34270) and *ACTIN2* (*ACT2*, At3g18780). Data are means ± standard error (SE, n = 4) of one representative experiment. **(B)** Top view of 42-day-old rosettes from the WT, *nlp2-3*, and *nlp7-1* grown under non-limiting (5 mM) nitrate supply. **(C)** Principal DEGs identified by RNA-seq in 42-d-old roots from *nlp2-3*, *nlp7-1*, and *nlp2-3 nlp7- 1* compared to the WT. Black, DEGs in all mutants; blue, DEGs specific to *nlp2-*3; yellow, DEGs specific to *nlp7-1*; green, DEGs specific to *nlp2-3 nlp7-1*. All presented DEGs show significant expression changes with Log2(fold-change) ≥ 0.4 or ≤ –0.4 with false discovery rate adjustment (*P*-value ≤ 0.05).

We also measured the biomass accumulation of all genotypes (the WT, single mutants, and the *nlp2-3 nlp7-1* double mutant) under non-limiting nitrate supply conditions. Interestingly, the growth defects observed for the *nlp2* and *nlp7* mutants were even more pronounced in the *nlp2-3 nlp7-1* double mutant (Figure 7B), likely underscoring the additive effect between NLP2 and NLP7.

At the transcript levels in the roots of 7-week-old *nlp2-3 nlp7-1* double mutant plants measured by RNA-seq, we obtained further support for additivity and functional redundancy between NLP2 and NLP7, as several genes belonging to N metabolism and related processes displayed similar expression levels (*NRT1.1*, *GLN1;4*, and *G6PD3*) or stronger differential expression (*NRT2.4*, *NRT2.5*, *NIA1*, *NIR1*, *FNR1*, and *FNR2*) compared to the single mutants (Figure 7). Interestingly, several DEGs specifically associated with the loss of function of either NLP2 (*PPCK2*, *GAPC2*, *PGM*, *AO*, and *HHO1,* Figure 7C) or NLP7 (*NRT2.1*, *GLT1*, *TKL1*, and *HRS1,* Figure 7C) were also misexpressed in the roots of the *nlp2-3 nlp7-1* double mutant, confirming the specificity of some functions between NLP2 and NLP7. Furthermore, the joint loss of NLP2 and NLP7 function led to additional DEGs closely associated with N metabolism and related processes (e.g., *FERREDOXIN 3* [*FD3*], *PGD1*, and *PFK3,* Figure 7C), illustrating the important complementary role played by both NLP2 and NLP7 in the regulation of the nitrate assimilation pathway and associated processes.

### *nlp2* and *nlp7* mutants show distinct growth phenotypes under non-limiting ammonium nitrate supply

Because *nlp2* and *nlp7* single mutants showed perturbations associated with the nitrate assimilation pathway and closely related processes, we grew *nlp2*, *nlp7*, and *nlp2-3 nlp7-1* mutants in the presence of 2.5 mM ammonium nitrate corresponding to a 5 mM N dose (non-limiting N supply) but with a reduced N source. Non-limiting ammonium nitrate nutrition increased biomass accumulation in the WT (2.1-fold) but also in the *nlp2-3* mutant (4.1-fold) when compared with the respective genotypes grown in non-limiting 5 mM nitrate conditions. Although the biomass of the *nlp2-3* single mutant only reached 60% that of the WT under non-limiting ammonium nitrate nutrition, the difference between genotypes was less pronounced than under non-limiting nitrate supply, when the *nlp2-3* single mutant only produced 30% of the biomass measured in the WT, indicating that ammonium nitrate supply partially rescues the non-limiting nitrate-dependent growth defect of the *nlp2-3* mutant (Figure 8).

**Figure 8.**
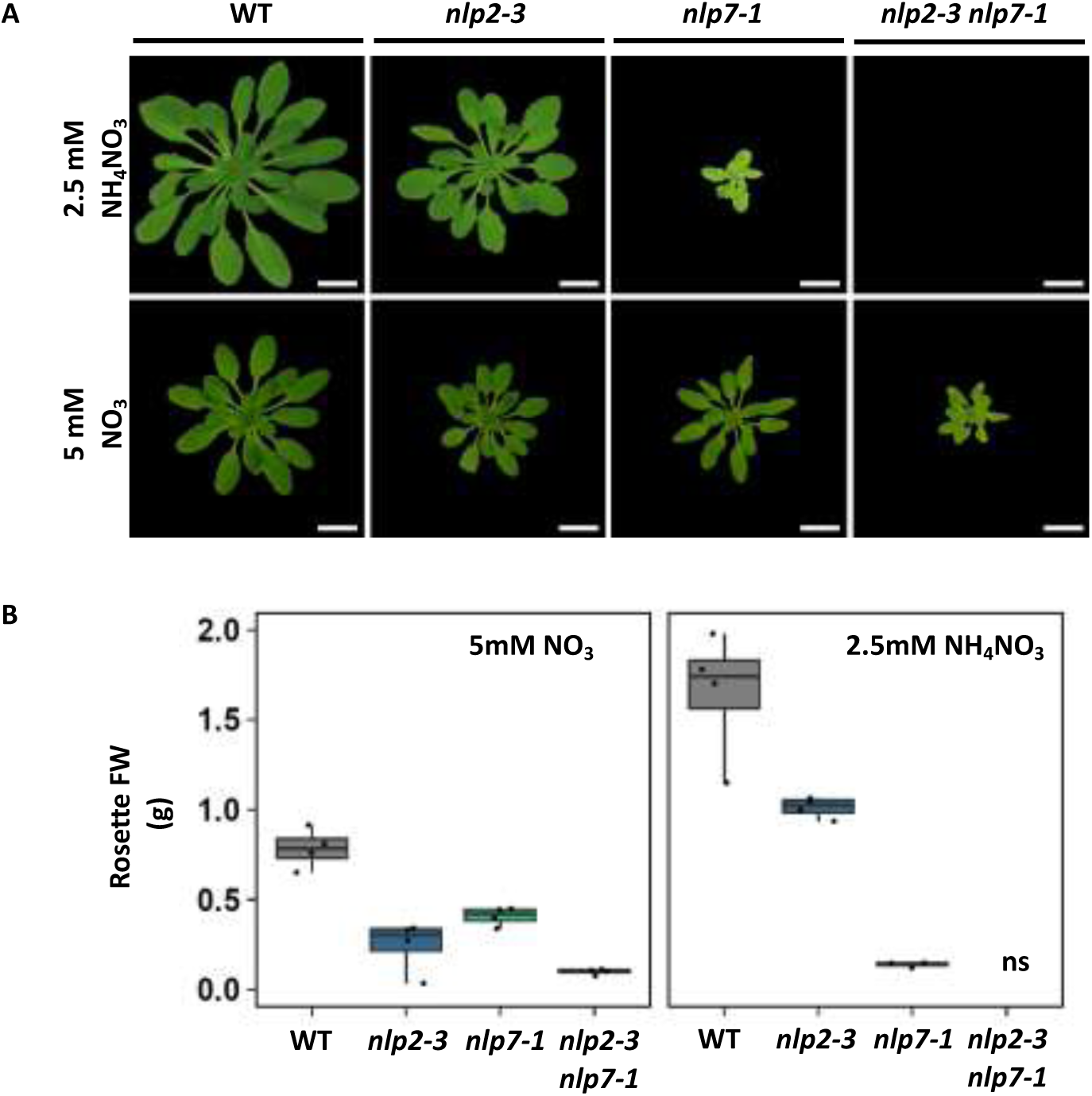
Growth under non-limiting ammonium nitrate supply distinguishes *nlp2* and *nlp7* mutants. Plants were cultivated as in Figure 4. **(A)** Top view of 42-day-old rosettes from the WT, *nlp2-3*, *nlp7-1*, and *nlp2-3 nlp7-1* grown under non-limiting (2.5 mM) ammonium nitrate and non-limiting (5 mM) nitrate supply in short-day conditions. Scale bar = 2 cm. **(B)** Rosette fresh weights. On the left, plants cultivated on 5mM nitrate nutrient solution, on the right plants cultivated on 2.5mM NH4NO3 nutrient solution Data were obtained from 4 plants. ns: no survivor.

Surprisingly, growth of the *nlp7-1* mutant was dramatically impaired under non-limiting ammonium nitrate nutrition compared to non-limiting nitrate supply, experiencing a 70% decrease in biomass when grown with 2.5 mM ammonium nitrate and only accumulating 10% of the biomass seen in the WT, revealing a striking difference between the *nlp2-3* and *nlp7-1* mutants. Moreover, loss of both NLP2 and NLP7 function was almost seedling-lethal under ammonium nitrate nutrition, as the *nlp2-3 nlp7-1* double mutant wasted away around the four-leaf stage, indicating that the additional loss of NLP2 function severely exacerbated the *nlp7-1* growth defect observed under ammonium nitrate supply (Figure 8). These results further illustrate the additivity and the specificity suggested between NLP2 and NLP7.

Taken together, the use of an alternative N source clearly discriminates several aspects of the specificities related to either NLP2 or NLP7 loss of function. Indeed, while NLP2 and NLP7 are crucial for sustaining nitrate-dependent growth, the use of ammonium nitrate partially rescued the *nlp2* growth defect but worsened that of *nlp7*.

### NLP2 and NLP7 directly interact and accumulate in shared and distinct root cell types

The additivity of the *nlp2* and *nlp7* single mutants raised the possibility that these proteins may physically interact and form heterodimers, as shown previously for other NLP proteins (Konishi and Yanagisawa, 2019). Using a bimolecular fluorescence complementation (BiFC) approach in *N. benthamiana* leaves, we detected an interaction between NLP2 and NLP7, as evidenced by the reconstitution of yellow fluorescent protein (YFP) in the nucleus of *N. benthamiana* leaves infiltrated with the constructs *YFPn-NLP2* and *YFPc-NLP7* from nitrate resupplied plants (Figure 9, D, E and F). By contrast, we observed YFP fluorescence only in the cytosol in N-starved plants (Figure 9, A, B and C), in agreement with the localization of the individual proteins (Supplemental Figure S12).

**Figure 9.**
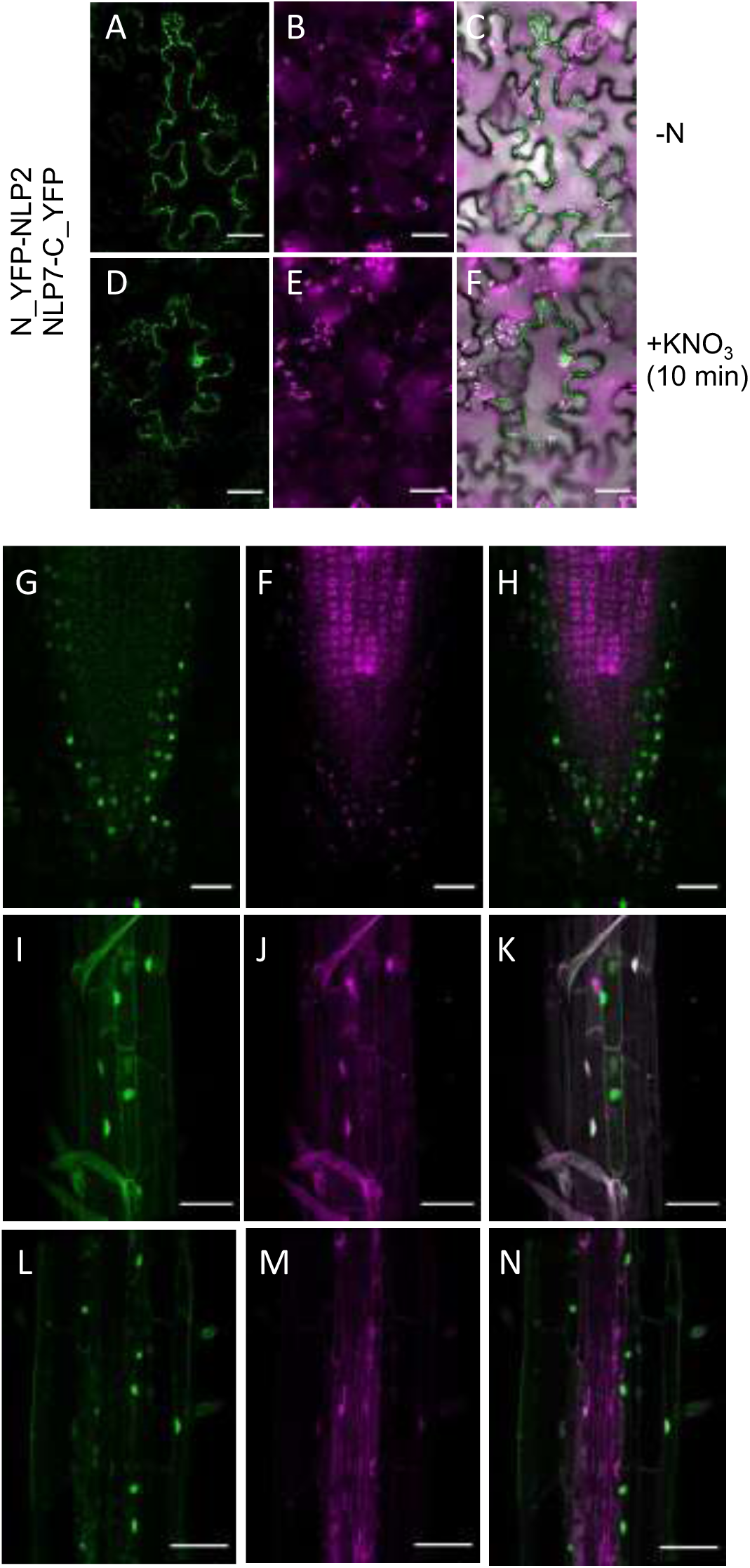
Interaction and cellular distribution of NLP2 and NLP7. Plants were cultivated as in Figure 1. **(A-E)** NLP2 and NLP7 directly interact in BiFC assays in the absence (**A-C**) and presence of nitrate (**D-E**). Scale bar = 50 µm. (**G-N**) Confocal imaging of *NLP2pro:NLP2-mCherry* and *NLP7pro:GFP-NLP7* lines at the root tip **(G-H)** and root differentiation zone **(I-N).** Scale bars = 50 µm.

We also analyzed the spatial accumulation domains of both proteins in nitrate-treated Arabidopsis roots from seedlings harboring both the *NLP2pro:NLP2-mCherry* and *NLP7pro:GFP-NLP7* transgenes. We mainly detected GFP-NLP7 in the root columella and epidermal cells, whereas NLP2-mCherry strongly accumulated in cortex and stele cells (Figure 9, G and H). In the differentiation zone, both NLP2 and NLP7 accumulated in the cortex and pericycle cells (Figure 9, I to N), with nuclei containing either one or both proteins.

## Discussion

### NLP2 is a central player of the early nitrate response GRN

Over the last decade, NLP7 has emerged as one of the major transcription factors governing the primary nitrate response in Arabidopsis (Marchive et al., 2013; Alvarez et al., 2020). However, nitrate responses are not completely abolished in single *nlp7* mutants (Castaings et al., 2009; Marchive et al., 2013), which is similar to mutants in other nitrate signaling-related TFs (reviewed in Vidal et al., 2020). These observations suggest the existence of a complex GRN that other members of the NLP family might contribute to, as recently suggested based on growth and gene expression analysis of higher order *nlp* mutants (Guan et al., 2017; Konishi et al., 2021).

Here, we identified NLP2 as a key player in the early nitrate response GRN, as illustrated by our ChIP-seq results showing that nitrate-triggered NLP2 binding is chiefly directed toward dozens of strong nitrate-responsive genes (Figure 1 and Figure 2), including a majority of those highlighted as most consistently regulated by nitrate (Canales et al., 2014). We established that NLP2 binds genes related to nitrate metabolism including N transport, N assimilation, and the OPP, as well as many of the most influential genes encoding TFs participating in the nitrate GRN (*TGA1*, *TGA4*, *LBD37*, *LBD38*, *LBD39*, *HRS1*, *HHO2*, *HHO3*, *COL5*, *HYH*, *TCP23*, *BEE2*, and *ABF2*) (reviewed in Vidal et al., 2020). This result points to the existence of direct and indirect connections between NLP2 binding to its target genes and the transcriptional regulation of nitrate assimilation–related metabolism.

RNA-seq analysis of the *nlp2-2* mutant and the WT under the same nitrate resupply conditions used for ChIP-seq confirmed that NLP2 acts as a master upstream TF triggering a regulatory cascade. Indeed, at least 75% of the perturbations resulting from the loss of NLP2 function can be explained by NLP2 binding and by several of the NLP2-dependent downstream events. Interestingly, the NLP2-TF_2/3_-dependent downstream events also contributed to the direct NLP2-dependent transcriptional wave, since 75% of the *nlp2-2* nitrate-regulated DEGs bound by NLP2 were also predicted to be bound by at least one NLP2-TF_2_ and/or NLP2-TF_3_ member. These multi-tier regulations via TF_2/3_s acting as positive or negative feedback loops are consistent with recent observations underscoring the robustness of GRNs imparted by such feedback loops (Hebbar et al., 2021).

Dual transcriptional reinforcement likely increases the robustness of the expression of NLP2 target genes with key functions in the nitrate response, as described in other systems (Hemming et al., 2021; Osterwalder et al., 2014). Indeed, in addition to master regulators, a complex GRN such as the nitrate-response GRN probably requires other TFs that facilitate rapid, coordinated, and dynamic signaling similar to what was demonstrated for ABA signaling (Song et al., 2016). Taken together, we demonstrated that NLP2 is a major orchestrator of the early nitrate response-related GRN. The positioning and role of NLP2 in relation to NLP7 constitute a major breakthrough allowing elucidation of the nitrate response–related GRN.

### NLP2 co-orchestrates the early nitrate response GRN together with NLP7

Motif enrichment analysis of the genomic sequences associated with the NLP2 binding site suggested its high affinity for a DNA consensus sequence (TGNCYYTT) identified in all regions bound by NLP2. This sequence was highly similar to the motif associated with the NLP7 binding site identified by DAP-seq (O’Malley et al., 2016) and strongly resembled the 5′ part of the pseudo palindromic NRE described previously (Konishi and Yanagisawa, 2010). In addition, almost 20% of the NLP2-binding sites were enriched in the full pseudo palindromic NRE motif (TGNCYYTT[N_8-12_]AARRG), confirming *in planta* and globally that several NLPs, including NLP2 and NLP7, bind to the NRE in the *NIR1* promoter, as shown previously by yeast one-hybrid analysis (Konishi and Yanagisawa, 2013).

The presence of either the full pseudo palindromic binding sequence or solely the 5′ region suggests that NLP2 binds either as a dimer or as a heterodimer, as NLP proteins have been shown to dimerize via their PB1 (Phox and Bem1) domain (Konishi and Yanagisawa, 2019). Moreover, the NLP2-bound genes identified in this work substantially overlapped with NLP7-bound genes (Marchive et al., 2013; O’Malley et al., 2016; Alvarez et al., 2020), suggesting that the RWP-RK DNA binding domains of both NLP2 and NLP7 share a high affinity *in planta* for the same consensus DNA sequence. Thus, our data strongly indicate that NLP2 and NLP7 have similar binding properties *in planta*, which is reflected by the significant overlap between NLP2-bound DEGs and previously identified NLP7-bound DEGs in response to nitrate resupply (Marchive et al., 2013) (Supplemental Figure S13). The accumulation pattern of NLP2 and NLP7 in various tissues also supports this notion, as both proteins were present in root cortex and pericycle cells.

Not surprisingly, we showed redundant and additive effects associated with the loss of NLP2 and NLP7 function by dissecting the early nitrate response in the WT, the *nlp2-3* and *nlp7-1* single mutants, and the *nlp2-3 nlp7-1* double mutant. Thus, in a cell accumulating both NLP2 and NLP7, genes regulated additively by NLP2 and NLP7 are candidate targets for an NLP2/NLP7 heterodimer, which deserves further study. Notably, we also showed that NLP2 and NLP7 have specific roles in the primary nitrate response, as evidenced by the partial overlap in their target gene repertoire and the often distinct transcriptome perturbations associated with the respective loss-of-function mutants in response to nitrate resupply. Interestingly, although co-expressed in several tissues, *NLP2* and *NLP7* were also expressed in distinct areas of the root: the epidermal cells for *NLP7* and the vascular tissue for *NLP2*. Thus, the spatial distribution of NLP2 and NLP7 may partially explain differences in gene expression specifically related to either NLP2 or NLP7 loss of function. However, constructs swapping the *NLP7* and *NLP2* promoter to drive *NLP2* or *NLP7* transcription only complemented the *nlp2* mutant phenotype with the *NLP2* coding sequence regardless of the promoter used (Konishi et al., 2021), pointing to a more complex mechanism explaining the specificities of NLP2 and NLP7. Indeed, other TFs such as ABF2 and ABF3 support a highly localized transcriptional response to nitrate in the pericycle, and other cell type–specific TF-target interactions have been predicted (Contreras-López et al., 2022). Taken together, our data strongly indicate that NLP2 co-orchestrates the primary nitrate response together with NLP7, likely in a cell-type-specific manner.

### NLP2 and NLP7 integrate N availability to regulate plant growth

Phenotypic analyses of *nlp2-3* and *nlp7-1* single mutants under limiting and non-limiting nitrate supply revealed that both NLP2 and NLP7 are important for an efficient use of nitrate as the sole N source, since both mutants were largely unable to convert additional nitrate into a substantial growth enhancement. Under non-limiting nitrate supply, roots of *nlp2* and *nlp7-1* single mutants displayed molecular features of N- starved seedlings, since DEGs related to either *nlp2*or *nlp7-1* mutants markedly and significantly overlapped with N starvation–responsive genes (Krapp et al., 2011, Supplemental Figure S14), in agreement with results reported for the *nlp7-1* mutant and extending these observations to the *nlp2-3* mutant (Castaings et al., 2009). Thus, the activity of NLP2 and NLP7 is essential for integrating nitrate availability toward the establishment of nitrate-associated signaling and metabolic pathways in roots. Consequently, loss of NLP2 and NLP7 function would be expected to limit the supply of nitrate and N compounds from the root to the rosette, likely explaining the growth defects reported for *nlp2-3* and *nlp7-1* mutants under non-limiting nitrate supply. However, *nlp7-1*, but not *nlp2-3*, accumulated about 1.5-fold more nitrate in the rosette under non-limiting nitrate supply than the WT, a feature already reported and previously attributed to the lower NR-activity in the rosettes of the *nlp7-1* mutant (Castaings et al., 2009). This difference in nitrate contents between *nlp7* and *nlp2* mutants was consistent with the levels of NR activity measured here, which only decreased significantly in the rosettes of the *nlp7-1* mutant grown under non-limiting nitrate supply. We speculate that lower NR activity cannot explain the growth or nitrate accumulation phenotypes observed here, as plant growth was shown to suffer under standard conditions only when NR activity falls below 10% of that of WT plants (Wilkinson and Crawford, 1983). Nevertheless, such distinct patterns concerning rosette nitrate metabolism revealed highly specific consequences for the loss of function of either NLP2 or NLP7, despite the similar growth response to nitrate availability seen in the single mutants, and raise questions about the nature of the specificity of NLP2 and NLP7.

### NLP2 ensures the high supply of reductants for nitrate assimilation

Our data revealed that NLP2 directly regulates genes encoding components of the OPP in response to nitrate resupply in seedlings. Similarly, we observed changes in gene expression and metabolite levels related to the OPP, such as Glc-6P and Fru- 6P, in adult *nlp2* mutants grown under nitrate supply. The OPP is one of the critical steps in providing reductants for nitrate assimilation (Kruger and von Schaewen, 2003). Our results suggest that NLP2 tightly coordinates the supply of reductants with the demand for nitrate reduction in a nitrate-dependent manner. Moreover, NLP2 appeared to directly tune NAD metabolism, since the NLP2-specific target gene *AO* in response to nitrate encodes an aspartate oxidase catalyzing the first step of de novo NAD biosynthesis (Hao et al., 2018). *AO* expression was specifically downregulated in both roots and rosettes of *nlp2* mutants under non-limiting nitrate supply. In addition, the contents of nicotinate, an intermediate of the NAD salvage pathway, accumulated to lower levels in the rosettes of *nlp2* but not *nlp7* mutants, reinforcing the specific link between NLP2 and NAD metabolism (summarized in Supplemental Figure S15). Impairment of NAD and OPP metabolism due to NLP2 loss of function would limit nitrate and nitrite reduction, particularly when nitrate is supplied as a sole N source in the soil, which likely prevents the efficient conversion of additional nitrate into growth enhancement, as observed here. Based on these indications, we propose that NLP2 orchestrates the processes required for energy supply to sustain nitrate assimilation, particularly when soil nitrate is non-limiting.

### NLP2 connects glycolysis and the TCA cycle to nitrate assimilation

We showed that several steps of glycolysis are directly (*GAPCP2*, *GAPC2*, *PGM*, At1g22170) or indirectly regulated by NLP2 in response to nitrate. In addition, we measured lower transcript levels for several genes related to glycolysis (*PFK3*, *GAPC2*, *Phosphoglycerate mutase [iPGAM1*], and *PYRUVATE KINASE* [*PK*]), together with the accumulation of inorganic phosphate and phosphorylated intermediary metabolites associated with glycolysis (Glc6-P, Fru6-P, Fru-1,6-BP, 3- PGA, and PEP) in the rosettes of *nlp2* mutants under non-limiting nitrate supply. Such a pattern argues in favor of a slowdown of glycolysis in *nlp2* mutants. More precisely, the misexpression of genes associated with glycolysis in *nlp2* mutants appeared to mainly affect the conversion of glyceraldehyde-3-phosphate to phosphoglyceric acid (PGA) (via GAPDH, phosphoglycerate kinase, and PGM), stressing the central role of NLP2 in regulating glycolysis. Indeed, *iPGAM1* is one of the most strongly and robustly induced genes in response to nitrate supply (Wang et al., 2000; Canales et al., 2014) and encodes the enzyme that catalyzes the conversion of 3-PGA to 2-PGA, thereby exporting organic carbon from the photosynthetic Calvin-Benson-Bassham cycle. This strong regulation by nitrate is probably due to the particular importance of glycolysis as an energy and carbon skeleton supply source. Moreover, the NLP2-dependent regulation of *GAPCP*s is noteworthy, as they encode metabolic connectors between glycolysis with other pathways such as N metabolism (Anoman et al., 2015). Also, the identification of a moonlighting role for a complex comprising PGM, enolase, and PK in the association of chloroplastic and mitochondrial activities that would be beneficial for metabolic pathways that span organelles, such as photorespiration and N assimilation (Zhang et al., 2020), is intriguing, as the genes encoding these proteins are less expressed in *nlp2* mutants. This might indicate that the coordination of organelle activities is disturbed in *nlp2* mutants. Thus, our data strongly suggest that NLP2 is a major player in the nitrate-dependent regulation of glycolysis that supports provision of energy and C skeletons for N metabolism.

Moreover, nitrate limitation is accompanied by a decrease in TCA cycle intermediates and pyruvate in adult plants under non-fluctuating nutrient conditions (Tschoep et al., 2009; Fataftah et al., 2018). We noticed such a metabolic reprogramming in the *nlp2* and *nlp7* mutants grown under non-limiting nitrate supply (Supplemental Figure S16A). This observation highlights the potential involvement of NLP2 in the regulation of central C metabolism, especially since fumarate and malate, just like starch, represent a high proportion of transient and non-structural C in the Arabidopsis rosette (Chia et al., 2000; Gibon et al., 2009). Indeed, the genes encoding PEP carboxylase and β-carboanhydrase that both participate greatly in the filling of C skeletons into the TCA were both directly regulated by NLP2 and less expressed in *nlp2* mutant roots (Supplemental Figure S10). In addition, *PK* expression, encoding an enzyme catalyzing another anaplerotic pathway for filling the TCA cycle, was indirectly regulated via NLP2-TF_2/3_ and less expressed in root and rosettes of *nlp2* mutants.

Interestingly, we also noticed an *nlp2*-specific decrease in the expression of *FUM2*, encoding a cytosolic fumarase (Supplemental Figure S16B) that is required for the accumulation of fumarate in leaves, itself necessary for rapid nitrogen assimilation and growth with a high level of nitrogen (Pracharoenwattana et al., 2010). However, how *FUM2* transcript levels are downregulated in *nlp2* mutants remains unclear, since neither NLP2 nor NLP2-TF_2/3_s bind to the *FUM2* locus. However, the targets of more TF_2/3_s remain to be investigated. We also cannot exclude the possibility that the alteration of fumarate metabolism is a consequence of the lower energy supply for nitrate assimilation resulting from the loss of NLP2 function.

Taken together, our data show that NLP2 is a key regulator, both directly and indirectly, of carbon metabolism, likely to sustain the provision of C skeletons for N assimilation and support plant growth under non-limiting nitrate supply.

### The *nlp2* and *nlp7* mutants show distinct phenotypes when grown under mixed N supply

Non-limiting ammonium nitrate supply partially rescued the growth defect of *nlp2* mutants while it exacerbated the poor growth of the *nlp7-1* mutant, and even more so for the *nlp2-3 nlp7-1* double mutant, relative to growth under non-limiting nitrate supply. These observations contradict the recently reported rescue of the nitrate-dependent growth defect of *nlp7*, but not *nlp2*, mutant seedlings by ammonium when grown on agar plates containing sucrose (Konishi et al., 2021). Indeed, supply of ammonium in addition to nitrate stimulates ammonium assimilation by bypassing nitrate reduction as the rate-limiting step of N assimilation, which, in turn, requires high levels of C skeletons (Masakapalli et al., 2013; Hachiya and Sakakibara, 2017). As a consequence, carbon availability is essential for maximizing organic N compound biosynthesis when both nitrate and ammonium are available as N sources. Thus, *nlp7* mutants that show a global C depletion are unable to cope with this condition. The presence of sucrose in the medium might be central for the observed rescue of the growth defect of *nlp7* seedlings grown on ammonium nitrate medium and its ability to efficiently use ammonium nitrate as N supply. The growth recovery of *nlp2* mutants cultivated under non-limiting ammonium nitrate supply under our conditions is consistent with the putative NLP2 loss-of-function-dependent impairment of energy supply through perturbations of NAD and OPP metabolism, as ammonium nutrition is less energetically costly than nitrate nutrition.

In conclusion, our data uncovered the role of NLP2 as a major molecular regulator of early gene expression responses to a nitrate signal, orchestrating N assimilation and central carbon metabolism either directly or indirectly via a cascade of transcription factors (Figure 10A). Loss-of-function mutations in NLP2 or NLP7 altered growth under non-limiting nitrate supply, which is likely due to the impairment of the nitrate assimilation pathway. However, in addition to these common features, our results strongly suggest additional and specific causes responsible for the growth defects observed in each *nlp* mutant, based on specific metabolic and transcriptome changes in each mutant. Such a dichotomy is particularly relevant under non-limiting ammonium nitrate supply because this alternative N source allows partial rescue of the *nlp2* growth defect while aggravating the *nlp7* growth defect. We propose a model that explains the growth impairment of *nlp2* mutants cultivated under constant non-limiting nitrate supply by effects not only on N assimilation but also on carbon and energy supplying pathways (Figure 10B).

**Figure 10.**
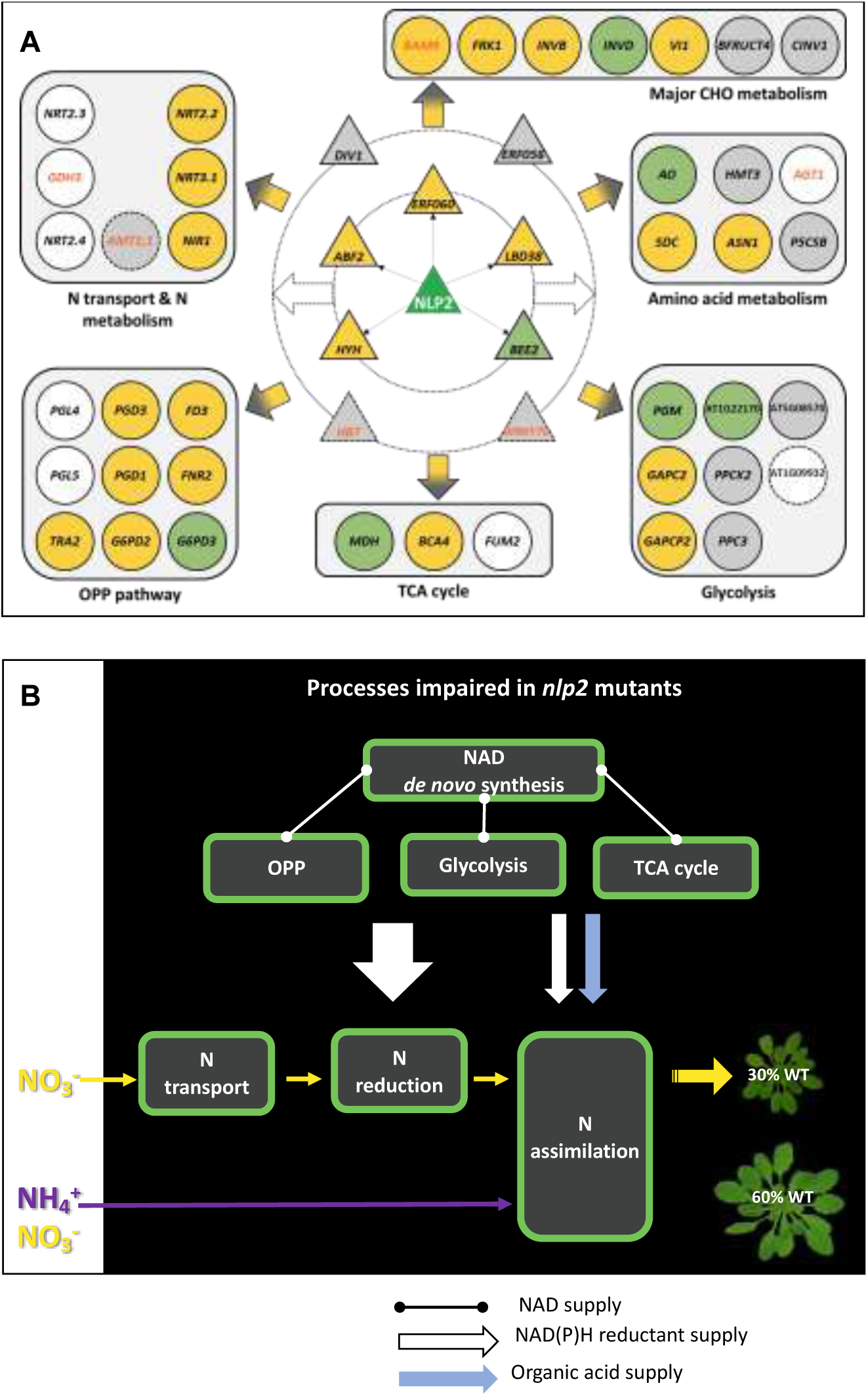
Schematic representation of the major NLP2-orchestrated early nitrate response pathways and a summary of *nlp2* mutant impairment in NAD, C, and N metabolism. **(A)** NLP2 directly and indirectly regulates many steps of N metabolism and C metabolism, in addition to many N-influential transcription factors. Triangles and circles represent TFs and genes associated with primary metabolism, respectively. Green, genes only bound by NLP2; yellow, genes bound by NLP2 and TF2/TF3; green, genes only bound by NLP2; gray, genes only bound by TF2/TF3; white, genes not bound by NLP2 or TF2/TF3. Dashed and solid lines represent *nlp2-2* DEGs compared to the WT after nitrate treatment, with dashed line for upregulated genes and a solid line for downregulated genes. Orange names, downregulated genes in the WT in response to nitrate; black names, upregulated genes in the WT in response to nitrate. Gene names are provided in Supplemental Table S10. **(B)** Impairments in NAD, C, and N metabolism related to NLP2 loss of function affects biomass accumulation under nitrate supply, while ammonium nitrate feeding allows a partial bypass of the limiting nitrate reduction step.

## Materials and Methods

### Plant materials

The mutant *nlp2-2* (SK19641) was derived from a T-DNA–mutagenized population of the Columbia-4 (Col-4) accession (Robinson et al., 2009), while the mutant *nlp2-3* (SAIL_139_D05) was derived from a T-DNA–mutagenized population of the Col-0 accession (Alonso-Blanco et al., 2003). All mutant seeds were obtained from the NASC and homozygous lines were identified by genotyping using the primers listed in Supplemental Table S9. The absence of transcripts was confirmed by RT-PCR (Supplemental Figure S4). The mutant *nlp7-1* (SALK_26134) was in the Col-0 (WT) background (Castaings et al., 2009). The *nlp7-1*, the *nlp2-2* and nlp2-3 mutants were backcrossed twice to their respective wild-type strain. The double mutant *nlp2-3 nlp7- 1* was obtained by crossing single mutants and PCR genotyping of the F2 progeny using primers listed in Supplemental Table S9.

The *nlp2-2* mutant was used for complementation with *NLP2pro:NLP2-GFP*, *NLP2pro:mCherry-NLP2* and *UBQ10pro:NLP2-GFP* constructs. The 2-kb 5’ of the NLP2 start codon and the full-length *AtNLP2* cDNA were amplified by PCR or RT-PCR from DNA or total RNA extracted from roots, rosettes and stems of *A. thaliana* using the primers as indicated in Supplemental Table S9 with high-fidelity Taq enzyme (Roche) and cloned into pGGA000 or pGGC000 GreenGate entry vector (Lampropoulos et al., 2013), respectively. The nucleotide sequence of the insert modules was checked before assembling into the pGGZ001 destination vector. Binary vectors were introduced into Agrobacterium (*Agrobacterium tumefaciens*) strain C58C1 (pMP90). The *nlp2-2* mutant was transformed by the *in-planta* method using the surfactant Silwet L-77, and transformants were selected on 50 μg/mL of Kanamycin.

Plants expressing the *NLP2pro:mCherry-NLP2* and *NLP7pro:GFP-NLP7* (Castaings et al., 2009) constructs were obtained by crosses and heterozygous seeds were used for confocal analysis.

### Growth conditions

For ChIP and Primary Nitrate Response assays, seeds were stratified for 3 days at 4°C and then grown in liquid medium containing 0.07% (w/v) MES pH 5,8, 1 mM KH_2_PO_4_, 0.5 mM MgCl_2_, 0.5 mM CaSO_4_, microelements as in (Estelle and Somerville, 1987), 0.25% (w/v) sucrose and 3 mM KNO_3_ for 10 days in long-day photoperiod (16- h light/8-h dark) at 25°C with a light intensity of 80 µmol photons.m^−2.^s^−1^. After 7 days of growth, the medium was changed for fresh medium each 2 days. Eleven-day-old seedlings were then nitrogen-starved for 3 days using fresh medium containing 3 mM KCl instead of 3 mM KNO_3_. After 3 days of N starvation, 3 mM nitrate was added for the indicated time periods and harvests were performed at several time points (0 min and 30 min for the ChIP assay and 0, 15, 30, 45 and 60 min after nitrate resupply for the PNR assays). Transcriptome analysis was performed on samples harvested at 0’ and 30’ after nitrate resupplying.

For the analyses of adult plants under steady state growth conditions, seeds were stratified for 3 days at 4°C in the dark and then grown on sand culture supplemented with nutrient solutions (Loudet et al., 2003) containing either 0.5 mM (limiting nitrate supply), 5 mM (non-limiting nitrate supply) nitrate, or 2.5 mM ammonium nitrate (non-limiting ammonium nitrate supply) in short day photoperiod with an 8-h light/16-h dark cycle at 21°C/18°C, respectively, 65% relative humidity, and a constant 150 μmol m^−2^ s^−1^ irradiation. For N limitation conditions, ion equilibrium of the medium was ensured by partly replacing KNO_3_ and Ca(NO_3_)_2_ by KCl and CaSO_4_. Forty-two-day-old plants were harvested at 3-4 h after onset of the light period for all analyses (transcriptome, metabolome, nitrate content, NR activity). The tissues were flash-frozen in liquid nitrogen and stored at −80°C until use.

### Chromatin immunoprecipitation

Seedlings were harvested and instantly frozen in liquid nitrogen. Chromatin extraction and immunoprecipitation was performed essentially as described in Bouyer et al., (2018) with minor modifications. Briefly, 800 mg of liquid nitrogen-ground seedlings were subject to double crosslinking (2.5 mM Di(N-succinimidyl) glutarate 60 min followed by 1% formaldehyde 7 min). Following nuclei purification, chromatin fragmentation was done using a Covaris S220 ultra-sonicator with the following parameters: 12 min with 5% duty cycle, 105W peak power and 200 cycles per burst. Chromatin immunoprecipitation was performed using 3 µg of anti-GFP antibodies (ab290, Abcam) pre-bound to proteinA/G-coupled magnetic beads (Dynabeads, Invitrogen).

### ChIP-sequencing and data analysis

For each biological replicate, 250 pg of immunoprecipitated (IP) or genomic (INPUT) DNA were used to prepare libraries with the MicroPlex Library Preparation kit v2 (Diagenode). Quality of libraries was validated using Agilent / Advanced Analytical fragment analyzer. Multiplexed libraries were sequenced using a NextSeq 500 system (Illumina) with paired-ends 41 bp reads. Reads were mapped to the tair10 genome assembly using Bowtie2 v2.3.4.1 with --very-sensitive -p 10 parameters and allowing one mismatch. Duplicate flagging was done using PicardTools MarkDuplicates v2.8.1 and filtering out genomic regions with aberrant coverage or low sequence complexity (Quadrana et al., 2016) was done using samtools v1.7 to remove sequencing biases. Peak-calling was done using MACS2 v2.1.1 in narrow mode, format BAMPE, with genome size set to 10e7, --keep-dup parameter set to 5, and using the -B option. Irreproducibility Discovery Rate (IDR, (Li et al., 2011)) was used to measure consistency between biological replicates by statistically determining the reproducibility in score ranking between peaks in each replicate. IDR analysis code is available at https://github.com/mlin/encode2/tree/master/chip-seq2/idr/resources/opt/idrCode. Following ENCODE guidelines (Landt et al., 2012), each biological replicate is subject to peak calling and IDR analysis is performed with a 5% threshold. It was verified that the number of bound regions identified between replicates was greater than 50% of the number of regions identified between two “pseudoreplicates” generated by pooling and then randomly partitioning all available reads from all replicates. To ensure similar weighting of individual replicates for identifying binding regions, it was also verified that the number of significant peaks identified using IDR on each individual replicate (obtained by partitioning reads into two equal groups for the IDR analysis) was within a factor of 2 of one another. The number of consistent peaks was considered as the highest number N between the number of reproducible peaks obtained by comparing true replicates and pooled pseudo replicates. To generate the list of reproducible peaks, peak-calling was performed on the alignment resulting from pooling all reads from the different replicates and the top N peaks were considered. Peaks overlapping regions showing aberrant enrichment in the INPUT sample as detected by MACS2 were removed from the final list. ChIP-seq peaks were assigned to an annotated gene as follows: if a peak overlaps completely, partially or is included in a gene, it is associated to this gene. If no overlap is found, the peak is assigned to the nearest annotated gene within a 5 kb window. In rare cases where a peak is equally distant from two genes, the two annotations are kept. The Integrative Genomics Viewer (Robinson et al., 2011) was used to visualize ChIP-seq signal distribution over representative genes.

Binding in response to nitrate was corroborated by selecting for further analyses peaks that were detected specifically at 30 minutes. All peaks were analyzed visually and corrected (106 peaks detected in the input have been removed, and 8 false positif peaks detected by MACS2 suppressed).

### DNA binding motif search

We used MEME (Bailey et al., 2015) to search for enriched DNA motifs in the 200 bp centered on NLP2 peak summits with -mod zoops -dna -revcomp parameters. We used HOMER (Heinz et al., 2010) to find the AARRGNCA and the AARRG DNA motif in the 3’ distal region to the TGNCYYTT motif identified by MEME.

### Gene expression analysis

For genome-wide early nitrate response analysis, three independent biological replicates were produced. Col-0, *nlp2-3* and *nlp7-1* before and after nitrate treatment were collected (plants at 1.02 developmental growth stages (Boyes et al., 2001) and cultivated as described above). Each sample is composed of 15 seedlings. Total RNA was extracted using the RNeasy plant minikit (QIAGEN) including a DNase treatment according to manufacturer’s instructions and was further purified using the RNA Clean & Concentrator Kits (Zymo Research®, California, U.S.A.). RNA-seq libraries were constructed by the POPS platform (IPS2) using the TruSeq Stranded mRNA library prep kit (Illumina®, California, U.S.A.) according to the supplier’s instructions. The libraries were sequenced in single-end (SE) mode with 75 bases for each read on a NextSeq500 to generate between 15.2 and 26.5 millions of SE reads per sample.

Adapter sequences and bases with a Q-Score below 20 were trimmed out from reads using Trimmomatic (v0.36, (Bolger et al., 2014)) and reads shorter than 30 bases after trimming were discarded. Reads corresponding to rRNA sequences were removed using sortMeRNA (v2.1, (Kopylova et al., 2012)) against the silva-bac-16s-id90, silva-bac-23s-id98, silva-euk-18s-id95 and silva-euk-28s-id98 databases.

Filtered reads were then mapped using Bowtie version 2 (Langmead and Salzberg, 2012) with the --local option against the Arabidopsis thaliana transcriptome. The 33602 genes were extracted from TAIR(v10) database with one isoform per gene (corresponding to the longest CDS). The reads were counted using sam2count. Between 96.97 and 98.53 % of the reads were associated with transcripts.

Next, genes with less than 1 read per million in at least half of the samples were discarded. The resulting raw count matrix was fed into edgeR (Robinson et al., 2010) for differential expression testing by fitting a negative binomial generalized log-linear model (GLM) including a group factor and a replicate factor to the TMM-normalized read counts for each gene. The group variable is a concatenation of genotype (wild type or mutant) and treatment. We performed pairwise comparisons between groups. The distribution of the resulting p-values followed the quality criterion described by (Rigaill et al., 2018). Genes with an adjusted p-value (FDR, (Benjamini and Hochberg, 1995)) below 0.05 were considered as differentially expressed.

Transcriptomes from 7-week-old plants (Col-0, Col-4, *nlp2-2*, *nlp2-3* and *nlp2-3 nlp7-1*) grown under constant non-limiting nutrient supply, were analyzed using RNA-seq and RNA extracted from three independent biological replicates. Total RNA was extracted from root and rosette using the RNeasy plant minikit (QIAGEN) including a DNase treatment according to manufacturer’s instructions. Preparation of RNA library and transcriptome sequencing was conducted by Novogene Co., LTD (Beijing, China). Sequencing libraries were generated using NEBNext® UltraTM RNA Library Prep Kit for Illumina® (NEB, USA) following manufacturer’s recommendations and index codes were added to attribute sequences to each sample. The clustering of the index-coded samples was performed on a cBot Cluster Generation System using PE Cluster Kit cBot-HS (Illumina) according to the manufacturer’s instructions. After cluster generation, the library preparations were sequenced on an Illumina platform and paired-end reads were generated. After standard quality control of raw reads, paired-end clean reads were mapped to the reference genome (TAIR10) using HISAT2 software. HTSeq was used to count the read numbers mapped of each gene and then RPKM of each gene was calculated based on the length of the gene and reads count mapped to this gene. Differential expression analysis between two conditions/groups was performed using DESeq2 R package. |log2(FoldChange)| > 0.4 and adjusted p-value (FDR) < 0.05 were set as the threshold for significantly differential expression.

For expression analyses via RT-qPCR, cDNAs were first checked by PCR for contamination by genomic DNA, then relative expression was determined in 384-well plates with a QuantStudio 5 System device (40 3-step cycles; step 1, 95°C, 5 s; step 2, 58°C, 30 s; step 3, 72°C, 30 s) using Takyon qPCR kit for SYBR assay (Eurogentec) (2.5 µL TAKYON SYBER 2X, 0.015 µL Forward primer 100 µM, 0.015 Reverse primer 100 µM, 2.5 µL cDNA 1/20 per well). Relative expression was calculated according to the 2^-ΔCt^ method. Target gene expression was normalized to the geometric mean of the expression of two plant genes either *UBIQUITIN10* (At4g05320) & *ACTIN2* (At3g18780) or *TIP41 (*At4g34270) and *ACTIN2* All primer sequences are given in Supplemental Table S9.

### GO term analysis

GO terms were defined using the classification superviewer tool at http://bar.utoronto.ca/ntools/cgi-bin/ntools_classification_superviewer.cgi using Mapman bins and sorted on frequency related to Arabidopsis set. Associated P-values of the hypergeometric distribution are indicated, which were calculated as follows: P=BC(M, x) × BC(N−M, n−x)/BC(N, n). BC is the binomial coefficient, calculated as follows: BC(n, k)=n!/(k! × [n−k]!). In the above equations, x is the number of input genes with the selected classification, n is the total number of input genes, M is the number of genes with the selected classification in the database (MapMan) and N is the total number of genes in this database

### Nitrate and nitrate reductase activity measurements

After ethanolic extraction of flash-frozen and then finely ground plant material, nitrate contents were determined in rosette and root samples (Orsel et al., 2004). A further aliquot of the same frozen plant material was used for determination of the maximum *in vitro* activities of nitrate reductase (Ferrario-Méry et al., 1998) and for GC-MS and LC-MS/MS metabolite profiling.

### Metabolite profiling

For GC-MS based quantifications, approximately 25 mg of finely ground plant material were resuspended in 1 mL of frozen (−20°C) Water:Acetonitrile:Isopropanol (2:3:3, v/v/v) containing Ribitol at 4 μg/mL and analyzed as in Forzani et al.(2019).

For LC–MS/MS-based quantifications, 20 mg of finely ground plant material was quenched in a 3:7 (v/v) chloroform: methanol mixture at liquid nitrogen temperature, and allowed to warm up to –20°C for 2 h. The chloroform phase was then extracted twice with ice-cold ultra-pure water. The two water phases were pooled together and evaporated to dryness in a centrifugal vacuum drier at 20°C. Pellets were resuspended in 350 mL of ultra-pure water, filtered through a Multiscreen Ultracel-10 (Milipore) membrane to remove high molecular weight components, and diluted 1:10 before analysis by high-performance anion-exchange chromatography coupled to tandem mass spectrometry, as described in (Lunn et al., 2006) with modifications (Figueroa and Lunn, 2016).

### Statistical analysis

Except for genomic analyses, the statistical analyses were performed using the rcompanion package in the R environment.

### Confocal Microscopy Analysis

Seedlings were grown for 5 days on the 3 mM KNO_3_ liquid medium (see above), nitrogen-starved for 2 days by exchanging the culture medium with a nitrogen-free medium, in which KNO_3_ was replaced by KCl. Plants were mounted in 40 μL of a top agar (0.1%, w/v) solution. During confocal imaging, either 5 μL of 0.5 M KNO_3_, 5 μL of 0.5 M NH4Cl, were added to the mounting medium; the same plantlets were imaged during a time course. For the nuclear export inhibition experiment, a 3-h preincubation with LMB (Sigma, at 100 nM final concentration) was performed in N-free medium. Laser scanning confocal imaging was performed using the Zeiss 710 or the Leica SP8 microscope equipped with an argon laser (488 nm for green fluorescent protein (GFP) excitation and 561 nm for mCherry). Emission was collected at 510 to 545 nm (GFP) and at 600 to 630 nm (mCherry). A sequential scanning mode was used when mCherry was combined with GFP to minimize the crosstalk between the two partially overlapping emission spectra. The images were coded green for GFP and magenta for mCherry giving white colocalization in merged images. Images were processed in ImageJ and Photoshop Element (Adobe Systems). Each image shown represents a single focal plane.

### Bimolecular Fluorescence complementation (BiFC)

Open reading frames (without stop codon) flanked by AttB1 and AttB2 sites were amplified from Arabidopsis cDNA clones (Columbia accession), cloned into Gateway vector pDONR207 using BP recombination (Invitrogen), and sequenced. An LR reaction between the entry vector and the complete set of four pBiFP vectors (Azimzadeh et al., 2008) produced the final expression vectors, where coding sequences are cloned in fusion with the N- and C-terminal parts of YFP, either as N- terminal or C-terminal fusions, under the control of the cauliflower mosaic virus 35S promoter. For BiFC assays, the vectors were transformed into Agrobacterium strain C58C1(pMP90) and co-infiltrated into *Nicotiana benthamiana* leaves of five-week-old plants that have been grown under 2mM nitrate nutrition and then starved for N for 10 days. Agrobacterium carrying clones of interest were grown overnight at 28°C in 5 mL LB medium with appropriate antibiotics. Agrobacterium cells were collected by centrifugation and resuspended in 10 mM MgCl_2_ and 1 mM 2-(N-morpholine)- ethanesulphonic acid (MES), pH 5.6, to obtain a final OD 600nm of 0.5 for *N. benthamiana* leaf infiltration. YFP fluorescence was detected 2 days after infiltration. Observations were carried out using a ZEISS 710 Anisotropy 405 confocal laser microscope. YFP fluorescence was recorded after an excitation at 514 nm (Argon laser) and a selective emission band of 520–540 nm. Chlorophyll autofluorescence was excited by the Argon laser (514 nm) and recorded with a selective emission band of 650–700 nm.

## Supplemental Data

Supplemental Table S1-10.

**Supplemental Figure S1.**
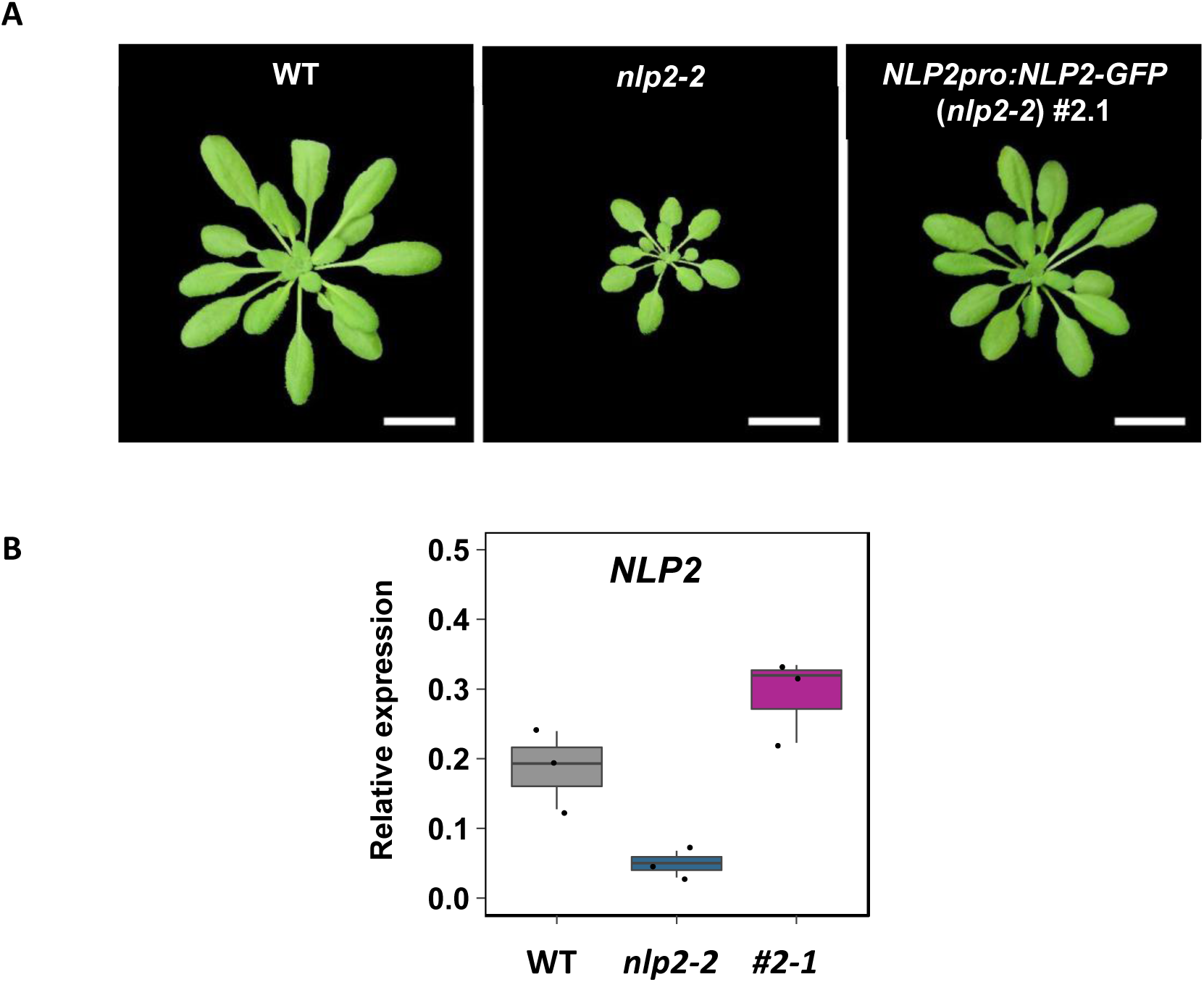
Complementation of *nlp2-2* mutant expressing *NLP2pro*- driven NLP2-GFP fusion protein (#2.1). **(A)** Top view of 42-d-old rosettes from WT, *nlp2-2* and *nlp2-*2 line expressing *NLP2pro*- driven NLP2-GFP fusion protein grown under non-limiting (5 mM) nitrate supply. **(B)** Relative expression of *NLP2* measured in 14-d-old seedlings. Data are presented as boxplots and each value is shown as black point.

**Supplemental Figure S2.**
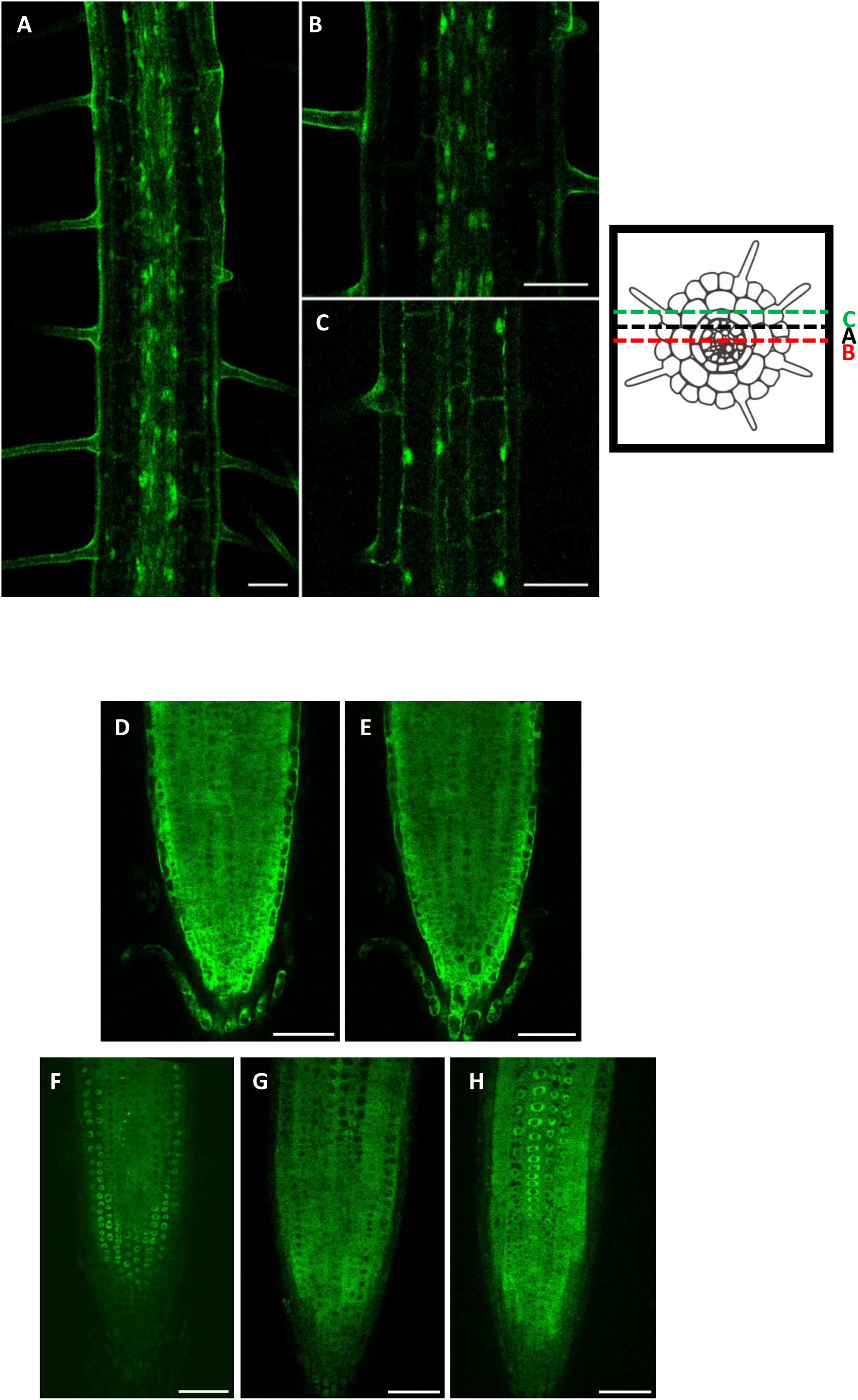
Characterization of the nucleocytosolic shuttling and spatial distribution of NLP2 along the root. **(A to F)** Confocal imaging was performed on 14-d-old *nlp2-2* seedlings expressing *NLP2pro:NLP2-GFP.* (**A-C)** Z sections of the elongation zone highlighting pNLP2:NLP2-GFP signal in stele **(A, B)** and cortex (**C**) cells in the presence of nitrate. (**D, E**). Root tip of N-starved seedlings before (D) after 10 min ammonium resupply (E), (**F**) Root tip of N-starved seedlings after 3h of Leptomycin B treatment, an inhibitor of active nuclear export. (**G, H**) Confocal imaging was performed on *nlp2-2* seedlings expressing *pUbi10:NLP2-GFP*. Scale bar = 50 µm.

**Supplemental Figure S3.**
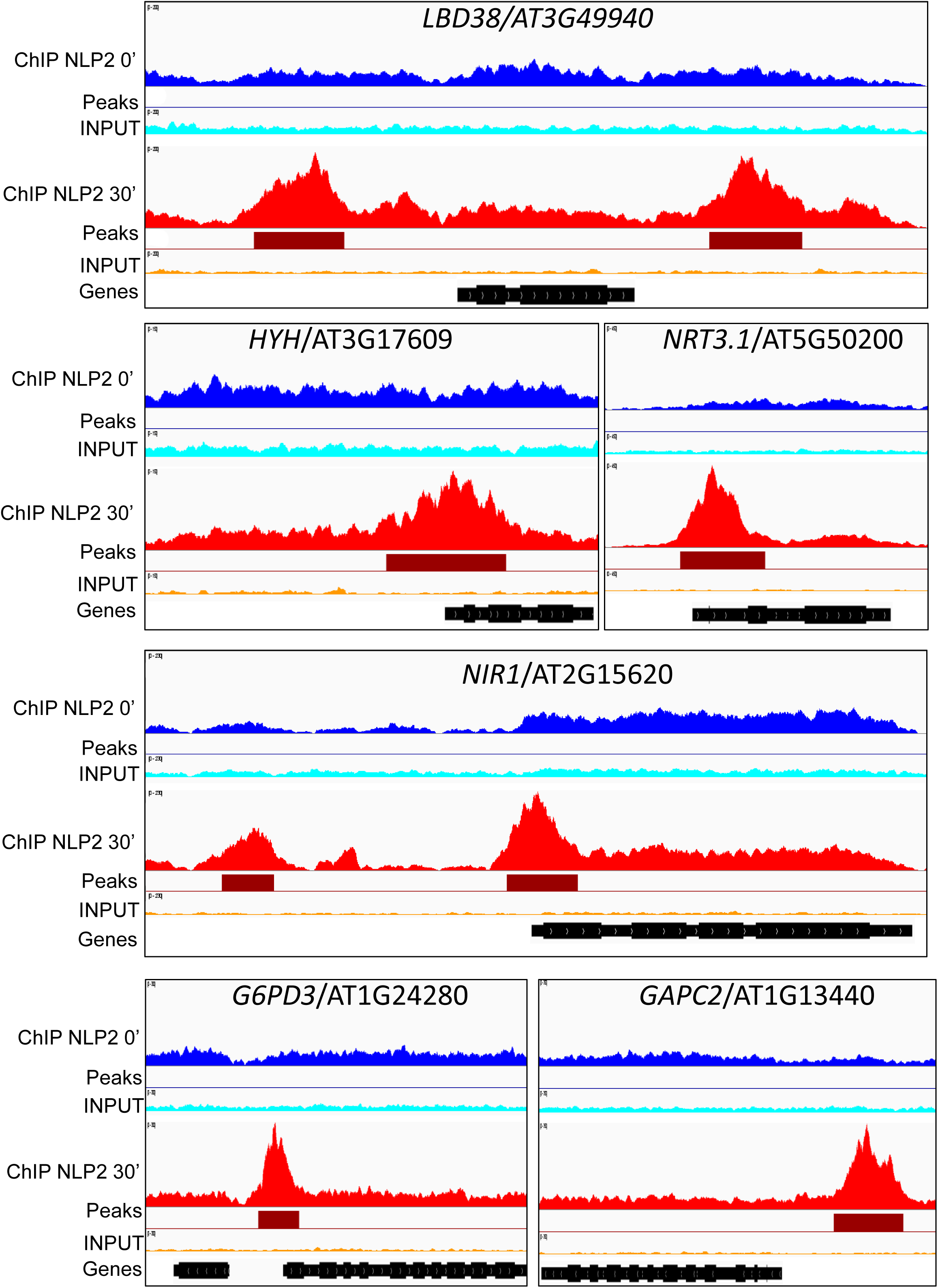
Integrative Genomics Viewer (IGV) visualization of representative NLP2-ChIP-seq results. For each locus, the ChIP-seq signal of pooled replicates obtained from samples before (T0, blue) and 30 min after nitrate addition (T30, red) is shown together with the INPUT (control, turquoise) signal. IDR-selected peaks identified by MACS2 are shown in dark red boxes. Note that all tracks within a given window are at the same scale.

**Supplemental Figure S4.**
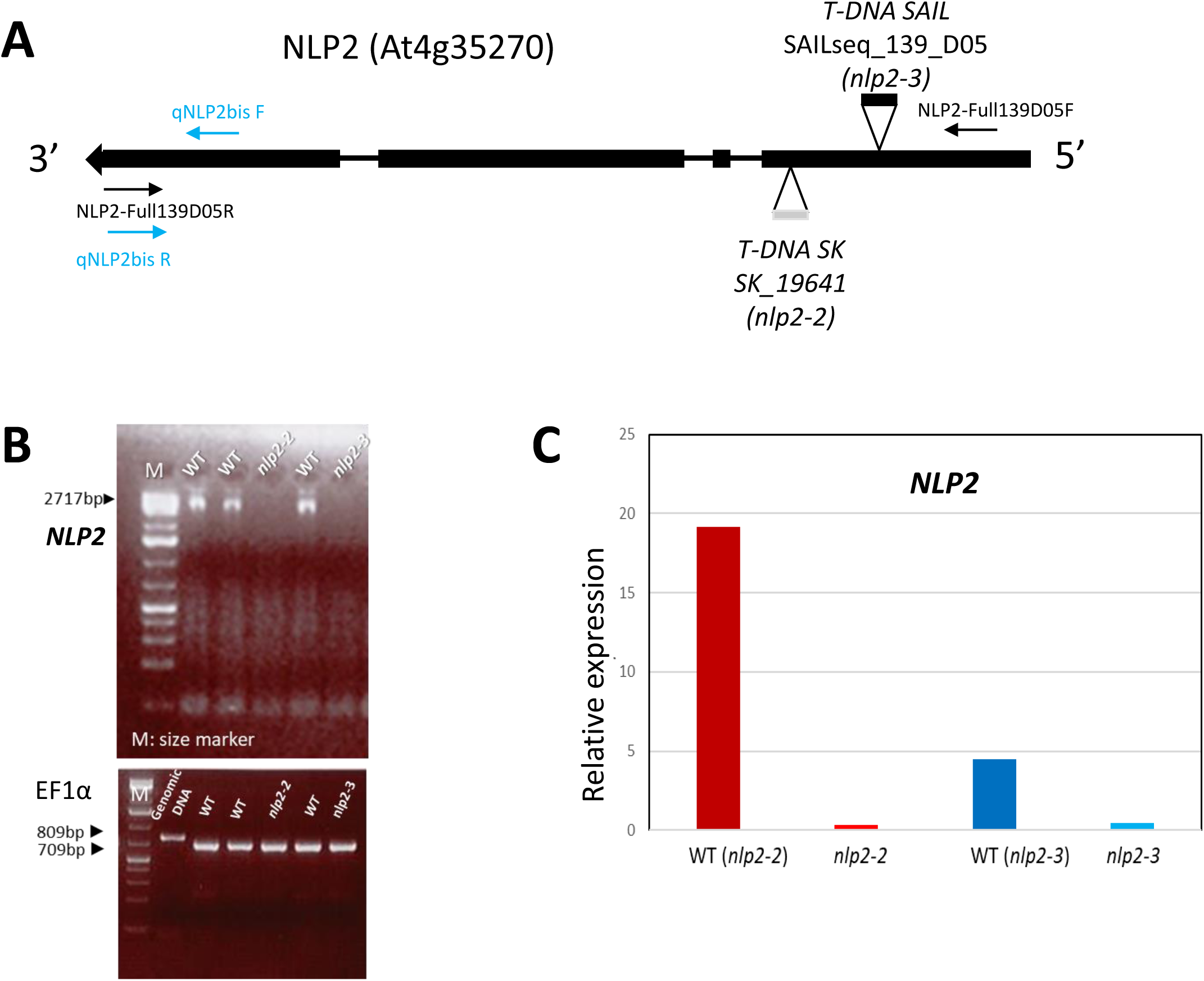
Molecular characterization of *NLP2* T-DNA insertion mutants. **(A)** Schematic map of the T-DNA insertion sites in *nlp2-2* and *nlp2-3* mutants. Black boxes and black lines represent exons and introns, respectively. Primers used for RT-PCR and RT-qPCR are indicated in black and blue, respectively and sequences are given in Supplemental Table S9. **(B,C)** *NLP2* expression in 21-day-old seedlings grown on soil in the greenhouse. **(B)** RT-PCR, *EF1α* was used as internal control. Both *NLP2* and *EF1α* were amplified for 35 cycles. **(C)** RT-qPCR analysis with *NLP2* expression levels represented as % of the expression of a constitutive synthetic gene composed of *TIP41* (At4g34270) and *ACTIN2* (*ACT2*, At3g18780).

**Supplemental Figure S5.**
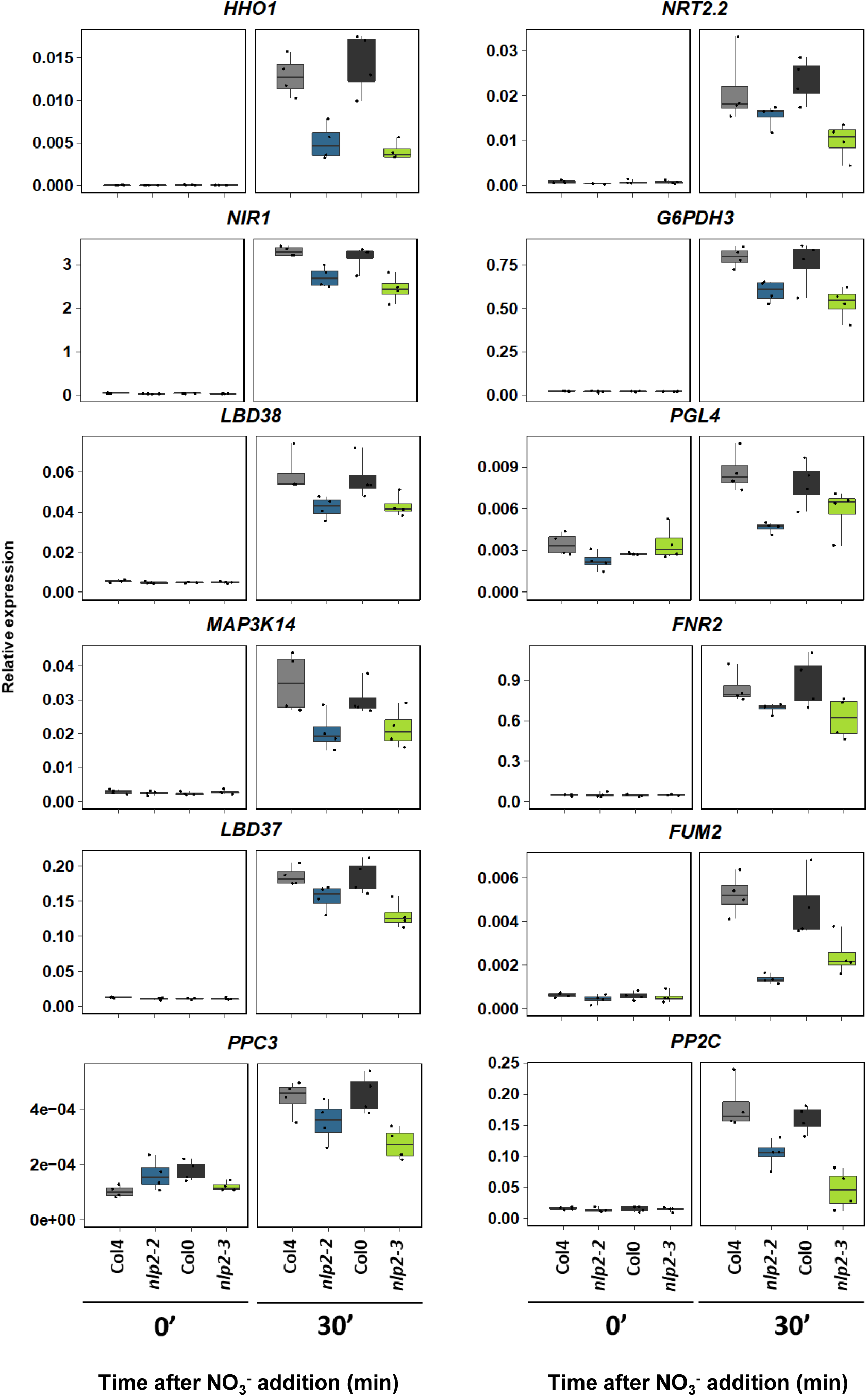
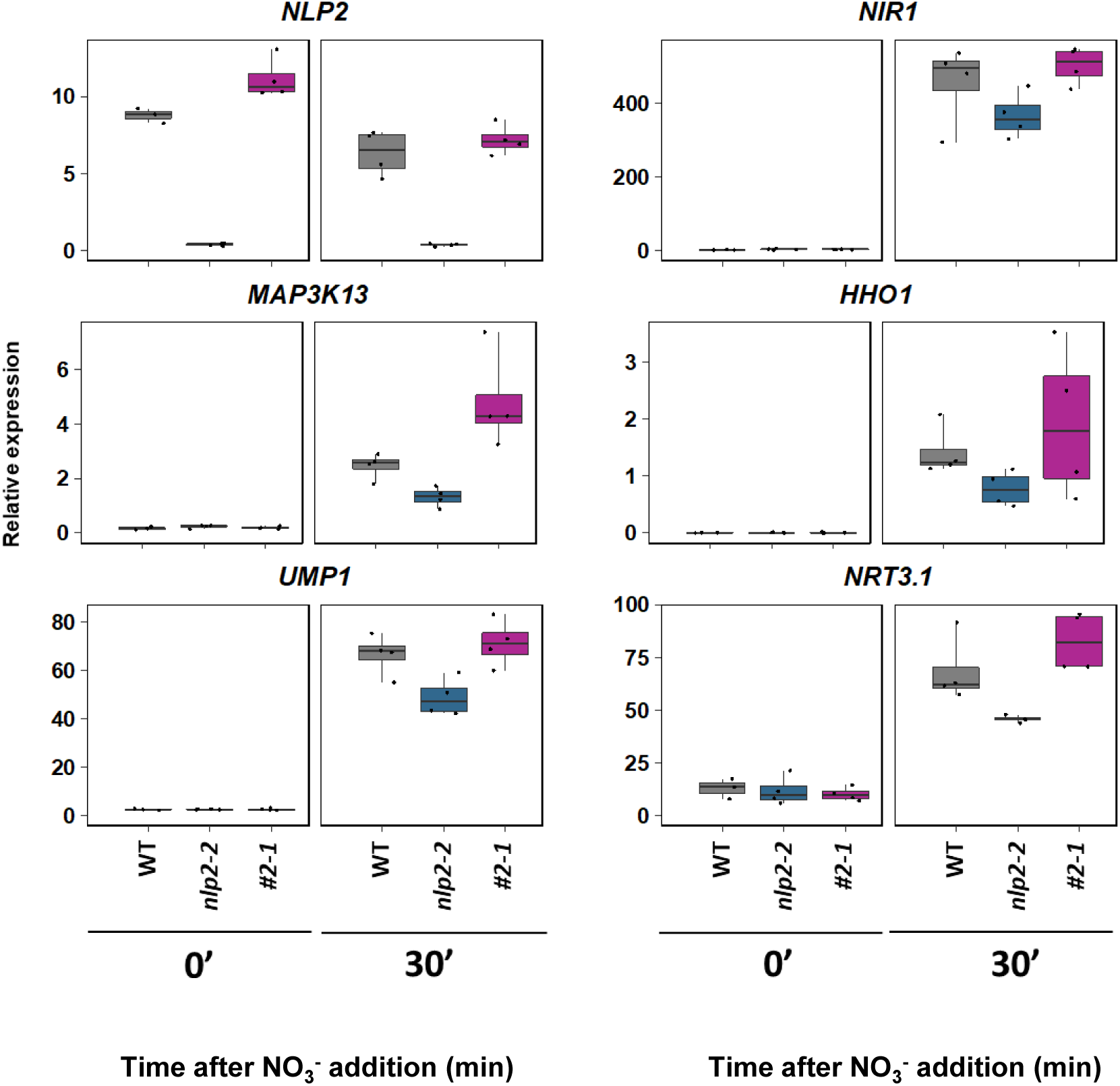
RT-qPCR validation of *nlp2-2* DEGs as identified by transcriptomics. **(A)** Gene expression before and after 30 min of nitrate supply to N depleted seedlings in the nlp2-2, *nlp2-3* and wild-type background. Gene expression is indicated as % of the expression of a constitutive synthetic gene composed of *UBIQUITIN10* and *ACTIN2*. **(B)** Gene expression before and after 30 minutes of nitrate supply to N depleted seedlings in *nlp2-2, NLP2pro:NLP2-GFP (nlp2-2) #2.1* and wild-type background. Gene expression is indicated as % of the expression of a constitutive synthetic gene composed of *TIP41* and *ACTIN2*. A, B, Plants were grown and samples were treated as in Figure 3. Data of one representative experiment are presented as boxplots and each value is shown as black point.

**Supplemental Figure S6.**
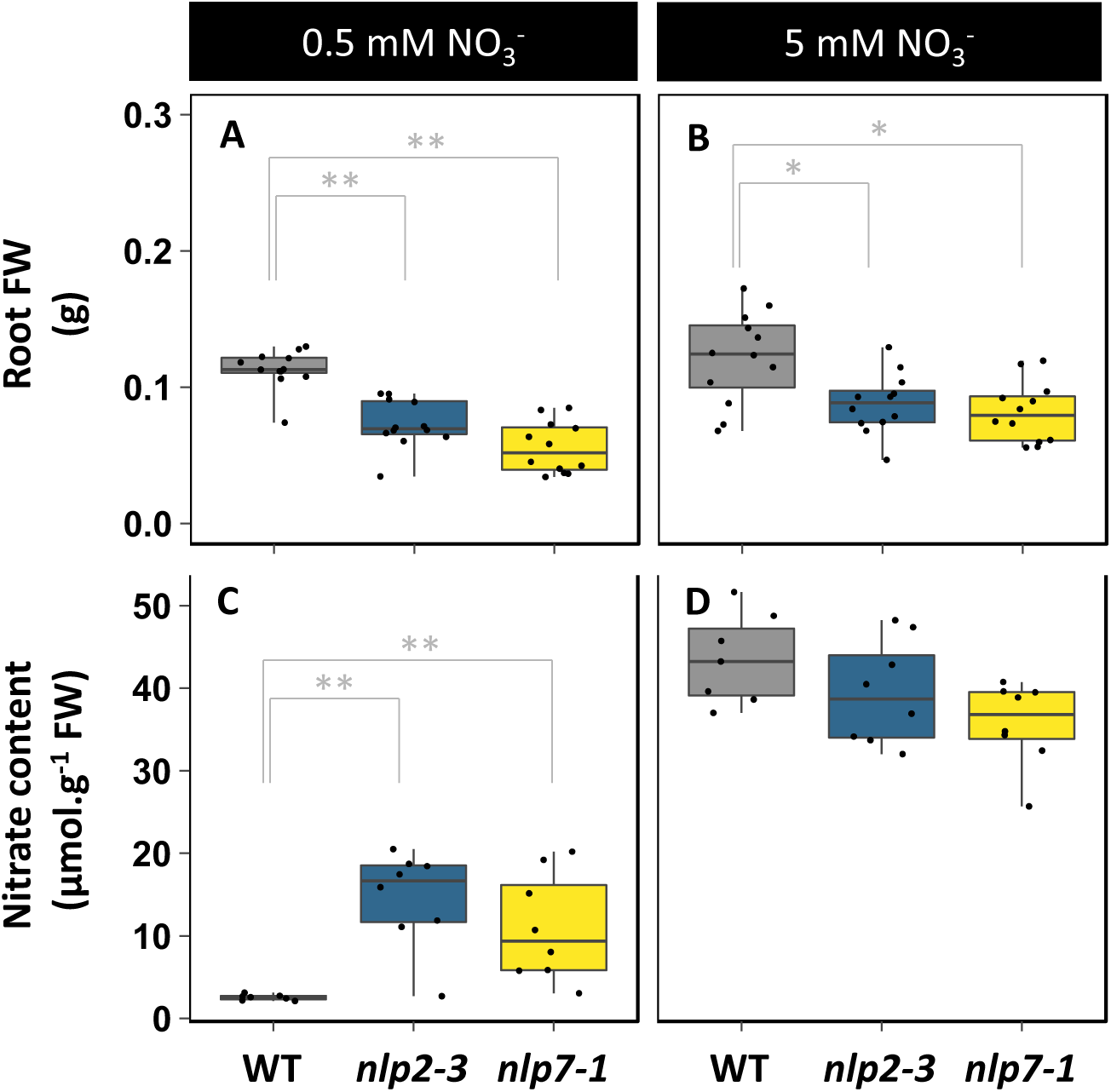
Biomass and nitrate content of roots from *nlp2-3* and *nlp7- 1* mutants grown under limiting and non-limiting nitrate supply. **(A-B)** Root fresh weight of WT, *nlp2-3* and *nlp7-1* grown under limiting (0.5 mM, A) and non-limiting (5 mM, B) nitrate supply. (**C-D**) Root nitrate content of WT, *nlp2-3* and *nlp7-1* grown under limiting (0.5 mM, C) and non-limiting (5 mM, D) nitrate supply. For A to D, data are presented as boxplots and each value is shown as a black point. For A and B, data are obtained from six plants per experiment and two independent experiments. For C and D, data are obtained from four plants per experiment from two independent experiments.

**Supplemental Figure S7.**
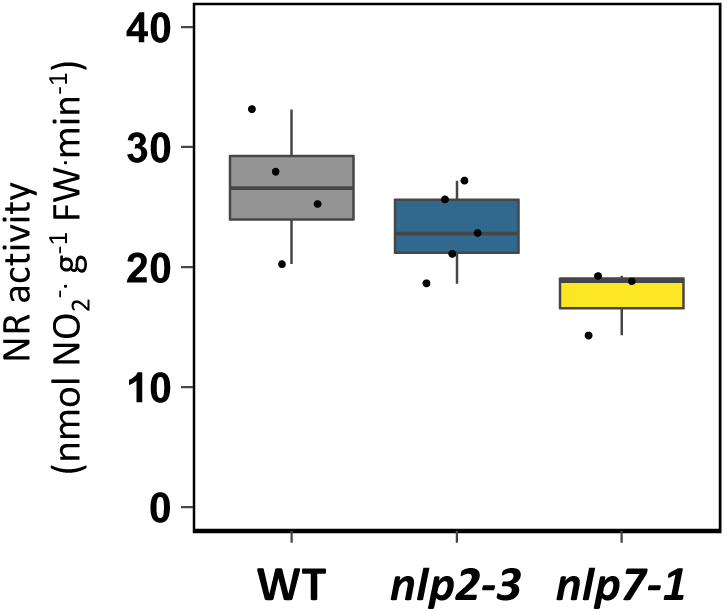
Biomass and nitrate content of *nlp2-2* mutant and *nlp2-2* complemented line (#2.1) harboring *NLP2pro:NLP2-GFP*, grown under limiting and non-limiting nitrate supply. Plants were grown on sand in short days with a constant light condition of 150 µE. **(A)** Top view of 42-d-old rosettes grown under limiting (0.5 mM) and non-limiting (5 mM) nitrate supply, (**B-C**) Rosette fresh weight of plants grown under limiting (0.5 mM, B) and non-limiting (5 mM, C) nitrate supply. (**D-E**) Rosette nitrate content of plants grown under limiting (0.5 mM, D) and non-limiting (5 mM, E) nitrate supply. For B to E, data are presented as boxplots and each value is shown as black point. For B to C, data were obtained from six plants per experiment from two independent experiments. For D and E, data were obtained from four plants per experiment and two independent experiments. Significant differences between genotypes for each nitrate supply condition was determined by a pairwise permutation test followed by false discovery rate adjustment (* *P*-value ≤ 0.05, ** *P*-value ≤ 0.001).

**Supplemental Figure S8.**
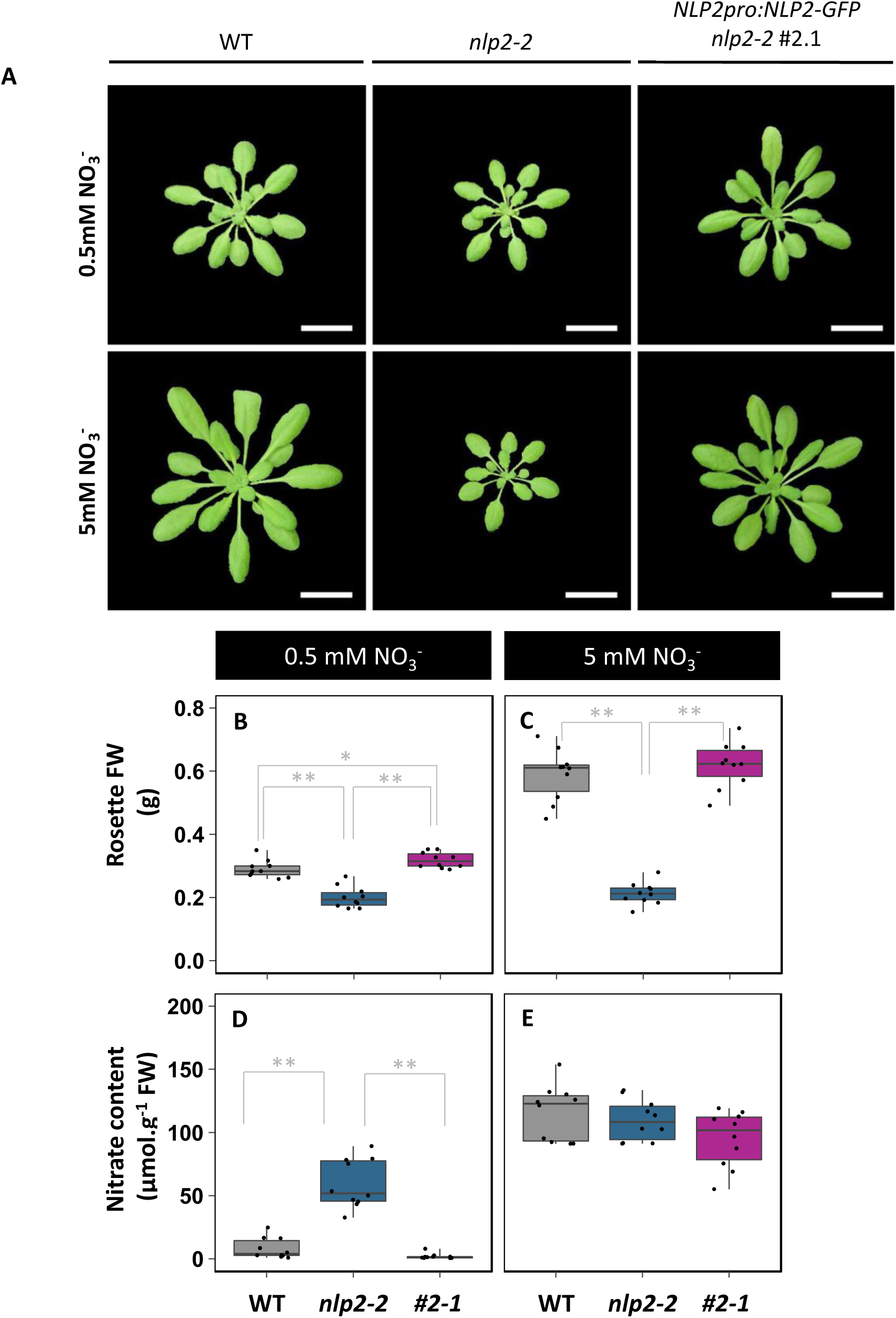
Nitrate reductase activity in rosettes of WT, *nlp2-3* and *nlp7-1* grown as in Figure 4. Data are presented as boxplots and each value is shown as black point. Data were obtained from three to four plants.

**Supplemental Figure S9.**
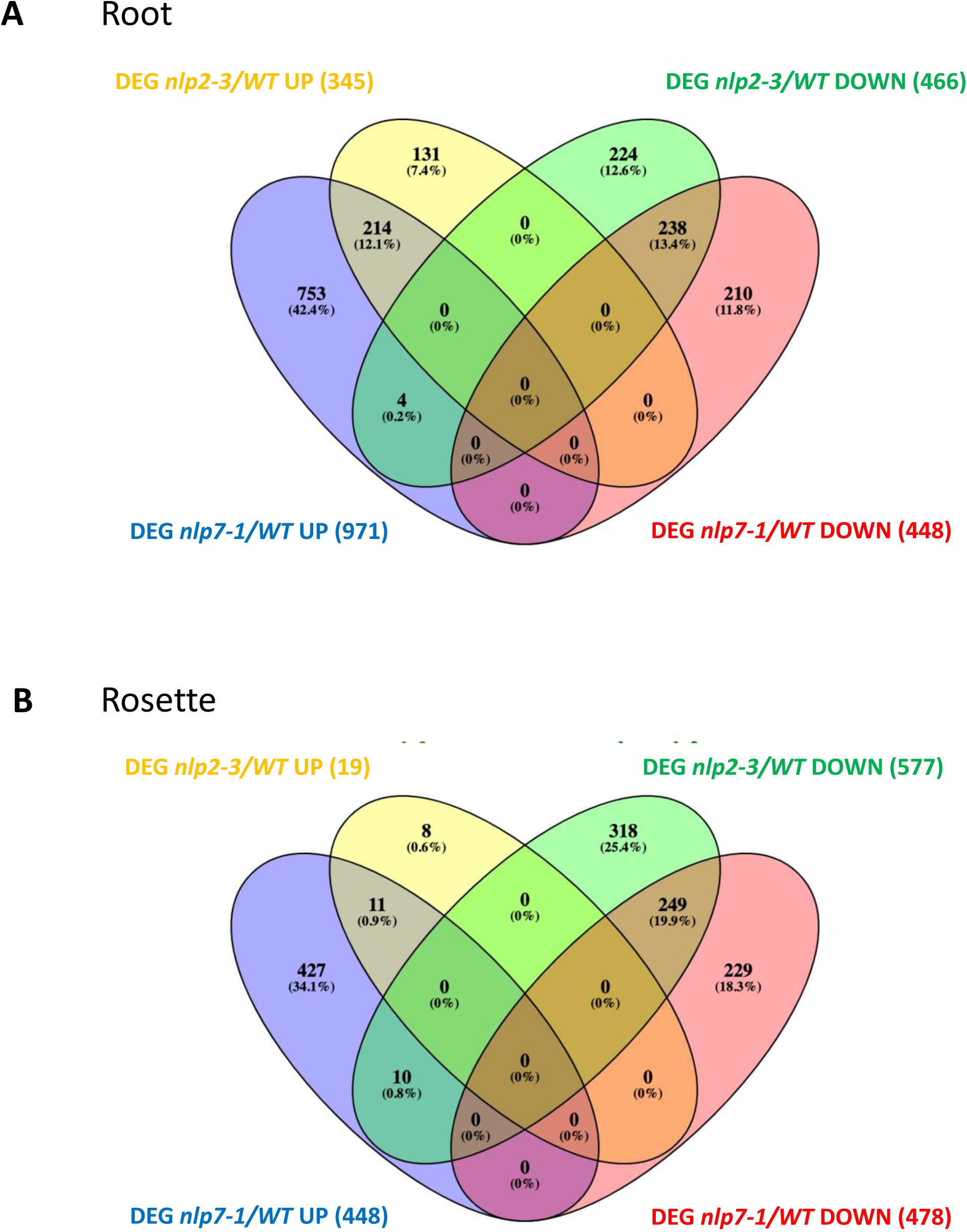
Venn Diagram summarizing DEGs compared to WT of *nlp2-3* and *nlp7-1* mutants grown under non-limiting nitrate supply as in Figure 4. Significant differences in gene expression were determined according to log_2_-fold changes (Log_2_-FC ≥ 0.4 or Log_2_-FC ≤ -0.4) with false discovery rate adjustment (*P*- value ≤ 0.05). **(A)** Roots. **(B)** Rosettes.

**Supplemental Figure S10.**
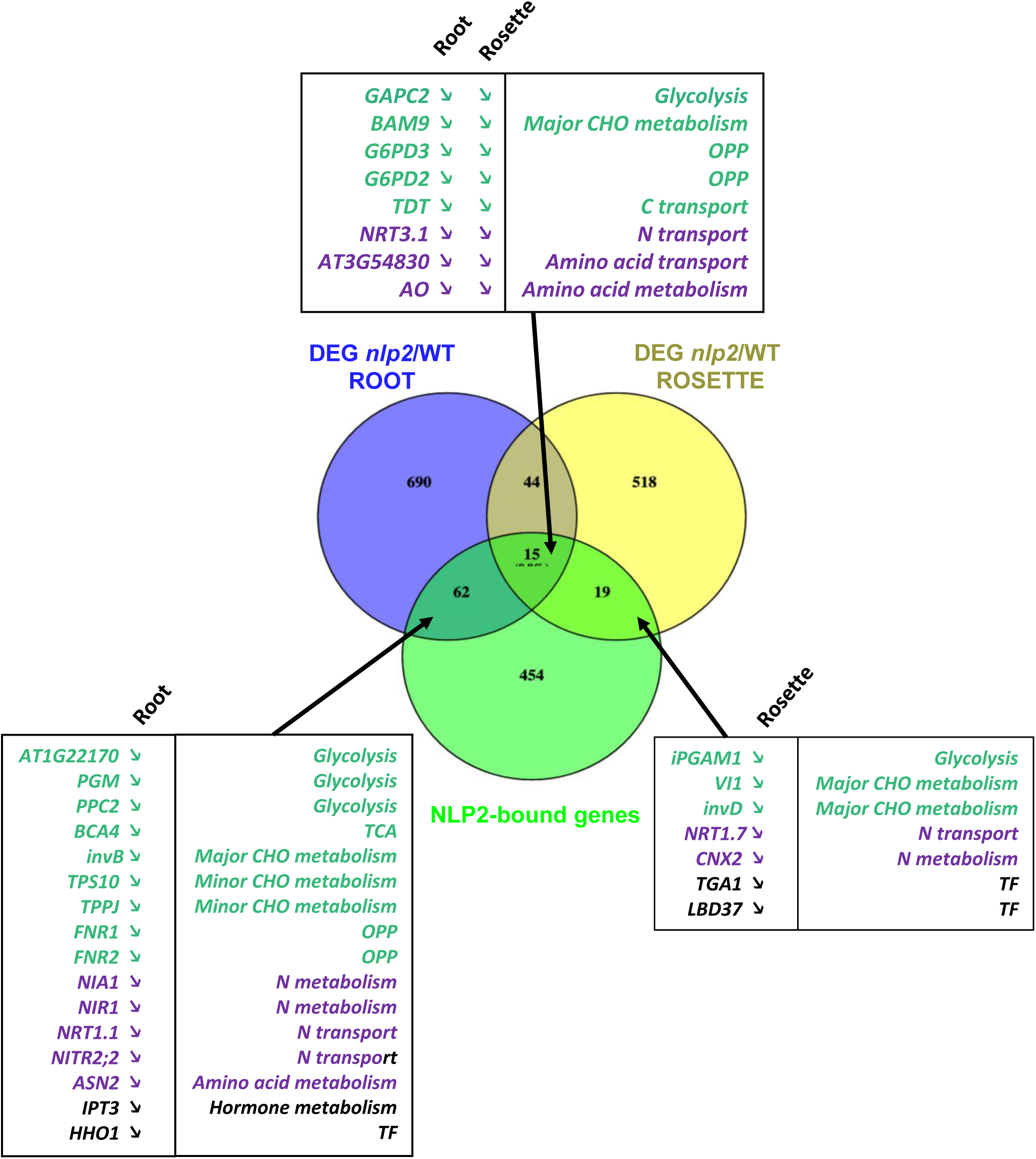
Overlap of NLP2-bound genes after 30 min of nitrate resupply in 14-d-old seedlings with DEGs in 42-d-old plants cultured under non-limiting nitrate supply as in Figure 4.

**Supplemental Figure S11.**
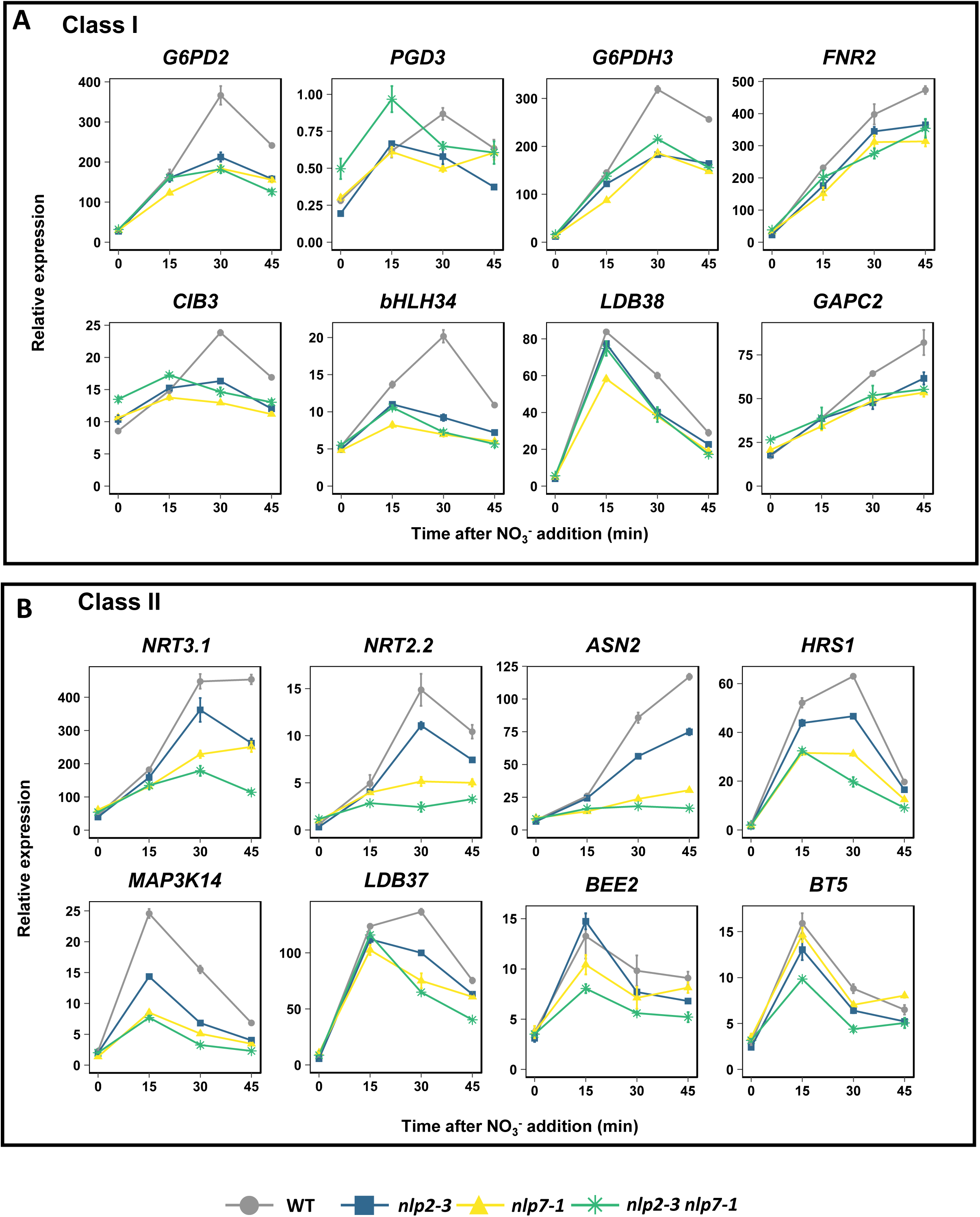
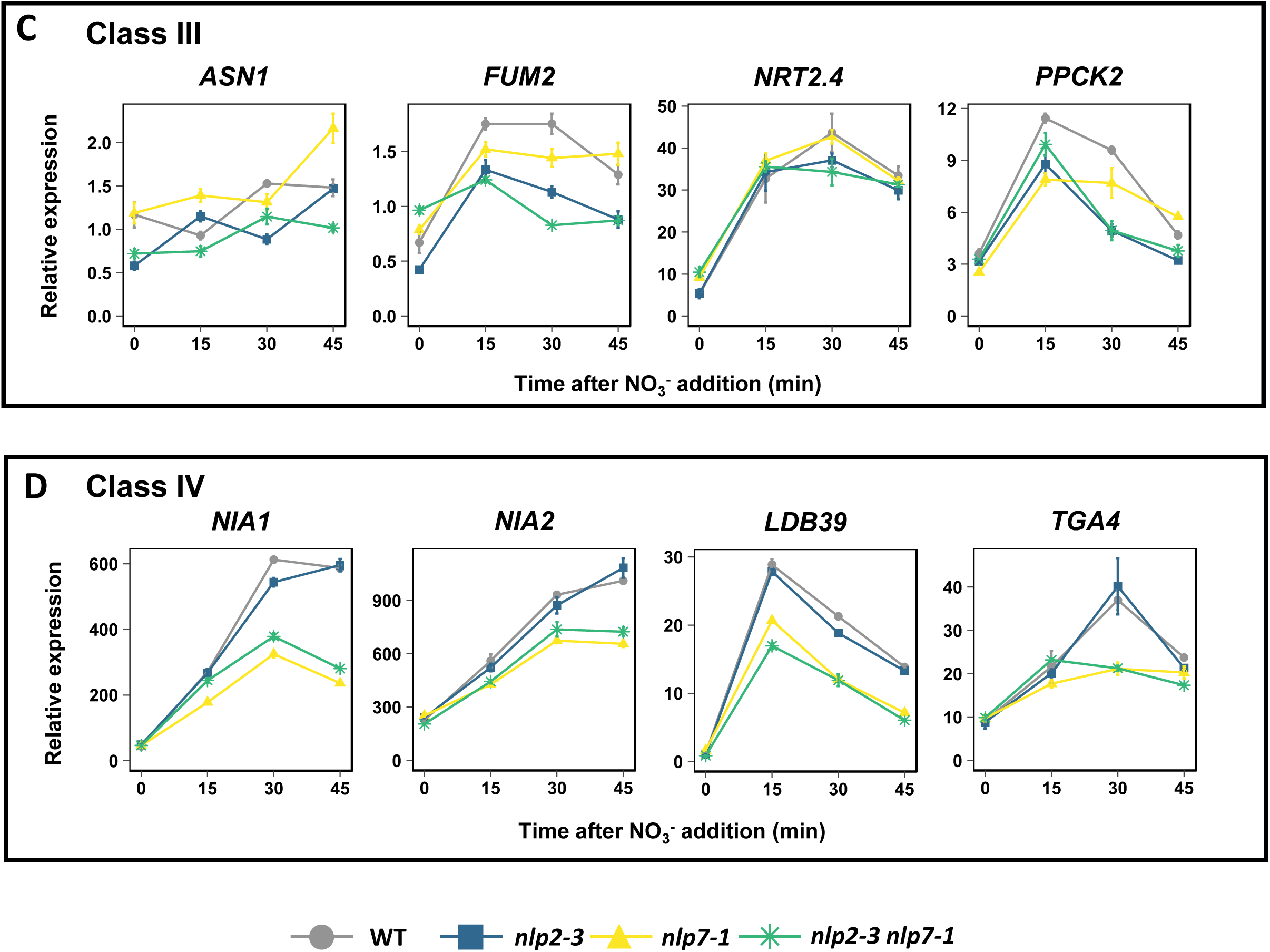
Representative examples of gene expression in *nlp2-3, nlp7-1* and *nlp2-3 nlp7-1* during early nitrate response. **(A-D)** Expression level of genes measured after nitrate addition on N-starved seedlings of WT (grey), *nlp2-3* (blue), *nlp7-1* (yellow) and *nlp2-3 nlp7-1* (green). Class I (double mutant showed no accentuated changes compared to single mutants), II (accentuated changes in double mutant), III (changes more related to NLP2 loss of function) and IV (changes more related to NLP7 loss of function) are shown in A, B, C and D, respectively. Gene expression was measured by RT-qPCR and expression indicated as % of the expression of a constitutive synthetic gene composed of *TIP41* (At4g34270) and *ACTIN2* (*ACT2*, At3g18780). Data are means ± SE (n=4) of one representative experiment.

**Supplemental Figure S12.**
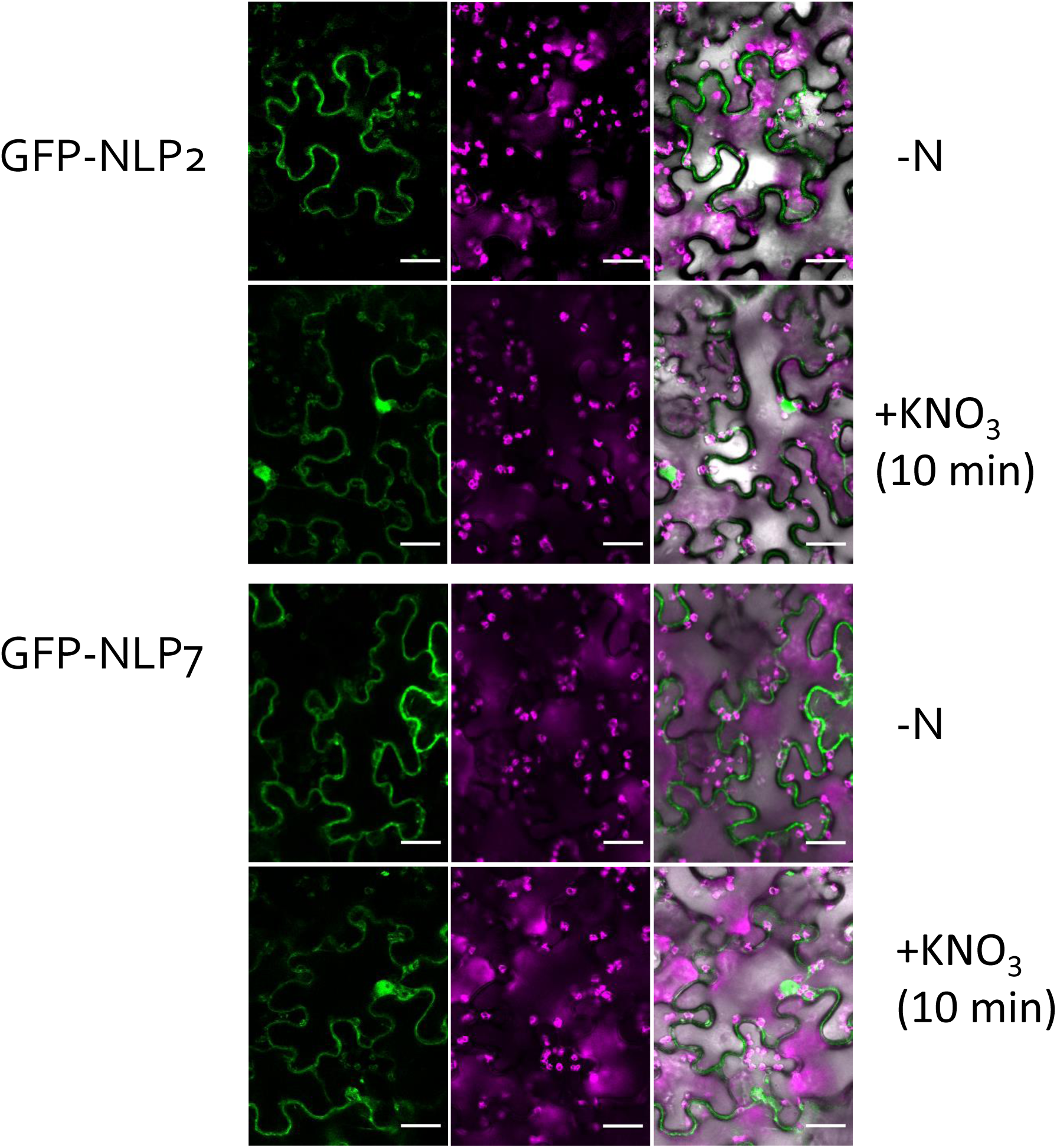
Transient expression of GFP-NLP2 and GFP-NLP7 in leaves of *Nicotiana benthamiana* in presence or absence of nitrate.

**Supplemental Figure S13.**
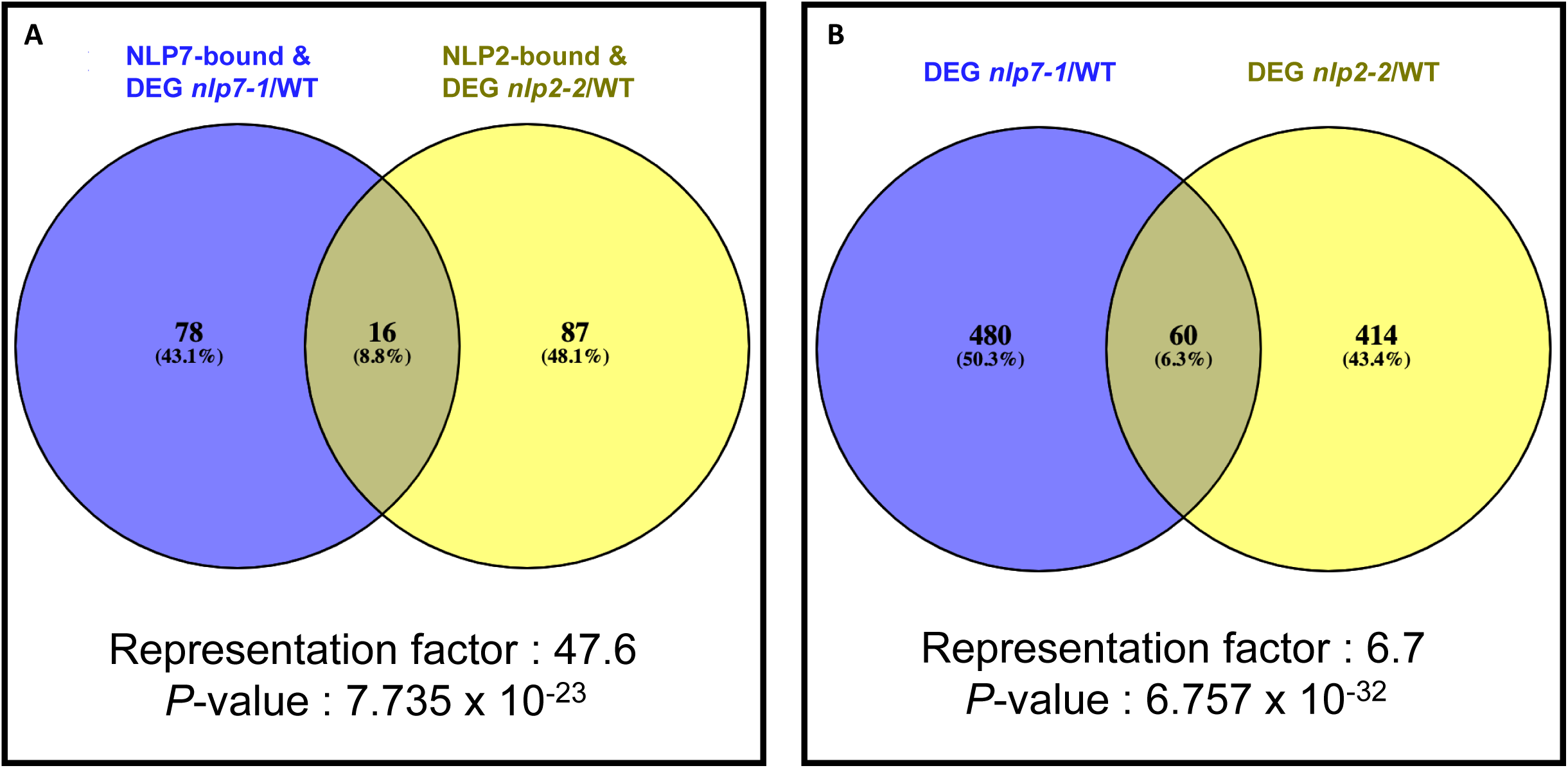
Comparison between NLP2 and NLP7 bound and differential expressed genes (DEGs). **(A)** Venn diagram showing the overlap between genes bound by NLP7 and differentially expressed *nlp7-1/WT* after 30 min of nitrate resupply with genes bound by NLP2 and differentially expressed *nlp2-2/WT* after 30 min of nitrate resupply *nlp2-2*. **(B)** Venn diagram showing the overlap between DEGs *nlp7-1/WT* with the DEGs *nlp2-2/WT*. NLP7 data are from Marchive et al. (2013).

**Supplemental Figure S14.**
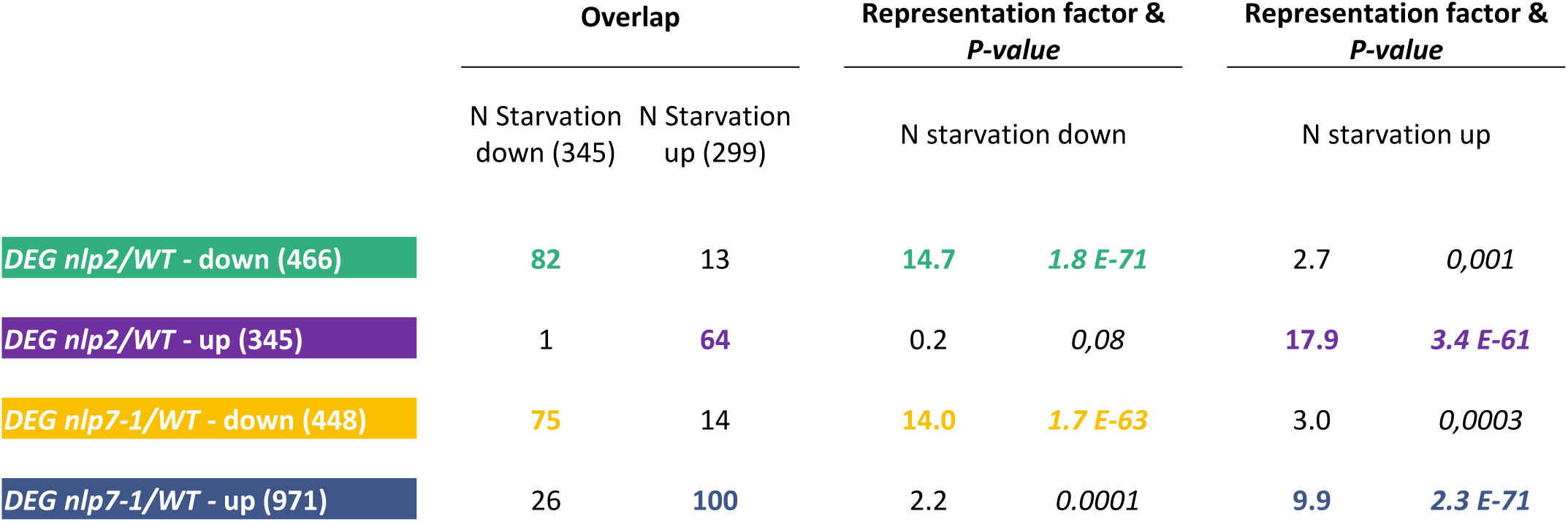
Comparison between DEGs *nlp2*/WTand *nlp7-1/WT*from roots of 42-d-old plants grown under non-limiting nitrate supply with N starvation-regulated genes (from Krapp et al., 2011). The overlap is given as numbers of genes, representation factors and *P*-values are presented.

**Supplemental Figure S15.**
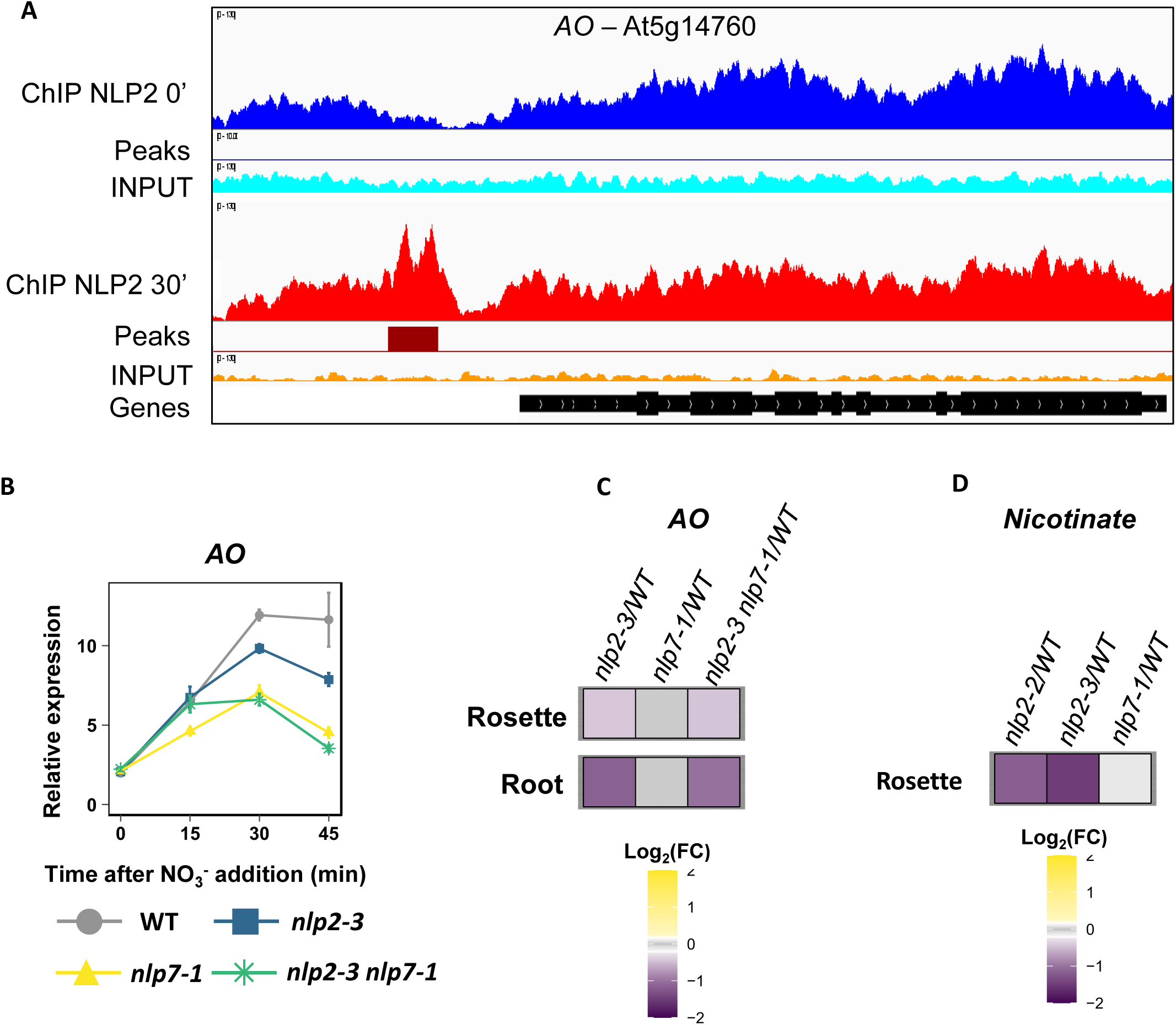
Focus on NAD metabolism related alterations in *nlp2* mutants. **(A)** The *AO* gene, encoding aspartate oxidase involved in the first step of NAD synthesis *de novo* pathway, is bound by NLP2 after nitrate resupply to 14-d-old N deprived seedlings . Tracks are as described in Supplemental Figure S3. **(B)** *AO* is differentially expressed in *nlp2-3*, *nlp7-1* and *nlp2-3 nlp7-1* compared to WT after nitrate addition to 14-d-old seedlings. **(C)** *AO* is differentially expressed in rosette and root of *nlp2-3* and *nlp2-3 nlp7-1* mutants grown under non-limiting nitrate supply. **(D)** The content in nicotinate, a precursor of NAD synthesis, is specifically reduced in rosettes of *nlp2* mutants grown under non-limiting nitrate supply.

**Supplemental Figure S16.**
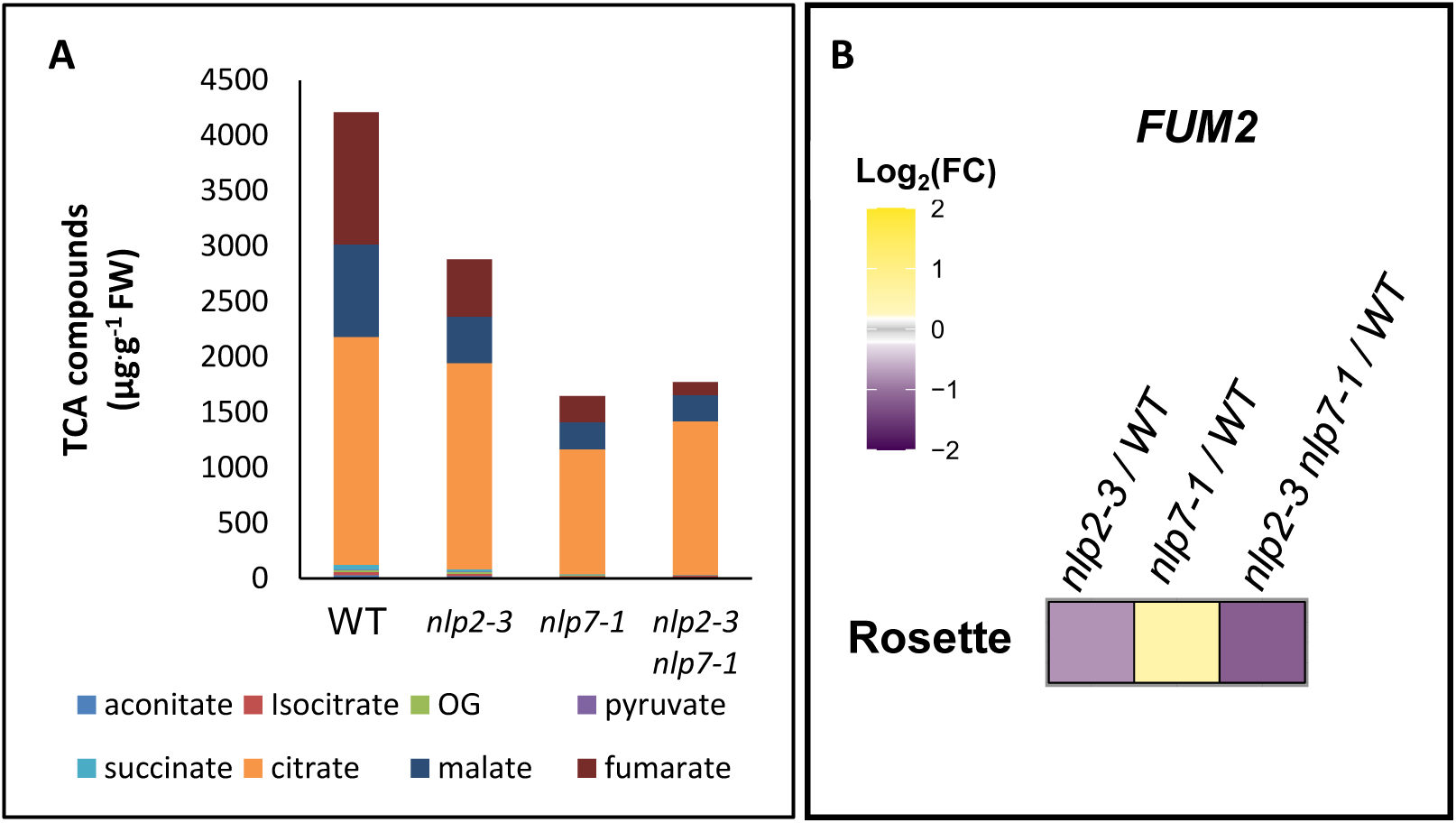
Focus on alteration of organic acid content and *FUMARASE2 (FUM2*) expression in *nlp2* mutants. **(A)** Content in major TCA compounds of rosettes of WT, *nlp2-3*, *nlp7-1* and *nlp2-3 nlp7-1* mutants grown under non-limiting nitrate supply. **(B)** *FUM2*, encoding cytosolic fumerase2 involved in the massive accumulation of fumarate to sustain growth under high N supply, is specifically down-regulated in the rosette of *nlp2-3* and *nlp2-3 nlp7- 1* mutants grown under non-limiting nitrate supply.

**Supp. Movie 1**. Temporal dynamic of the nucleocytosolic shuttling of NLP2-GFP from 0 to 15 min after nitrate treatment. Movie was performed with confocal imaging on *nlp2-2* seedlings expressing *NLP2pro:NLP2-GFP*. N-starved seedlings were resupplied with nitrate during 15 min. Each second (one frame) is equivalent to one minute.

## Acknowledgements

We thank the entire NUTS team for enriching discussions and are very grateful to Sylvie Ferrario-Méry for excellent comments on earlier versions of this manuscript. We thank Philippe Marechal for plant care and our apprentices Cécile Larchévêgue and Kevan Barlafante for initial characterization of *nlp2* mutants. We also thank the NASC and ABRC for supplying Arabidopsis T-DNA insertion lines. This work has benefited from the support of IJPB’s Plant Observatory technological platforms and was supported by ANR IMANA (ANR-14-CE19-0008, MD, FR, AK) and ANR NITRASENSE (ANR-16-CE20-022, MD, ZK, AK). The IJPB and the IPS2 benefit from the support of the Saclay Plant Sciences (SPS) Network (ANR-17-EUR-0007).

## Author contributions

MD, FR and AK conceived the project. MD and VB produced all new plant material, MD, VB, and GD performed the RT-qPCR analyses. MD and VB performed growth analysis of adult plants, ALA, CPL and MD performed and analyzed RNA-seq, MD, JM and FR performed and analyzed ChIP-seq experiments, ZK performed BiFC, MD, ZK and AK performed confocal imaging, GC and RF performed mass spectrometry quantification of metabolites and GC, RF, JL analyses these data; MD and AK wrote the first version of the manuscript and all authors have been involved in producing the final version.

## References

Alonso-Blanco, C., Bentsink, L., Hanhart, C.J., Vries, H.B., and Koornneef, M. (2003). Analysis of Natural Allelic Variation at Seed Dormancy Loci of Arabidopsis thaliana. Genetics 164: 711–729.

Alvarez, J.M., Riveras, E., Vidal, E.A., Gras, D.E., Contreras-López, O., Tamayo, K.P., Aceituno, F., Gómez, I., Ruffel, S., Lejay, L., Jordana, X., and Gutiérrez, R.A. (2014). Systems approach identifies TGA1 and TGA4 transcription factors as important regulatory components of the nitrate response of Arabidopsis thaliana roots. Plant J 80: 1–13.

Alvarez, J.M., Schinke, A.-L., Brooks, M.D., Pasquino, A., Leonelli, L., Varala, K., Safi, A., Krouk, G., Krapp, A., and Coruzzi, G.M. (2020). Transient genome-wide interactions of the master transcription factor NLP7 initiate a rapid nitrogen-response cascade. Nat Commun 11: 1157.

Anoman, A.D., Muñoz-Bertomeu, J., Rosa-Téllez, S., Flores-Tornero, M., Serrano, R., Bueso, E., Fernie, A.R., Segura, J., and Ros, R. (2015). Plastidial Glycolytic Glyceraldehyde-3-Phosphate Dehydrogenase Is an Important Determinant in the Carbon and Nitrogen Metabolism of Heterotrophic Cells in Arabidopsis. Plant Physiol 169: 1619–1637.

Azimzadeh, J., Nacry, P., Christodoulidou, A., Drevensek, S., Camilleri, C., Amiour, N., Parcy, F., Pastuglia, M., and Bouchez, D. (2008). Arabidopsis TONNEAU1 Proteins Are Essential for Preprophase Band Formation and Interact with Centrin. Plant Cell 20: 2146–2159.

Bailey, T.L., Johnson, J., Grant, C.E., and Noble, W.S. (2015). The MEME Suite. Nucleic Acids Research 43: W39–W49.

Benjamini, Y. and Hochberg, Y. (1995). Controlling the False Discovery Rate: A Practical and Powerful Approach to Multiple Testing. Journal of the Royal Statistical Society. Series B (Methodological) 57: 289–300.

Bolger, A.M., Lohse, M., and Usadel, B. (2014). Trimmomatic: a flexible trimmer for Illumina sequence data. Bioinformatics 30: 2114–2120.

Bouyer, D., Heese, M., Chen, P., Harashima, H., Roudier, F., Grüttner, C., and Schnittger, A. (2018). Genome-wide identification of RETINOBLASTOMA RELATED 1 binding sites in Arabidopsis reveals novel DNA damage regulators. PLOS Genetics 14: e1007797.

Boyes, D.C., Zayed, A.M., Ascenzi, R., McCaskill, A.J., Hoffman, N.E., Davis, K.R., and Görlach, J. (2001). Growth stage-based phenotypic analysis of Arabidopsis: a model for high throughput functional genomics in plants. Plant Cell 13: 1499–1510.

Brooks, M.D., Cirrone, J., Pasquino, A.V., Alvarez, J.M., Swift, J., Mittal, S., Juang, C.-L., Varala, K., Gutiérrez, R.A., Krouk, G., Shasha, D., and Coruzzi, G.M. (2019). Network Walking charts transcriptional dynamics of nitrogen signaling by integrating validated and predicted genome-wide interactions. Nature Communications 10: 1569.

Canales, J., Moyano, T., Villarroel, E., and Gutiérrez, R. (2014). Systems analysis of transcriptome data provides new hypotheses about Arabidopsis root response to nitrate treatments. Frontiers in Plant Science 5.

Castaings, L., Camargo, A., Pocholle, D., Gaudon, V., Texier, Y., Boutet-Mercey, S., Taconnat, L., Renou, J.-P., Daniel-Vedele, F., Fernandez, E., Meyer, C., and Krapp, A. (2009). The nodule inception-like protein 7 modulates nitrate sensing and metabolism in Arabidopsis. The Plant Journal 57: 426–435.

Chardin, C., Girin, T., Roudier, F., Meyer, C., and Krapp, A. (2014). The plant RWP-RK transcription factors: key regulators of nitrogen responses and of gametophyte development. Journal of Experimental Botany 65: 5577–5587.

Chia, D.W., Yoder, T.J., Reiter, W.D., and Gibson, S.I. (2000). Fumaric acid: an overlooked form of fixed carbon in Arabidopsis and other plant species. Planta 211: 743–751.

Contreras-López, O., Vidal, E.A., Riveras, E., Alvarez, J.M., Moyano, T.C., Sparks, E.E., Medina, J., Pasquino, A., Benfey, P.N., Coruzzi, G.M., and Gutiérrez, R.A. (2022). Spatiotemporal analysis identifies ABF2 and ABF3 as key hubs of endodermal response to nitrate. PNAS 119.

Edgar, R., Domrachev, M., and Lash, A.E. (2002). Gene Expression Omnibus: NCBI gene expression and hybridization array data repository. Nucleic Acids Res 30: 207–210.

Estelle, M.A. and Somerville, C. (1987). Auxin-resistant mutants of Arabidopsis thaliana with an altered morphology. Mol Gen Genet 206: 200–206.

Fataftah, N., Mohr, C., Hajirezaei, M.-R., Wirén, N. von, and Humbeck, K. (2018). Changes in nitrogen availability lead to a reprogramming of pyruvate metabolism. BMC Plant Biol 18: 77.

Ferrario-Méry, S., Valadier, M.-H., and Foyer, C.H. (1998). Overexpression of Nitrate Reductase in Tobacco Delays Drought-Induced Decreases in Nitrate Reductase Activity and mRNA. Plant Physiol 117: 293–302.

Figueroa, C.M. and Lunn, J.E. (2016). A Tale of Two Sugars: Trehalose 6-Phosphate and Sucrose. Plant Physiol 172: 7–27.

Forzani, C., Duarte, G.T., Van Leene, J., Clément, G., Huguet, S., Paysant-Le- Roux, C., Mercier, R., De Jaeger, G., Leprince, A.-S., and Meyer, C. (2019). Mutations of the AtYAK1 Kinase Suppress TOR Deficiency in Arabidopsis. Cell Rep 27: 3696–3708.e5.

Gagnot, S., Tamby, J.-P., Martin-Magniette, M.-L., Bitton, F., Taconnat, L., Balzergue, S., Aubourg, S., Renou, J.-P., Lecharny, A., and Brunaud, V. (2008). CATdb: a public access to Arabidopsis transcriptome data from the URGV-CATMA platform. Nucleic Acids Res 36: D986–D990.

Gaudinier, A. et al. (2018). Transcriptional regulation of nitrogen-associated metabolism and growth. Nature 563: 259–264.

Gibon, Y., Pyl, E.-T., Sulpice, R., Lunn, J.E., Höhne, M., Günther, M., and Stitt, M. (2009). Adjustment of growth, starch turnover, protein content and central metabolism to a decrease of the carbon supply when Arabidopsis is grown in very short photoperiods. Plant, Cell & Environment 32: 859–874.

Gowri, G., Kenis, J.D., Ingemarsson, B., Redinbaugh, M.G., and Campbell, W.H. (1992). Nitrate reductase transcript is expressed in the primary response of maize to environmental nitrate. Plant Mol Biol 18: 55–64.

Guan, P., Ripoll, J.-J., Wang, R., Vuong, L., Bailey-Steinitz, L.J., Ye, D., and Crawford, N.M. (2017). Interacting TCP and NLP transcription factors control plant responses to nitrate availability. Proc Natl Acad Sci U S A 114: 2419– 2424.

Hachiya, T. and Sakakibara, H. (2017). Interactions between nitrate and ammonium in their uptake, allocation, assimilation, and signaling in plants. Journal of Experimental Botany 68: 2501–2512.

Hao, J., Pétriacq, P., de Bont, L., Hodges, M., and Gakière, B. (2018). Characterization of l-aspartate oxidase from Arabidopsis thaliana. Plant Sci 271: 133–142.

Hebbar, A., Moger, A., Hari, K., and Jolly, M.K. (2021). Interplay of Positive and Negative Feedback loops Governs Robustness in Multistable Biological Networks.: 2021.10.08.463488.

Heinz, S., Benner, C., Spann, N., Bertolino, E., Lin, Y.C., Laslo, P., Cheng, J.X., Murre, C., Singh, H., and Glass, C.K. (2010). Simple combinations of lineage-determining transcription factors prime cis-regulatory elements required for macrophage and B cell identities. Mol Cell 38: 576–589.

Hemming, M.L., Coy, S., Lin, J.-R., Andersen, J.L., Przybyl, J., Mazzola, E., Abdelhamid Ahmed, A.H., van de Rijn, M., Sorger, P.K., Armstrong, S.A., Demetri, G.D., and Santagata, S. (2021). HAND1 and BARX1 Act as Transcriptional and Anatomic Determinants of Malignancy in Gastrointestinal Stromal Tumor. Clin Cancer Res 27: 1706–1719.

Kiba, T. et al. (2018). Repression of Nitrogen Starvation Responses by Members of the Arabidopsis GARP-Type Transcription Factor NIGT1/HRS1 Subfamily. The Plant Cell 30: 925–945.

Konishi, M., Okitsu, T., and Yanagisawa, S. (2021). Nitrate-responsive NIN-like protein transcription factors perform unique and redundant roles in Arabidopsis. Journal of Experimental Botany 72: 5735–5750.

Konishi, M. and Yanagisawa, S. (2013). Arabidopsis NIN-like transcription factors have a central role in nitrate signalling. Nat Commun 4: 1617.

Konishi, M. and Yanagisawa, S. (2010). Identification of a nitrate-responsive cis-element in the Arabidopsis NIR1 promoter defines the presence of multiple cis-regulatory elements for nitrogen response. The Plant Journal 63: 269–282.

Konishi, M. and Yanagisawa, S. (2019). The role of protein-protein interactions mediated by the PB1 domain of NLP transcription factors in nitrate-inducible gene expression. BMC Plant Biol 19: 90.

Kopylova, E., Noé, L., and Touzet, H. (2012). SortMeRNA: fast and accurate filtering of ribosomal RNAs in metatranscriptomic data. Bioinformatics 28: 3211–3217.

Krapp, A., Berthomé, R., Orsel, M., Mercey-Boutet, S., Yu, A., Castaings, L., Elftieh, S., Major, H., Renou, J.-P., and Daniel-Vedele, F. (2011). Arabidopsis Roots and Shoots Show Distinct Temporal Adaptation Patterns toward Nitrogen Starvation. Plant Physiology 157: 1255–1282.

Krapp, A., David, L.C., Chardin, C., Girin, T., Marmagne, A., Leprince, A.-S., Chaillou, S., Ferrario-Méry, S., Meyer, C., and Daniel-Vedele, F. (2014). Nitrate transport and signalling in Arabidopsis. Journal of Experimental Botany 65: 789–798.

Kruger, N.J. and von Schaewen, A. (2003). The oxidative pentose phosphate pathway: structure and organisation. Curr Opin Plant Biol 6: 236–246.

Landt, S.G. et al. (2012). ChIP-seq guidelines and practices of the ENCODE and modENCODE consortia. Genome Res. 22: 1813–1831.

Langmead, B. and Salzberg, S.L. (2012). Fast gapped-read alignment with Bowtie 2. Nat Methods 9: 357–359.

Li, Q., Brown, J.B., Huang, H., and Bickel, P.J. (2011). MEASURING REPRODUCIBILITY OF HIGH-THROUGHPUT EXPERIMENTS. The Annals of Applied Statistics 5: 1752–1779.

Liu, J. and Bisseling, T. (2020). Evolution of NIN and NIN-like Genes in Relation to Nodule Symbiosis. Genes (Basel) 11: E777.

Liu, K. et al. (2017). Discovery of nitrate-CPK-NLP signalling in central nutrient-growth networks. Nature 545: 311–316.

Lunn, J.E., Feil, R., Hendriks, J.H., Gibon, Y., Morcuende, R., Osuna, D., Scheible, W.-R., Carillo, P., Hajirezaei, M.-R., and Stitt, M. (2006). Sugar-induced increases in trehalose 6-phosphate are correlated with redox activation of ADPglucose pyrophosphorylase and higher rates of starch synthesis in Arabidopsis thaliana. Biochemical Journal 397: 139–148.

Maeda, Y., Konishi, M., Kiba, T., Sakuraba, Y., Sawaki, N., Kurai, T., Ueda, Y., Sakakibara, H., and Yanagisawa, S. (2018). A NIGT1-centred transcriptional cascade regulates nitrate signalling and incorporates phosphorus starvation signals in Arabidopsis. Nat Commun 9: 1376.

Marchive, C., Roudier, F., Castaings, L., Bréhaut, V., Blondet, E., Colot, V., Meyer, C., and Krapp, A. (2013). Nuclear retention of the transcription factor NLP7 orchestrates the early response to nitrate in plants. Nature Communications 4: 1713.

Marschner, H. (1995). 8 - Functions of Mineral Nutrients: Macronutrients. In Mineral Nutrition of Higher Plants (Second Edition), H. Marschner, ed (Academic Press: London), pp. 229–312.

Masakapalli, S.K., Kruger, N.J., and Ratcliffe, R.G. (2013). The metabolic flux phenotype of heterotrophic Arabidopsis cells reveals a complex response to changes in nitrogen supply. The Plant Journal 74: 569–582.

Medici, A., Marshall-Colon, A., Ronzier, E., Szponarski, W., Wang, R., Gojon, A., Crawford, N.M., Ruffel, S., Coruzzi, G.M., and Krouk, G. (2015). AtNIGT1/HRS1 integrates nitrate and phosphate signals at the Arabidopsis root tip. Nature Communications 6: 6274.

Mu, X. and Luo, J. (2019). Evolutionary analyses of NIN-like proteins in plants and their roles in nitrate signaling. Cellular and Molecular Life Sciences 76.

Noctor, G. and Foyer, C.H. (1998). ASCORBATE AND GLUTATHIONE: Keeping Active Oxygen Under Control. Annu Rev Plant Physiol Plant Mol Biol 49: 249– 279.

O’Malley, R.C., Huang, S.C., Song, L., Lewsey, M.G., Bartlett, A., Nery, J.R., Galli, M., Gallavotti, A., and Ecker, J.R. (2016). Cistrome and Epicistrome Features Shape the Regulatory DNA Landscape. Cell 165: 1280–1292.

Orsel, M., Eulenburg, K., Krapp, A., and Daniel-Vedele, F. (2004). Disruption of the nitrate transporter genes AtNRT2.1 and AtNRT2.2 restricts growth at low external nitrate concentration. Planta 219: 714–721.

Osterwalder, M. et al. (2014). HAND2 Targets Define a Network of Transcriptional Regulators that Compartmentalize the Early Limb Bud Mesenchyme. Developmental Cell 31: 345–357.

Pracharoenwattana, I., Zhou, W., Keech, O., Francisco, P.B., Udomchalothorn, T., Tschoep, H., Stitt, M., Gibon, Y., and Smith, S.M. (2010). Arabidopsis has a cytosolic fumarase required for the massive allocation of photosynthate into fumaric acid and for rapid plant growth on high nitrogen. The Plant Journal 62: 785–795.

Quadrana, L., Bortolini Silveira, A., Mayhew, G.F., LeBlanc, C., Martienssen, R.A., Jeddeloh, J.A., and Colot, V. (2016). The Arabidopsis thaliana mobilome and its impact at the species level. eLife 5: e15716.

Rigaill, G. et al. (2018). Synthetic data sets for the identification of key ingredients for RNA-seq differential analysis. Brief Bioinform 19: 65–76.

Robinson, J.T., Thorvaldsdóttir, H., Winckler, W., Guttman, M., Lander, E.S., Getz, G., and Mesirov, J.P. (2011). Integrative genomics viewer. Nat Biotechnol 29: 24–26.

Robinson, M.D., McCarthy, D.J., and Smyth, G.K. (2010). edgeR: a Bioconductor package for differential expression analysis of digital gene expression data. Bioinformatics 26: 139–140.

Robinson, S.J. et al. (2009). An archived activation tagged population of Arabidopsis thalianato facilitate forward genetics approaches. BMC Plant Biology 9: 101.

Rubin, G., Tohge, T., Matsuda, F., Saito, K., and Scheible, W.-R. (2009). Members of the LBD Family of Transcription Factors Repress Anthocyanin Synthesis and Affect Additional Nitrogen Responses in Arabidopsis. Plant Cell 21: 3567–3584.

Safi, A. et al. (2021). GARP transcription factors repress Arabidopsis nitrogen starvation response via ROS-dependent and -independent pathways. Journal of Experimental Botany 72: 3881–3901.

Schauser, L., Wieloch, W., and Stougaard, J. (2005). Evolution of NIN-like proteins in Arabidopsis, rice, and Lotus japonicus. J Mol Evol 60: 229–237.

Scheible, W.-R., Morcuende, R., Czechowski, T., Fritz, C., Osuna, D., Palacios-Rojas, N., Schindelasch, D., Thimm, O., Udvardi, M.K., and Stitt, M. (2004). Genome-Wide Reprogramming of Primary and Secondary Metabolism, Protein Synthesis, Cellular Growth Processes, and the Regulatory Infrastructure of Arabidopsis in Response to Nitrogen. Plant Physiology 136: 2483–2499.

Shannon, P., Markiel, A., Ozier, O., Baliga, N.S., Wang, J.T., Ramage, D., Amin, N., Schwikowski, B., and Ideker, T. (2003). Cytoscape: a software environment for integrated models of biomolecular interaction networks. Genome Res 13: 2498–2504.

Song, L., Huang, S.-S.C., Wise, A., Castanon, R., Nery, J.R., Chen, H., Watanabe, M., Thomas, J., Bar-Joseph, Z., and Ecker, J.R. (2016). A transcription factor hierarchy defines an environmental stress response network. Science 354: aag1550.

Sultan, S.E. (2000). Phenotypic plasticity for plant development, function and life history. Trends in Plant Science 5: 537–542.

Tschoep, H., Gibon, Y., Carillo, P., Armengaud, P., Szecowka, M., Nunes-Nesi, A., Fernie, A.R., Koehl, K., and Stitt, M. (2009). Adjustment of growth and central metabolism to a mild but sustained nitrogen-limitation in Arabidopsis. Plant Cell Environ 32: 300–318.

Varala, K. et al. (2018). Temporal transcriptional logic of dynamic regulatory networks underlying nitrogen signaling and use in plants. PNAS 115: 6494–6499.

Vidal, E.A., Alvarez, J.M., Araus, V., Riveras, E., Brooks, M.D., Krouk, G., Ruffel, S., Lejay, L., Crawford, N.M., Coruzzi, G.M., and Gutiérrez, R.A. (2020). Nitrate in 2020: Thirty Years from Transport to Signaling Networks. The Plant Cell 32: 2094–2119.

Wang, R., Guegler, K., LaBrie, S.T., and Crawford, N.M. (2000). Genomic Analysis of a Nutrient Response in Arabidopsis Reveals Diverse Expression Patterns and Novel Metabolic and Potential Regulatory Genes Induced by Nitrate. The Plant Cell 12: 1491–1509.

Wang, R., Okamoto, M., Xing, X., and Crawford, N.M. (2003). Microarray Analysis of the Nitrate Response in Arabidopsis Roots and Shoots Reveals over 1,000 Rapidly Responding Genes and New Linkages to Glucose, Trehalose-6- Phosphate, Iron, and Sulfate Metabolism. Plant Physiology 132: 556–567.

Wilkinson, J.Q. and Crawford, N.M. (1993). Identification and characterization of a chlorate-resistant mutant of Arabidopsis thaliana with mutations in both nitrate reductase structural genes NIA1 and NIA2. Mol Gen Genet 239: 289–297.

Xu, G., Fan, X., and Miller, A. (2012). Plant Nitrogen Assimilation and Use Efficiency. Annual review of plant biology 63: 153–82.

Yan, D. et al. (2016). NIN-like protein 8 is a master regulator of nitrate-promoted seed germination in Arabidopsis. Nat Commun 7: 13179.

Zhang, Y., Sampathkumar, A., Kerber, S.M., Swart, C., Hill, C., Seerangan, K., Graf, A., Sweetlove, L., and Fernie, AR. (2021). A moonlighting role for enzymes of glycolysis in the co-localization of mitochondria and chloroplasts. Nat Commun 11:4509.

